# Helicase-mediated mechanism of SSU processome maturation and disassembly

**DOI:** 10.1101/2025.09.29.679232

**Authors:** Olga Buzovetsky, Sebastian Klinge

**Author notes:** Correspondence (S.K.).

## Abstract

Eukaryotic ribosomal small subunit (SSU) assembly requires the SSU processome, a nucleolar precursor containing the RNA chaperone U3 snoRNA. The underlying molecular mechanisms of SSU processome maturation, remodeling, disassembly, RNA quality control, and the transitions between states remain elusive due to a paucity of intermediates^1–3^. Here we report 16 native SSU processome structures alongside genetic data, revealing how two helicases, the Mtr4-exosome and Dhr1, are controlled for accurate and unidirectional ribosome biogenesis. Our data show how irreversible pre-ribosomal RNA degradation by the redundantly tethered RNA exosome couples the transformation of the SSU processome into a pre-40*S* particle during which Utp14 can probe evolving surfaces, ultimately positioning and activating Dhr1 to unwind the U3 snoRNA and initiate nucleolar pre-40*S* release. This study highlights a paradigm for large dynamic RNA-protein complexes where irreversible RNA degradation drives compositional changes and communicates these changes to govern enzyme activity while maintaining overall quality control.

## MAIN

The nucleolus, a membrane-less organelle, is generated via eukaryotic ribosome biogenesis in a transcription-dependent manner. The maturation of ribosomal small subunit (SSU) and large subunit (LSU) precursors generates distinct phases of this multi-layered organelle, which is maintained via multi-valent RNA-protein and protein-protein interactions. Early stages of eukaryotic ribosomal small subunit (SSU/40*S*) assembly involve the SSU processome, a giant nucleolar particle containing approximately a quarter of all eukaryotic ribosome assembly factors and three RNAs (SSU pre-rRNA/18*S*, a 5′ external transcribed spacer (5′ ETS) and the RNA chaperone U3 snoRNA)^1–3^. The 18*S* is composed of four domains (5′, central, 3′ major, and 3′ minor) which are housed in separate modules. By serving as the architectural blueprint for the SSU processome, the chaperone U3 snoRNA base-pairs with parts of the SSU pre-rRNA (via its box A and box A’ segments) as well as the 5′ ETS (via its 3’ and 5’ hinges). High-resolution structural studies of SSU processome assembly and maturation in yeast and human cells have started to reveal isolated native states before and after cleavage of the mature rRNA 5′ end (site A1) ^4–13^. After a poorly characterized departure of most assembly factors (SSU processome disassembly) a pre-40*S* particle emerges, which in addition to compacted rRNA domains still contains U3 snoRNA and the RNA helicase Dhr1^ref9^. The current absence of intermediate states highlights our limited understanding of the mechanisms that guide transitions between visualized states. The transition of the SSU processome from maturation to disassembly was suggested to involve the RNA exosome (with associated exonuclease Rrp6 and RNA helicase Mtr4)^10,12–15^. As the arbiter of RNA quality control, the RNA exosome ensures correct RNA processing with Rrp6 acting as an initial distributive RNA exonuclease before more processive RNA degradation by Rrp44. This system requires synergistic interactions between the RNA exosome and its substrates for RNA surveillance. Substrates that remain tethered to the RNA exosome are subject to RNA turnover whereas correctly processed RNA substrates are released^16^. The limited number of visualized SSU processome maturation and disassembly states has prevented a mechanistic understanding of their interaction with the RNA exosome and how RNA exosome function is coupled to SSU processome maturation and disassembly. Specifically, the mechanisms of RNA exosome recruitment, substrate access, and disengagement remain unclear^12,13^.

At the end of SSU processome disassembly, the DEAH-box helicase Dhr1 (DHX37 in humans) plays a key role in the removal of U3 snoRNA from a pre-40*S* particle to facilitate the formation of the central pseudoknot. However, despite ample studies^17–21^, the mechanism with which Dhr1 is initially recruited to the SSU processome, kept in an inactive conformation, subsequently repositioned to engage its substrate U3 snoRNA, and ultimately activated, remains unknown. Presently there is a lack of understanding of how the SSU processome, the RNA exosome, and Dhr1 work in concert to bring about unidirectional maturation and disassembly while maintaining RNA quality control throughout the entire process, which enables completed particles to leave from the nucleolus.

To address this, we determined native high-resolution structures of 16 *Saccharomyces cerevisiae* SSU processome states undergoing maturation and disassembly with resolutions up to 2.7 Ångstroms. Together with complementary genetics and an *in silico* protein-protein interaction screen, we have elucidated an RNA exosome-mediated mechanism that explains both SSU processome maturation and disassembly as a function of the 3′ to 5′ helicase activity of the RNA exosome-bound helicase Mtr4. Within this framework SSU processome disassembly, exosome-mediated RNA surveillance, and Dhr1 activation are tightly coordinated to maintain RNA quality control during the entire pathway.

To determine how Dhr1-bound SSU processomes undergo maturation and disassembly, we obtained native cryo-EM structures of particles containing Dhr1 during early, intermediate, and late stages (**Fig. 1 and Extended Data Fig. 1**). Early assembly states were selected using Dhr1 and the early assembly factor Kre33 as affinity baits (Dataset 1, **Supplementary Fig. 1**). For unbiased views of all Dhr1-bound states, Dhr1 was used as the sole affinity bait (Dataset 2, **Supplementary Fig. 2**). The late disassembly state containing activated Dhr1 (here State O) was obtained by selecting for particles containing Dhr1 and its activator Utp14 and lacking Utp7, which is present in earlier intermediates (Dataset 3, **Supplementary Fig. 3).**

**Fig. 1.**
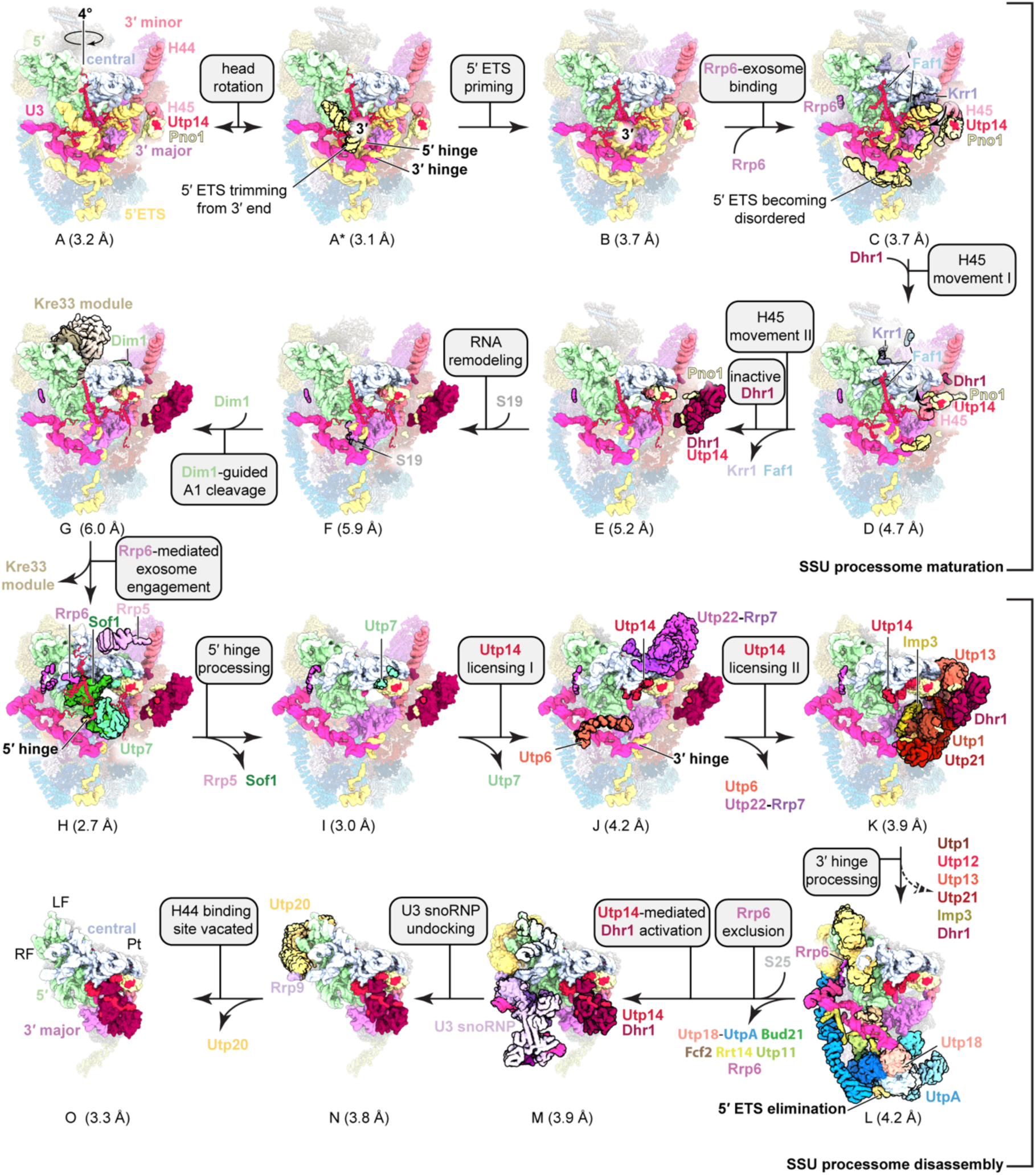
SSU processome maturation and disassembly pathway. Simplified atomic models of the SSU processome assembly pathway with color-coded rRNAs and ribosome assembly factors. Major changes in the rRNA between states are in the foreground and changes in protein compositions as well as functions are noted. Movements are indicated by arrows and protein or RNA elements that become ordered or depart between different assembly intermediates are highlighted with black outlines. The overall resolution of each cryo-EM reconstruction is noted below each state. A dashed arrow indicates tethering of protein components. Major structural features such as the left and right foot (LF and RF) along with the platform (Plat) are annotated in State O.

From a total of 213,766 cryo-EM images, initial data curation and comprehensive 3D classifications yielded 16 SSU processome reconstructions (states A-O) with overall resolutions of 2.7-6 Ångstroms (**Fig. 1 and Extended Fig. 1**). Focused refinements with 105 maps were used to obtain 16 composite maps for all structures (**Supplementary Figs. 4-25**), which enabled model building and assignment of 50 unique assembly factors, 21 ribosomal proteins, and three RNAs (5′ ETS, 18S rRNA, and U3 snoRNA) across all states (**Supplementary Tables 1 & 2**). Here, states are defined based on distinct rRNA folding transitions and the presence or absence of distinct assembly factors resulting in 15 unique states (States A-O), and a substate of State A that undergoes conformational changes only (State A*). Together these states provide unprecedented high-resolution coverage of native SSU processome maturation and disassembly that goes far beyond prior studies^9,10,12^ (**Extended Data Fig. 1-6, Supplementary Fig. 26 and 27**). High-resolution reconstructions further reveal the positions of post-translational modifications and their role in stabilizing distinct assembly intermediates (**Supplementary Fig. 28**)

### Chronology of SSU biogenesis

The large number of observed states allows us to visualize nucleolar SSU biogenesis as a function of RNA exosome-mediated 5′ ETS degradation, which first triggers cleavage of the 18*S* rRNA at site A1 (States A-G), and subsequently orchestrates SSU processome disassembly, rRNA compaction, and activation of Dhr1 (States H-O) (**Fig. 1, Extended Data Fig. 1 and 2**). The presence of the Rrp6 C-terminus in States C-L highlights the tethered presence of the RNA exosome, which is mediating transitions between states. This is contrary to prior studies which lacked the temporal resolution to precisely define both the chronology and causality of events leading to cleavage at site A1 and hence the underlying mechanism (**Supplementary Fig. 26**)^9^. Initial degradation of the 3′ end of the 5′ ETS on the exterior of the particle is followed by accommodation of the C-terminus of Rrp6 (State C), suggesting that the RNA exosome is responsible for this remodeling step. In State D the restructuring of the 5′ ETS in the interior of the SSU processome allows for partial release of Krr1 and Faf1 from their binding sites. This leads to the 3′ end of the 18*S* rRNA precursor (helix 45) being chaperoned by the assembly factor Pno1 towards the interior of the particle, thus allowing an N-terminal element of the DEAH-box helicase Dhr1 to bind. As the assembly factor Utp14 connects Pno1 with Dhr1 (**Supplementary Fig. 27 e and i**), the integration of helix 45 near the central domain results in the docking of the auto-inhibited Dhr1 in State E. Surprisingly we find that A1 cleavage in State G is solely triggered by the incorporation of Dim1. Here Dim1 probes the placement of helix 45 close to the central domain and stabilizes U3 snoRNA-bound regions of the 18S rRNA in an upward conformation. This facilitates Utp24-mediated A1 cleavage in particles that still contain the Kre33 module, which was previously thought to precede A1 cleavage^9^ (**Fig. 1, Extended Data Fig. 3**).

SSU processome disassembly further progresses as a function of 3′ to 5′ elimination of the 5′ ETS, resulting in compositional and conformational changes of the particle (**Fig.1**). During these compositional changes, Utp14, a largely unstructured protein containing several regulatory segments visualized in our structures (here referred to as a-f) can adopt different binding conformations as a response to SSU processome maturation and disassembly (**Extended Data Fig.6 a,d,e**).

With the processing of the 5′ hinge, distinct stages of disassembly are observed that involve the stepwise departure of assembly factors, such as Rrp5, Sof1, and Utp7 (States H-J). In States J and K, Utp14 licenses a new binding site for Dhr1, which was previously occupied by Utp7. The subsequent processing of the 3′ hinge (States K-L) releases much of the UtpB complex (except for Utp18) and the helicase Dhr1. In a major compositional remodeling step, the transition from State L to M marks the complete elimination of the 5′ ETS, which triggers the departure of the UtpA and UtpB complexes as well as individual assembly factors and Rrp6. This remodeling step subsequently allows Dhr1 to be repositioned and activated by Utp14 in a state-specific manner. States M-O therefore capture the activation of Dhr1 as further compaction of the particle triggers the undocking of U3 snoRNP proteins and Utp20. These results suggest a direct mechanism by which the RNA exosome could degrade the 5′ ETS to trigger large compositional and conformational changes that transform the SSU processome into a pre-40*S* particle.

### Integrated nucleolar RNA surveillance

Genetic studies suggested that the yeast RNA exosome binds the SSU processome via an interaction between the RNA helicase Mtr4 arch domain and the assembly factor Utp18 via an arch-interacting motif (AIM)^14^. While low-resolution density for the RNA exosome in the proximity of the SSU processome has been observed^10,12^, exactly how exosome recruitment and positioning is achieved across multiple states remains unclear. Moreover, the RNA exosome must efficiently detect and eliminate trapped SSU precursors within the nucleolus which have failed to complete biogenesis, yet how nucleolar RNA surveillance is set up to do so remains unresolved^10,12,14^. The 71 proteins present in the SSU processome together with the 14 proteins present within the RNA exosome provide a unique opportunity to screen pathway-wide protein-protein interactions *in silico* using Alphafold-Multimer^22–24^. We employed this approach to obtain an SSU processome predictome to test 3,570 unique binary interactions, which together with our high-resolution cryo-EM maps enabled the identification of many previously unknown interactions (**Fig. 2a, Extended Data Fig. 4, and Supplementary Table 3**). These data highlight key features of the SSU biogenesis pathway including the architectural roles of Sas10-domain containing complexes, tethering functions of Mpp10 for ribosomal protein S5 and Dim1, and predictions for the recruitment and activation of Dhr1/DHX37 (**Extended Data Fig. 4d,f**). In addition, mutually exclusive interactions between Kre33 and Faf1 or Bms1 are revealed, which now explain previously unassigned density for the human homologs of Kre33/Faf1 (NAT10/C1ORF13) in the human SSU processome^11^ (**Extended Data Fig. 4e**).

**Fig. 2.**
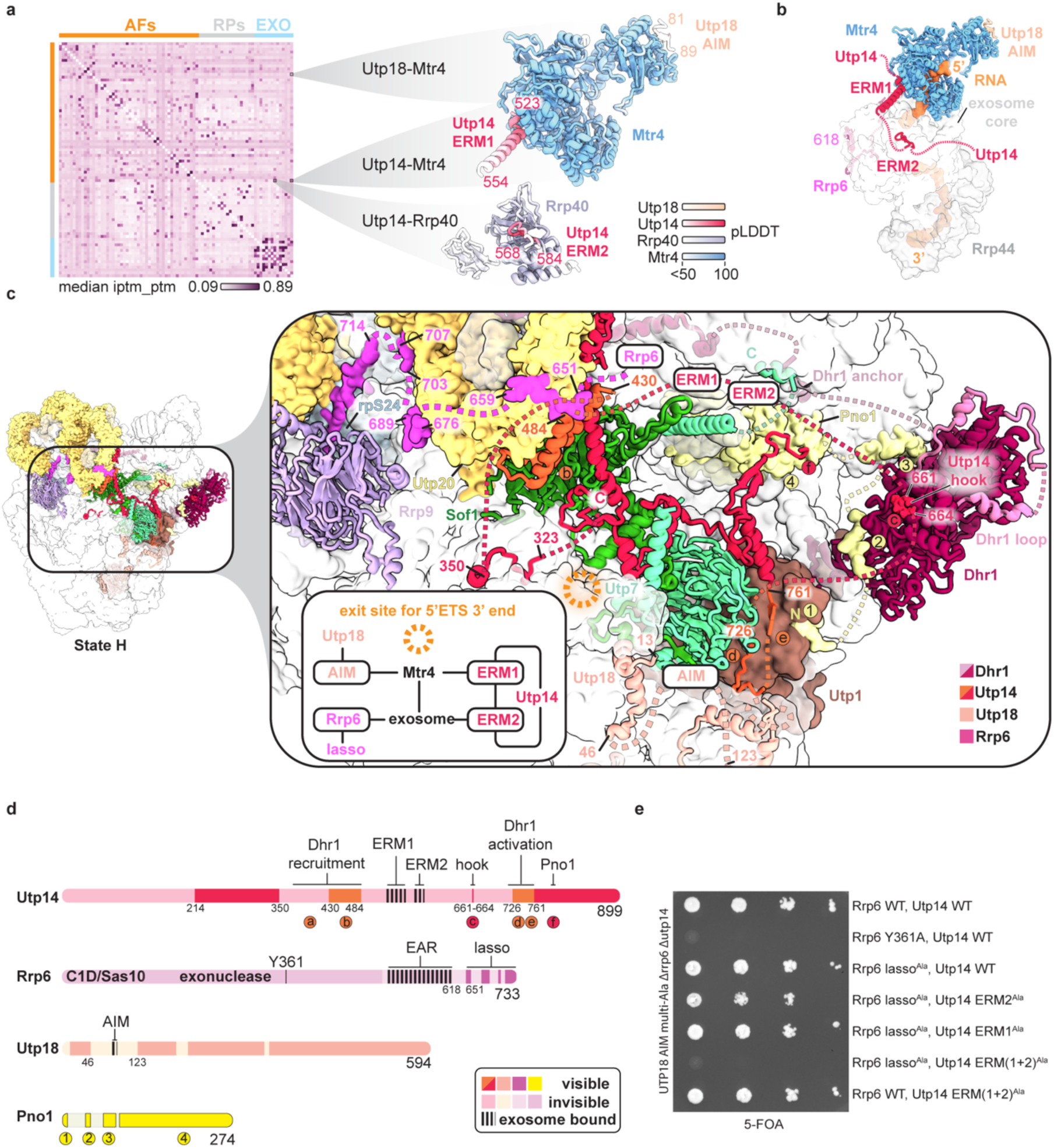
Interaction networks between SSU processome and the RNA exosome. **(a)** Yeast SSU processome predictome containing assembly factors (AFs), ribosomal proteins (RPs) and RNA exosome proteins (EXO) with color-coded confidence scores. Three AlphaFold-Multimer predictions are highlighted with color-coded proteins and confidence levels (pLDDT). **(b)** AlphaFold predictions for color-coded Utp14, Mtr4, and Utp18 AIM mapped onto a composite model of the RNA exosome with Rrp6 (PDBs 5vzj and 6d6q). **(c)** Cartoon representation of state H (left) and zoomed-in view (right) of the 5′ ETS 3′ end exit site (orange circles) with color-coded exosome interacting proteins. Regulatory elements of Utp14 are highlighted in orange. **(d)** Schematic of Utp14, Rrp6, Utp18, and Pno1 protein sequences in State H with interacting and functional elements labeled. **(e)** Growth analysis of Rrp6 and Utp14 mutants in a Utp18 multi-Ala background on 5-FOA plates.

A striking series of predicted interactions emerged from the SSU processome predictome that suggest extensive contacts between the RNA degradation machinery and the maturing and disassembling SSU processome **(Fig. 2a,b**). In addition to previously known interactions between the RNA helicase Mtr4 and Utp18^ref14^ and newly identified multivalent interactions between the yeast Rrp6 C-terminus (lasso) and the SSU processome in this study, the SSU processome predictome suggested that Utp14, the essential activator of Dhr1, additionally serves as a bridge between Mtr4 (via Exosome Recruitment Motif 1, ERM1), the core exosome component Rrp40 (via ERM2), and the SSU processome (**Fig. 2a,b**). This mode of binding is similar to the binding of the RNA exosome cofactor Mpp6^ref25^, with Mpp6 motifs having a high degree of sequence similarity to Utp14 ERM1 and ERM2 (**Extended Data Fig. 5a**).

Based on these predictions, a model emerges in which the RNA exosome contains four contact points for the SSU processome, which in addition to Utp18 AIM and the C-terminus of Rrp6 (lasso) include two peptides of Utp14 (ERM1 and ERM2) (**Fig. 2b**). Synergistically, the 2.7 Ångstrom structure of State H highlights that elements of Rrp6, Utp18 (AIM), and predicted Utp14 (ERM1 and ERM2) are all positioned in close proximity to the 3′ end of the 5′ ETS where the Mtr4 helicase needs to unwind the 5′ETS for both maturation and disassembly to occur (**Fig. 2c**). In State H, Utp14 occupies a strategic position with key regulatory elements that respond to conformational changes of the entire particle. Elements of Utp14 that will in later states provide a part of a Dhr1 binding site (segment b) or activate Dhr1 (segments d and e) are sequestered in State H near Sof1/Utp20 and Utp7/Utp1 respectively (**Fig. 2c,d).** Separately, the newly identified Utp14 hook (segment c) recruits the auto-inhibited Dhr1 and this inactive state is further stabilized by N-terminal elements of Pno1 (**Supplementary Fig. 27i,l**). This arrangement places ERM1 and ERM2 of Utp14 directly between segments b and c so that after cleavage at site A1 all elements that bridge the SSU processome and the RNA exosome are present near the 3′ end of the 5′ ETS (**Fig. 2c,d**).

The network consisting of Utp18, Rrp6 and Utp14 suggests that beyond exosome catalytic function, connective redundancy may play a major role in ensuring correct placement of the RNA exosome near the SSU processome. While prior data have indicated that Utp18 AIM mutants are synthetically lethal in the absence of Rrp6^ref12,14^, the relative contributions of Rrp6 exonuclease activity and its ability to bind to the SSU processome have remained unclear. Using yeast genetics we show that in the absence of the Utp18 AIM, an inactive Rrp6 exonuclease (Y361A)^26^ results in synthetic lethality, suggesting that catalytic activity of Rrp6 becomes essential if Mtr4 helicase cannot be correctly positioned (**Fig. 2e**). These data suggest that both positioning and the redundancy in connections of the RNA exosome to the SSU processome is essential for its function. Without the Utp18 AIM, three connections to the SSU processome remain via the Rrp6 lasso and the predicted Utp14 ERM1 & ERM2 (**Fig. 2a-d**). Our genetic data indicate that as long as any one of these connections is present, cell viability is maintained. By contrast, synthetic lethality is observed if all four connections are disrupted (**Fig. 2e**). Since the Rrp6 lasso is a multivalent binder of the SSU processome with four binding sites, we could separately show that iterative Rrp6 C-terminal truncations in a genetic background lacking the Utp18 and Utp14 connections gradually reduce growth (**Extended Data Fig. 5**). The SSU processome predictome together with structural data and genetics reveal a highly redundant nucleolar tethering network that couples SSU processome maturation, disassembly and RNA quality control. These findings are further consistent with the visualization of the human homolog of Rrp6 (EXOSC10) in the immediate vicinity of nucleolar SSU precursors^27^.

### Synchronized exosome-mediated disassembly

Elucidating the precise molecular mechanisms governing the controlled disassembly of the SSU processome remains a significant challenge, as current models lack the necessary sampling and resolution to mechanistically explain this process^9^. Here, States H-O capture a series of intermediates that follow RNA exosome-mediated disassembly (**Figs. 1 and 3**), allowing us to mechanistically explain how disassembly and RNA quality control are coupled. The irreversible 3′ to 5′ elimination of the 5′ ETS by the RNA exosome is directly coupled to the release of assembly factors in the exosome’s path. These include, among others, the Sof1-Utp7 complex at the 5′ hinge, the UtpB complex at the 3′ hinge, and finally Utp18 and the UtpA complex positioned at the start of the 5′ ETS (**Fig. 3a**). RNA exosome-mediated compositional changes to the SSU processome result in three key events that occur concurrently. First, the reduction of multi-valent RNA-protein and protein-protein interactions within the SSU processome leads to release of bound and tethered complexes. Second, the RNA exosome loses two of its interaction sites with the SSU processome and its function transitions from substrate engagement to substrate monitoring. Third, the changes of the SSU processome landscape provide new binding surfaces for Utp14, which organizes both Dhr1 repositioning and activation (**Fig. 3**).

**Fig. 3.**
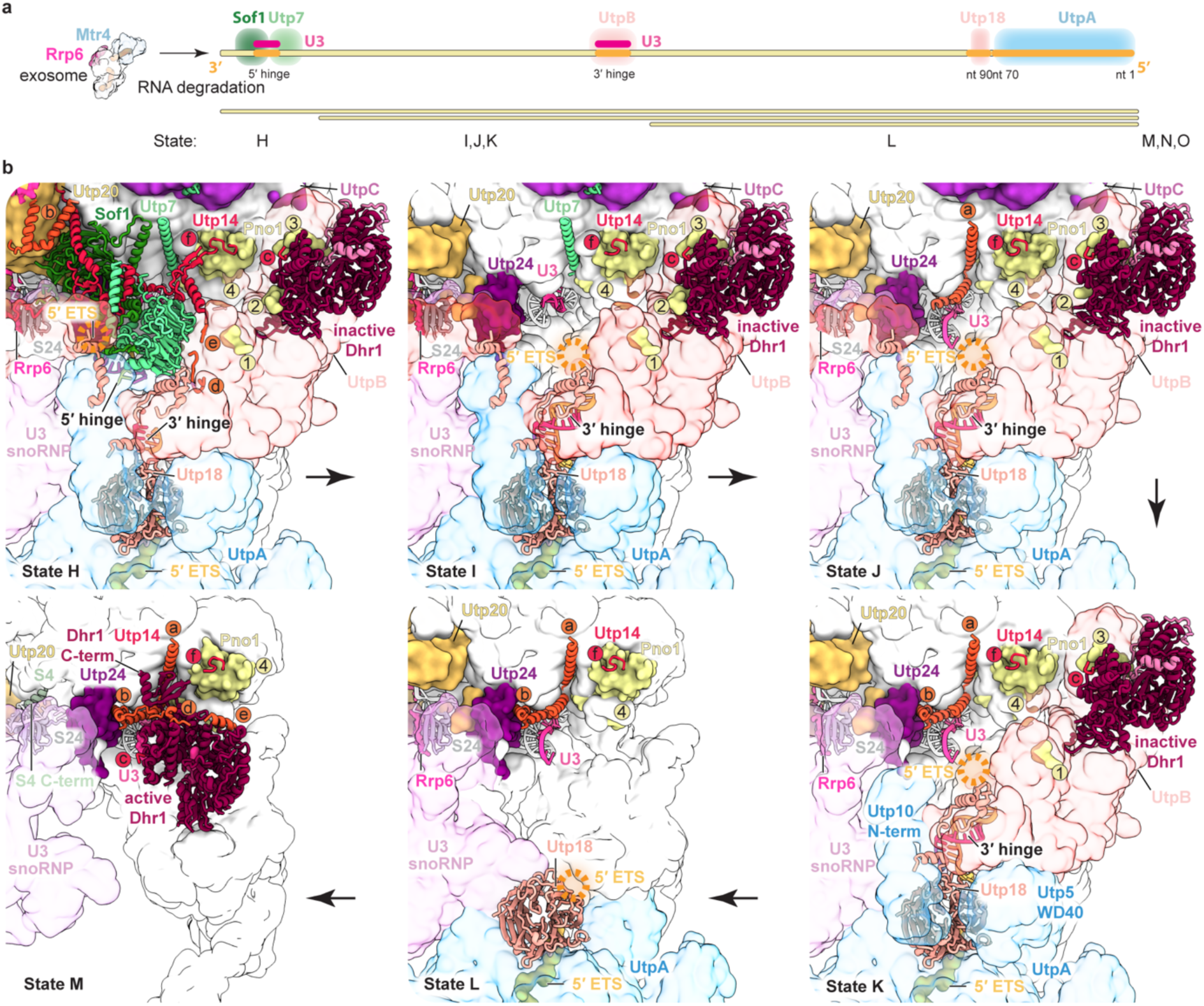
Helicase-mediated disassembly is coupled to RNA quality control. **(a)** Schematic of SSU processome disassembly as a function RNA degradation. RNA and protein elements bound to the 5′ ETS RNA are displayed as colored boxes and states with corresponding 5′ ETS length are listed below. **(b)** Architecture of the changes around the 5′ ETS exit sites (orange circles) from State H to M. Protein elements are color-coded and elements of Utp14 (b-f), Pno1 (1-4) and the state of Dhr1 are highlighted.

Beginning in State H, RNA exosome mediated processing of the 3′ end of the 5′ ETS alters the Sof1 binding site, weakening its interaction with Utp7 until only the C-terminus of Utp7 remains bound in State I. During this transition, key regulatory elements of Utp14 involved in subsequent Dhr1 recruitment (segment b) and activation (segments d and e) are released while remaining tethered via peptides bound to Dhr1 and Pno1 (segments c and f respectively). During subsequent 5′ ETS degradation, represented in states J and K, the Utp7 C-terminus is released, and its binding site is gradually occupied by elements of Utp14 (segments a and b) that prepare a binding site for Dhr1 as the UtpC complex is released. During the transition from State K to L, continued 5′ ETS processing results in UtpB losing its primary binding site at the 3′ hinge while remaining tethered to the SSU processome via Utp18, and Dhr1 remains connected to the particle via its N-terminus and the Utp14 hook (segment c) (**Fig. 3b**). Following complete 5′ ETS removal during the transition from State L to State M, UtpA, associated proteins, Utp18, and Rrp6 are fully released (**Fig. 3, Extended Data Fig. 6a**). During this key transition the roles of the RNA exosome and Dhr1 are dramatically changed.

The role of the RNA exosome shifts from the active substrate engaged state with four binding sites on the SSU processome to an overall surveillance state through the remaining two Utp14 ERM sites (**Extended Data Fig. 6a**). Furthermore, re-engagement of the substrate by the RNA exosome is prevented by the lack of a Utp18 binding site and the occlusion of the Rrp6 lasso binding site by the C-terminus of ribosomal protein S4 (**Fig. 3b, Extended Data Fig. 6b and c**). Separately, the role of Dhr1 changes from a tethered auto-inhibited state to a repositioned, substrate engaged, and activated enzyme poised to remove U3 snoRNA from the pre-40*S* particle.

### Molecular circuitry of Dhr1 activation

High-resolution crystal structures of Dhr1/DHX37 have shown that this DEAH-box helicase in isolation can exist in two states, an inactive state in which an auto-inhibitory loop prevents substrate binding, and an active state, in which single-stranded RNA acts as a substrate^11,19^. However, thus far no Utp14-mediated active state of Dhr1 has been observed. The comparison between the inactive state of Dhr1 in State H and the Utp14-mediated active state of Dhr1 in State O now rationalizes how irreversible RNA exosome-mediated 5′ ETS degradation orchestrates a cascade of events that triggers Dhr1 activation in a state-specific manner during SSU processome disassembly (**Fig. 4**).

**Fig. 4.**
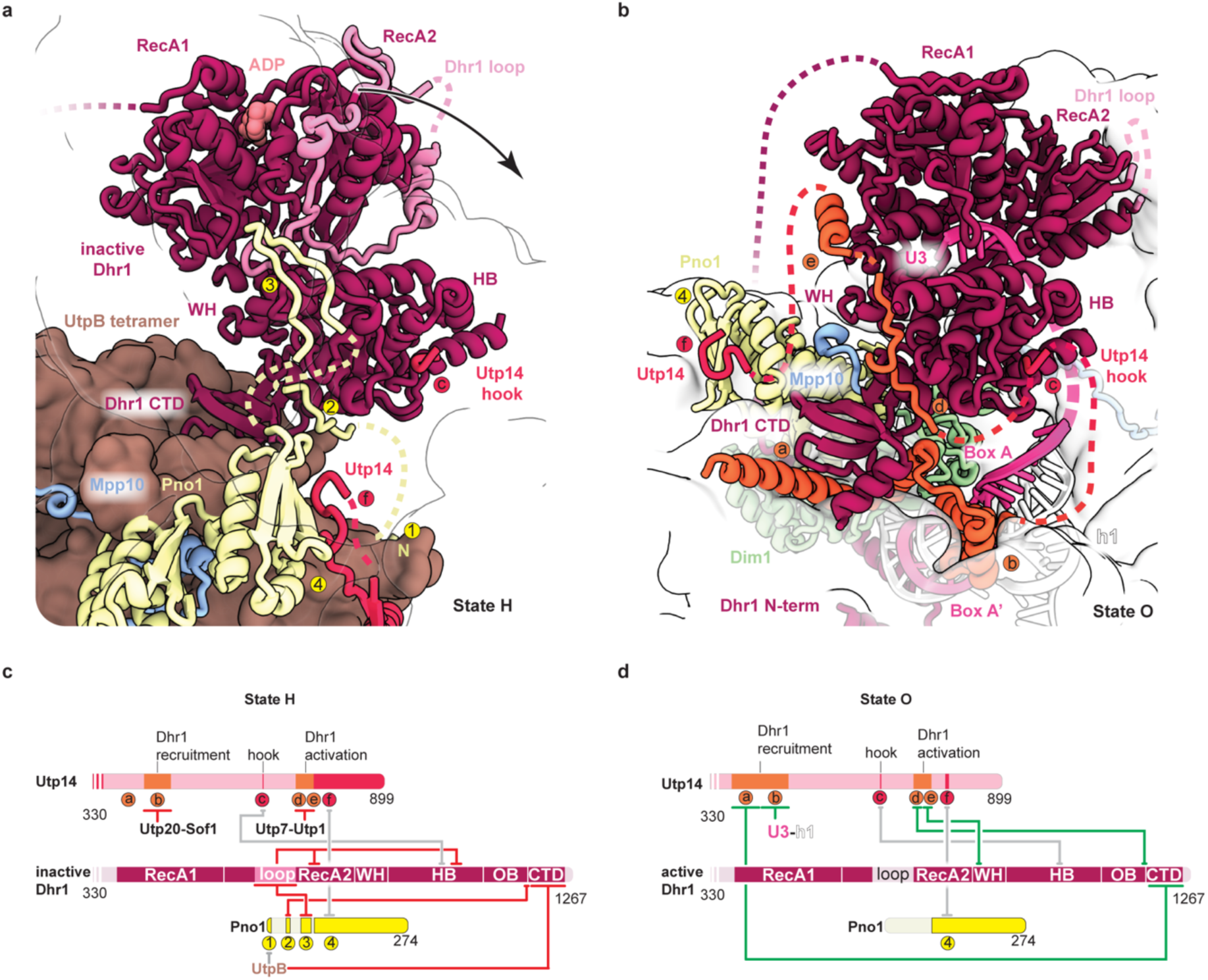
Mechanism of Utp14-mediated Dhr1 activation. **(a)** Zoomed-in view of inactive Dhr1 in State H, with auto-inhibitory loop (light pink) and inhibitory elements of Pno1 (1-4) and elements of Utp14 highlighted. The arrow indicates the conformational change of Dhr1 RecA domains. **(b)** Zoomed-in view of activated Dhr1 in State O. Activating Utp14 elements (a, b, d, and e) are highlighted in orange. **(c)** Schematic of State H molecular circuitry between Utp14, Dhr1 and Pno1. Interacting elements are colored grey and inhibitory interactions are indicated with red lines. **(d)** Schematic of State O molecular circuitry between Utp14, Dhr1 and Pno1. Interacting elements are colored grey and activating interactions are indicated with green lines.

In State H, the UtpB tetramer provides binding sites for both Dhr1 and Pno1 (elements 1 and 4), which enable additional elements of Pno1 (elements 2 and 3) to stabilize the auto-inhibited state of Dhr1. Element 3 of Pno1 stabilizes the auto-inhibitory loop of Dhr1, while element 2 of Pno1 prevents binding of segment d of Utp14, thus preventing a switch towards Dhr1 activation (**Fig. 4a**). Separately, segments of Utp14 that subsequently provide a binding site (segment b) or activate Dhr1 (segments d and e) are sequestered by Utp20/Sof1 and Utp7-Utp1 respectively (**Figs. 2c, 4a, c**).

In State O, the prior RNA exosome-mediated elimination of the UtpB complex, and the release of Utp14 segments a-e have dramatically changed the local environment for Dhr1. The new Dhr1 binding site (Utp14 segments a and b) is formed on top of its substrate U3 snoRNA, which still base-pairs with 18*S* elements that will form the central pseudoknot upon Dhr1-mediated U3 snoRNA release (**Fig. 4b**). The relative positions of Utp14 elements c and f now enable Utp14 elements d and e to stabilize a conformation of Dhr1 in which the RecA lobes are dramatically shifted to engage the substrate U3 snoRNA while displacing the auto-inhibitory loop (**Fig. 4b, d**). The architecture of the Utp14-mediated active form of Dhr1 in State O rationalizes a wealth of genetic and biochemical data from yeast and mammalian systems, highlighting that this mode of activation is distinct from G-patch mediated DEAH-box helicase activation (**Extended Data Fig. 6e-g**)^19,21^.

Thus, RNA exosome-mediated remodeling of the SSU processome triggers a series of compositional and conformational changes within the disassembling SSU processome that collectively switch Dhr1 from an “OFF” into an “ON” state that enables its engagement and processing of U3 snoRNA. The irreversible release of U3 snoRNA and the formation of the central pseudoknot prepare the resulting pre-40*S* particle for subsequent nuclear maturation where Utp14 is replaced by Nob1^ref2^, releasing the final link to nucleolar RNA surveillance.

### Conclusions

Nuclear bodies that contain biosynthetic processing pathways such as the nucleolus, Cajal bodies, and nuclear speckles share unified organizational principles by which RNA synthesis and processing results in the formation and dynamics of these multi-valent condensates^28,29^. For nucleolar SSU processome maturation and disassembly, we present a high-resolution temporal perspective, highlighting that this highly dynamic process is driven by the activities of two RNA helicases associated with the RNA degradation machinery (Mtr4) and the SSU processome itself (Dhr1) (**Figs. 1 and 2**). The irreversible unwinding and degradation of pre-ribosomal RNA (5′ ETS) changes the composition of disassembling SSU processomes, allowing regulatory peptides of Utp14 to probe the changing landscape of these particles to state-specifically reposition and activate Dhr1 for the terminal and irreversible release of U3 snoRNA (**Figs. 3 and 4**). The emerging model now highlights the importance of multivalent tethering at three levels. First, intra-SSU processome tethering enables pre-recruitment of assembly factors required for subsequent steps including restructuring of the SSU processome and A1 cleavage. Second, the extent (valency) of tethering between large particles such as the RNA exosome to the SSU processome changes as the exosome matures and disassembles the SSU processome, thereby shifting the role of the RNA exosome from initial recognition to substrate engagement and finally substrate surveillance. Third, at the level of the organelle, overall tethering of the maturing and disassembling SSU processomes to the nucleolus is decreased as more and more nucleolar assembly factors are released, thereby reducing the overall valency of particles before nucleolar release (**Fig. 5**). These data suggest that rather than the loss of non-specific interactions within a biomolecular condensate, it is the loss of specific protein-protein and protein-RNA interactions that enables the release of pre-ribosomal particles from the nucleolus. The scope of our study now allows us to illustrate molecular events at the Ångstrom scale that can rationalize the organizational principles at the micrometer scale as observed for the nucleolus^27^. This study highlights molecular transitions in large dynamic RNA-protein complexes driven by RNA helicases, and how these changes may be coupled to movement through biomolecular condensates in nuclear bodies.

**Fig. 5.**
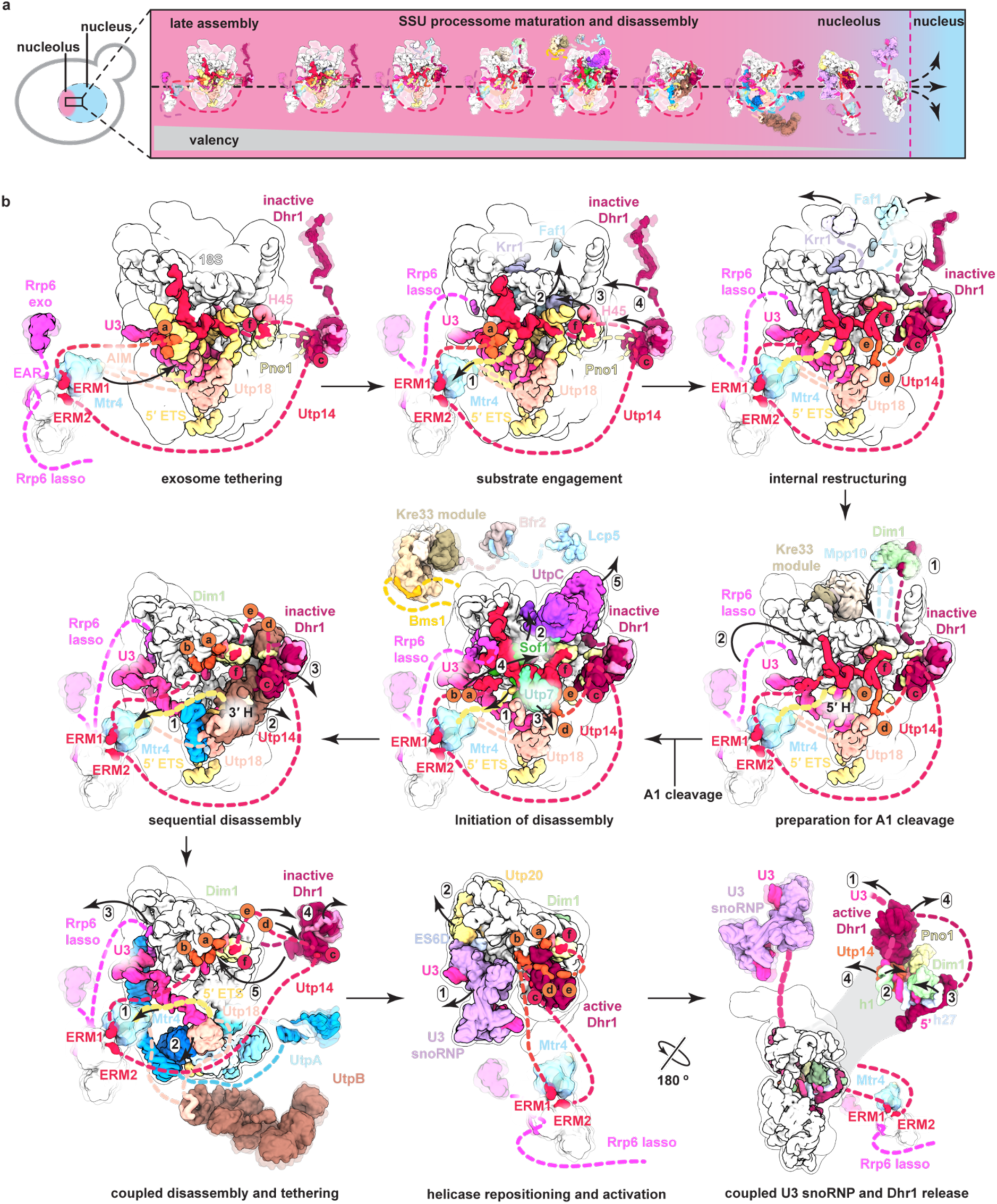
Model of the mechanism of SSU biogenesis in the nucleolus. **(a)** Schematic model of yeast SSU biogenesis in the nucleolus. Decreasing valency of maturing particles is indicated below. **(b)** SSU processome maturation and disassembly pathway with key transitions are indicated. Color-coded tethered proteins and complexes are indicated with linkers. Arrows and numbers highlight the order of transition steps (RNA degradation, conformational changes, protein release, or protein repositioning to new binding sites) that lead to particles with low valency during disassembly.

## METHODS

### Generation of endogenously tagged DHR1, KRE33, UTP7 and UTP14 strains

Assembly intermediates of the *S. cerevisiae* SSU processomes were purified from three genetically modified BY4741 (MATa his3Δ leu2Δ0 met15Δ0 ura3Δ0) strains. Where genes of interest (DHR1, KRE33, UTP7 and UTP14) were endogenously tagged: the first strain containing C-terminal tandem 3C protease-cleavable mCherry and TEV-cleavable alfa peptide tag on DHR1 along with a 3C cleavable C-terminal GFP-tag on KRE33 (MATa his3Δ leu2Δ0 met15Δ0 ura3Δ0 DHR1-linker-tev-alfa tag-3c-mCherry under Hygromycin B selection and KRE33-linker-tev-GFP under Nourseothricin selection), the second strain containing only C-terminal tandem 3C protease-cleavable mCherry and TEV-cleavable alfa peptide tag on DHR1 (MATa his3Δ leu2Δ0 met15Δ0 ura3Δ0 DHR1-linker-tev-alfa tag-3c-mCherry under Hygromycin B selection) and the third strain containing C-terminal tandem 3C protease-cleavable mCherry and a TEV-cleavable alfa peptide tag on DHR1 along with TEV cleavable C-terminal GFP-tag on UTP7 and C-terminally streptavidin-binding-peptide (sbp) tag on UTP14 (MATa his3Δ leu2Δ0 met15Δ0 ura3Δ0 DHR1-linker-tev-alfa tag-3c-mCherry under Hygromycin B selection and UTP7-linker-tev-GFP under G418 selection and UTP14-linker-SBP under Nourseothricin selection). The three strains were generated with standard genomic tagging techniques using the following primers for yeast genomic tagging:

DHR1 forward primer:

TCACCAGAAAGGGCTTCCAGACCATCACAGGTGAAGAGAAAGAAAAAAAAACGCTGCAGGTCGACGGATCC

DHR1 reverse primer:

TTAAGTGGTTGCAATTATTTGATGCCCTAGATAGGAATAATGTATTCTTCGGCAGATCCGCGGCCGCATAGG

Kre33 forward primer:

AAGAGATGAAAGCTATGAAAAAACCAAGAAAGTCTAAAAAGGCTGCAAATACGCTGCAGGTCGACGGATCC

KRE33 reverse primer :

TGTAAAGGTTCAAACATCAACTATGTTTCTATTCTATATTATTGTACAAAGGCAGATCCGCGGCCGCATAGG

UTP7 forward primer:

TTTCAGAAGACCACAAGGATGTCATCGAAGAGGCATTGAGCAGATTCGGCACGCTGCAGGTCGACGGATCC

UTP7 reverse primer:

GCTATATTAATATGCAATCGATTCTCATACTGTCAACTTTTTGAACATGAGGCAGATCCGCGGCCGCATAGG

UTP14 forward primer:

TCATGACTAAGCCAGGCCAAGTTATTGATCCTTTGAAGGCACCATTTAAGACGCTGCAGGTCGACGGATCC

UTP14 reverse primer:

ATTATTCCAGTATTACTATTCTACACAATGCATAATAAATAGATATAAAAGGCAGATCCGCGGCCGCATAGG

The three strains were grown at 30 °C in YPD media (1% yeast extract, 2% peptone and 2% glucose) to optical density of OD 1 measured at 600 nm. Cells were harvested at 4,000*g* for 5 min, washed once with 1 l of ice cold ddH2O and once with a volume of ddH2O supplemented with protease inhibitors (E64, pepstatin and PMSF) equal to the mass of the cell pellet. The final pellets of ∼30-40g were flash frozen in liquid nitrogen. Pellets were lysed by cryo-grinding using a Retsch Planetary Ball Mill PM100 and powder was stored at –80 °C.

### Purification of SSU processome intermediates

Cryo-ground powder was resuspended with buffer 1 (60mM Tris pH7.5, 50mM NaCl, 2mM MgCl2, 5% glycerol, 0.1% NP-40) with addition of PMSF, pepstatin and E64 protease inhibitors. The suspension was cleared by centrifugation at 4 °C and 40,000*g* for 20 min and lysate was incubated with 800 µl of packed NHS–sepharose beads (Cytiva) coupled to anti-mCherry nanobodies for the first capture for all purifications and incubated for 3.5 h at 4 °C on a nutator. Beads were pelleted by centrifugation at 4 °C for 1 min at 127*g*. After five washes in buffer 1, complexes were eluted in buffer EB (60mM Tris, 50mM NaCl, 5mM MgCl2, 2% glycerol, 0.01% NP-40) supplemented with 3C protease for 1 h at 4 °C. Beads were pelleted by centrifugation at 4 °C for 1 min at 127*g* and supernatant was eluted and incubated with 80 µl of packed NHS–sepharose beads (Cytiva) coupled to anti-GFP nanobodies for second capture (Dhr1/Kre33 and Dhr1/Utp14 purifications) while anti-alfa nanobodies were used for a second Dhr1 (Dhr1 only). Beads were incubated for 1 h at 4 °C on a nutator and pelleted by centrifugation at 4 °C for 1 min at 127*g* and washed twice with 2 ml of Buffer EB. After the second wash, for Dhr1/Kre33 and Dhr1 only purifications, beads were pelleted and resuspended with 25 µl of Buffer EB supplemented with 1mM DTT and TEV protease and incubated on ice for 1 h. Beads were pelleted by centrifugation at 4 °C for 10 min at 21,130g and the supernatant was collected. For Dhr1/Utp14 purification, after the second capture (here Utp7), beads were pelleted by centrifugation at 4 °C for 1 min at 127*g* and flowthrough was collected, the rest of the beads were discarded (this step was used to remove earlier contaminating complexes). The flowthrough was incubated with 40 µl of packed NHS–sepharose beads (Cytiva) coupled to streptavidin for 1 h at 4 °C on a nutator. Beads were pelleted by centrifugation at 4 °C for 1 min at 127*g* and washed twice with 2 ml of Buffer EB. After the second wash, beads were pelleted and resuspended with 40 µl of Buffer EB supplemented with 1mM DTT and 5 mM of D-biotin (Amresco) and incubated on ice for 20 min. Beads were pelleted by centrifugation at 4 °C for 10 min at 21,130g and the supernatant was collected.

### Cryo-EM grid preparation and data acquisition

Cryo-EM grids were prepared using a Vitrobot Mark IV robot (FEI Company) set to 90% humidity and 18 °C temperature. Three and a half microliters of the eluted solution was applied to a glow-discharged Quantifoil Au R3.5/1 with a layer of 2-nm ultrathin carbon (LFH7100AR35, Electron Microscopy Sciences). After 2.5 min incubation inside the Vitrobot chamber, the excess solution was manually blotted and a fresh sample of 3.5 µl was reapplied inside the Vitrobot chamber and incubated for another 2.5 min. The lower the concentration of particles of a given preparation, the higher the number of applications that were done on each grid. For the Dhr1 and Kre33 dataset, a total of five applications were done on each grid. The grid was then blotted (blot force of 8 and blot time of 9 s) and plunged into liquid ethane. For the Dhr1dataset, a total of two applications were done for each grid. The grid was then blotted (blot force of 8 and blot time of 7 s) and plunged into liquid ethane and for the Dhr1 and Utp14 dataset, a total of five applications were done on each grid to achieve a good distribution of particles. The grid was then blotted (blot force of 8 and blot time of 9 s) and plunged into liquid ethane. Grids were imaged on a Titan Krios electron microscope (FEI) with an energy filter (slit width of 20 eV) and a K3 Summit detector (Gatan) operating at 300 kV with a nominal magnification of ×64,000.

SerialEM^30^ was used to collect four datasets. 44,272 micrographs were collected for the Kre33/Dhr1 dataset, two datasets of Dhr1 were collected totaling to 84,463 micrographs and 85,031 micrographs were collected for Dhr1 and Utp14 dataset. All datasets were collected with a defocus range of −1 to −2.5 µm and a super-resolution pixel size of 0.54 Å. Micrographs contained 40 frames using a total dose of 25.3-30.8 e−per pixel per second (specimen pixel size of 1.08 Å per pixel) with an exposure time of 2-2.5 s and a total dose of 61.7 -63.1e− Å−2. A multi-shot strategy was used to record nine micrographs per hole at each stage position with the same defocus range, electron dose and frame count.

### Cryo-EM data processing Dhr1 and Kre33 dataset

The Dhr1 and Kre33 dataset was processed using a combination of RELION 5beta^31^ and cryoSparc v4.6^ref32^. 44,272 movies were gain corrected, dose weighted, aligned, with each dataset having different optic groups and binned to a pixel size of 1.08 Å using RELION’s implementation of a MotionCor2-like algorithm^33^. Micrograph defocus was estimated using Gctf^34^. Particles were picked using Laplacian (autopick) in RELION 5beta and post 2D classification a total of 2,817,609 particles were re-extracted at a pixel size of 4.32 Å per pixel (4× binning) and underwent 3D classification in RELION 5beta with alignment using a reference map from previous datasets. Two good classes were combined, and duplicates were removed to result in 308,351 total particles. These particles were subjected to three rounds of CTF refinement and Bayesian polishing in RELION 5beta. Post polishing all homogenous refinements were completed in CryoSparc v4.6 and classifications were all done in RELION 5beta, particle positions from cryosparc are converted into RELION using pyem software csparc2star.py^35^. The polished particles were subjected to a homogenous refinements resulting in a reconstruction at a global resolution of 3.3 Å. To separate the states present in the consensus reconstruction, multiple 3D classifications without alignment were performed. This was followed by 3D variability^36^ in CryoSparc v4.6 for analysis of the type of heterogeneity present in the data which guided the creation of a mask around the region of interest and further 3D classification without alignment on the region of variability. Eight distinct states (sates A-G) were isolated from the dataset that showed unique features in the progression of maturation of the SSU processome pathway (**Supplementary** Fig.1). The global resolution of the eight states ranged from 3-5.9 Å.

### Cryo-EM data processing Dhr1 dataset

Dhr1 dataset was processed using a combination of RELION 5beta^31^ and cryoSparc v4.6^ref32^. 84,463 movies were gain corrected, dose weighted, aligned, with each dataset having different optic groups and binned to a pixel size of 1.08 Å using RELION’s implementation of a MotionCor2-like algorithm^33^. Micrograph defocus was estimated using Gctf^34^. Particles were picked using crYOLO 1.7.5^37^ and post 2D classification a total of 13,198,550 particles were re-extracted at a pixel size of 4.32 Å per pixel (4× binning) and underwent heterogenous refinement in CryoSparc v4.6 using a reference map from previous classification in RELION 5beta. One good class was isolated resulting in 1,933,969 total particles. These particles were subjected to three rounds of CTF refinement and Bayesian polishing in RELION 5beta. Post polishing all homogenous refinements were completed in CryoSparc v4.6 and classifications were all done in RELION 5beta. Particle positions from cryosparc are converted into RELION using pyem software csparc2star.py^35^. The polished particles were subjected to a homogenous refinement resulting in a reconstruction at a global resolution of 3 Å. To separate the states present in the consensus reconstruction, multiple iterations of 3D classifications without alignment was performed. Followed by 3D variability^36^ in CryoSparc v4.6 for analysis of the type of heterogeneity present in the data which guided the creation of a mask around the region of interest and further 3D classification without alignment on the region of variability. Seven distinct states (sates H-N) were isolated from the dataset that showed unique features in the progression of the disassembly of the SSU processome pathway (**Supplementary Fig.2**). The global resolution of the seven states ranged from 2.65-4.3 Å.

### Cryo-EM data processing Dhr1 and Utp14 dataset

The Dhr1 and Utp14 dataset was processed using a combination of RELION 5beta^31^ and cryoSparc v4.6^ref32^. 85,031 movies were gain corrected, dose weighted, aligned, with each dataset having different optic groups and binned to a pixel size of 1.08 Å using CryoSparc v 4.6 motion correction. Micrograph defocus was estimated using Patch CTF and particles were picked using a template picker resulting in a total of 27,069,273 particles which underwent heterogenous refinement and subsequent global and local CTF refinements, followed by reference motion correction. Post polishing all homogenous refinements were completed in CryoSparc v4.6 and classifications were all done in RELION 5beta. Particle positions from cryosparc are converted into RELION using pyem software csparc2star.py^35^. The polished particles were subjected to a homogenous refinement resulting in a reconstruction at a global resolution of 2.9 Å. To separate the states present in the consensus reconstruction, multiple iterations of 3D classifications without alignment were performed. Followed by 3D variability^36^ in CryoSparc v4.6 for analysis of the type of heterogeneity present in the data which guided the creation of a mask around the region of interest and further 3D classification without alignment on the region of variability. State O was isolated from the dataset with global resolution of 3.25 Å that showed clear density for Utp14-bound Dhr1 (**Supplementary Fig. 3**).

### Generation of focused and composite maps for model buildings

Composite maps for the total of 16 states were generated from combined focused maps to facilitate model building. Focused maps were generated using subtraction and refinement masks generated in CryoSparc v4.6. Each focused map was made by particle subtraction with a masked region followed by masked local refinement. Local resolution estimation for overall and all focused maps and filtering of the overall map were performed using CryoSparc v4.6. Focused maps were combined into a composite map using the ‘vop max’ command in ChimeraX^38^ (**Supplementary Figs. 4-25**).

### Model building and refinement

A combination of AlphaFold structure predictions^39^, existing X-ray/EM structures, and *de novo* model building was used to build the 16 SSU processome assembly intermediates. A starting model (PDB: 5WLC/6KE6) that included all ribosomal proteins and RNA was used as initial template for rigid body docking into the State H composite map since it is of highest resolution. All template ribosomal protein models were manually adjusted using COOT^40^. State H was then used as a template to build the proteins and RNA into the other 15 states with manual adjustments in COOT. The final models for the 16 states were real-space refined with three cycles of refinement in PHENIX using phenix.real_space_refine^41^ using secondary structure restraints for proteins and RNA. In regions with medium to low resolution, protein sidechains were trimmed to the Cβ position after all-atom refinement. The final model refinement statistics can be found in **Supplementary Table 1**. The maps and models were analyzed and visualized in ChimeraX^38^.

### SSU processome predictome

Proteins present within States A-O together with 14 exosome proteins were screened for binary interactions. The resulting 3570 unique interactions were screened using the default settings in the AlphaPulldown^23^ implementation of Alphafold-Multimer^22^. For the SSU processome predictome (**Fig. 2a**) the median iptm_ptm score of five models was plotted (**Supplementary Table 3**).

### Yeast Growth Assays

The *rrp6Δ utp14 Δ* strain and *rrp6Δ utp18 Δ utp14 Δ* strain in the BY4741 background were used for all studies. These strains were transformed with two yeast centromeric vectors, one vector under URA3 selection bearing wild-type RRP6 and UTP14 derived from pRSII416 and second vector under LEU2 selection bearing wild-type RRP6 or rrp6 alleles containing mutations or c-terminal truncations in conjunction with UTP14 wild-type or utp14 alleles containing mutations derived from pRSII415. Strains carrying both pRSII415 and pRSII416 plasmids were selected on minimal media (SD-Ura and Leu) after transformation. Colonies grown on the selection plates were selected and grown in minimal media (SD-Ura and Leu) liquid cultures. Loss of the *URA3* plasmid was done on minimal media agar plates (SD–Leu + 5-FOA) plates by spotting serial 10-fold dilutions (starting at OD600 of 1) of liquid cultures. Growth was monitored at 30°C over a period of 5 days.

## Data availability

Raw unaligned multi-frame movies and aligned micrographs have been deposited in the Electron Microscopy Public Image Archive (EMPIAR). The cryo-EM maps and atomic models have been deposited in the Electron Microscopy Data Bank (EMDB) and the Protein Data Bank (PDB): State A (EMD-49075, PDB 9N6V), State A* (EMD-49076, PDB 9N6W), State B (EMD-49077, PDB 9N6X), State C (EMD-49078, PDB 9N6Y), Sate D (EMD-49079, PDB 9N6Z), Sate E (EMD-49080, PDB 9N70), State F (EMD-49082, PDB 9N72), State G (EMD-49083, PDB 9N73), State H (EMD-49084, PDB 9N74), State I (EMD-49085, PDB 9N75), State J (EMD-49086, PDB 9N76), State K (EMD-49087, PDB 9N77), State L (EMD-49088, PDB 9N78), State M (EMD-49089, PDB 9N79), State N (EMD-49090, PDB 9N7A), State O (EMD-49091, PDB 9N7B).

## Acknowledgements

We would like to thank M. Ebrahim, J. Sotiris, and H. Ng at the Evelyn Gruss Lipper Cryo-Electron Microscopy Resource Center at The Rockefeller University for assistance with grid screening and data collection. Mass spectrometry data was generated by the Proteomics Resource Center at The Rockefeller University (RRID:SCR_017797) using instrumentation funded by the Sohn Conferences Foundation and the Loena M. and Harry B. Helmsley Charitable Trust, and we particularly thank H. Molina, S. Heissel, and C. Peralta for assistance. We thank A. Vanden Broeck for advice with cryo-EM data processing and scripts for figures. We also thank K. Ye for sharing experimental data from a prior study^10^ and members of the Klinge laboratory for critical reading of this manuscript.

## Author Contributions

S.K. conceived the study. O.B. and S.K. designed the experiments and analyzed data. O.B. performed all experiments, including strain generation, complex purification, cryo-EM data processing, model building, and yeast genetics studies. All authors wrote and edited the manuscript.

## Author information

Laboratory of Protein and Nucleic Acid Chemistry, The Rockefeller University, New York, NY, USA

Sebastian Klinge, Olga Buzovetsky

## Ethics declarations

The authors declare no competing interests.

## EXTENDED DATA FIGURE LEGENDS

**Extended Data Fig. 1.**
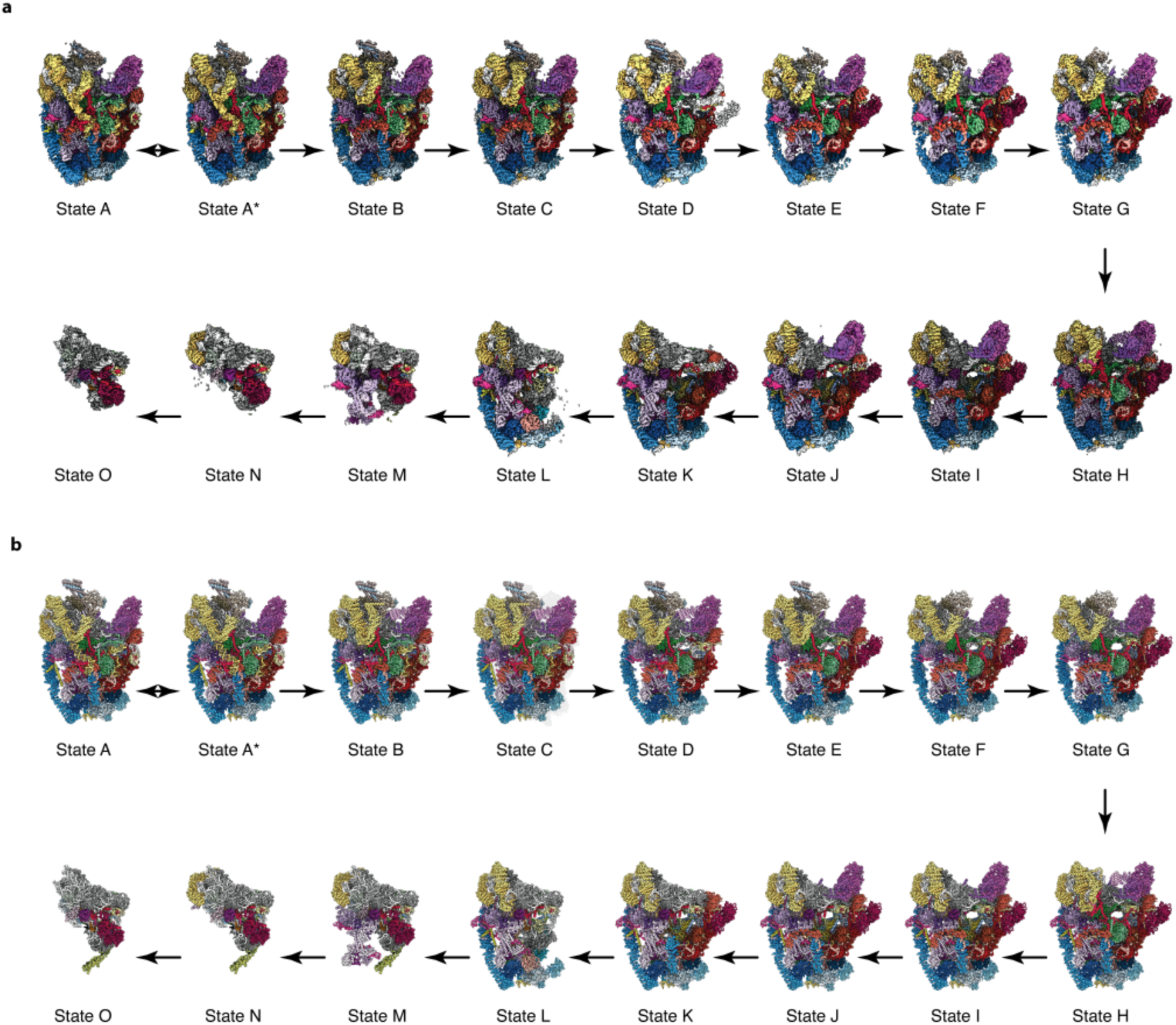
Cryo-EM maps and models of the SSU processomes. **(a)** Composite cryo-EM maps and **(b)** atomic models colored by protein components highlighting stepwise maturation of the SSU processome.

**Extended Data Fig. 2.**
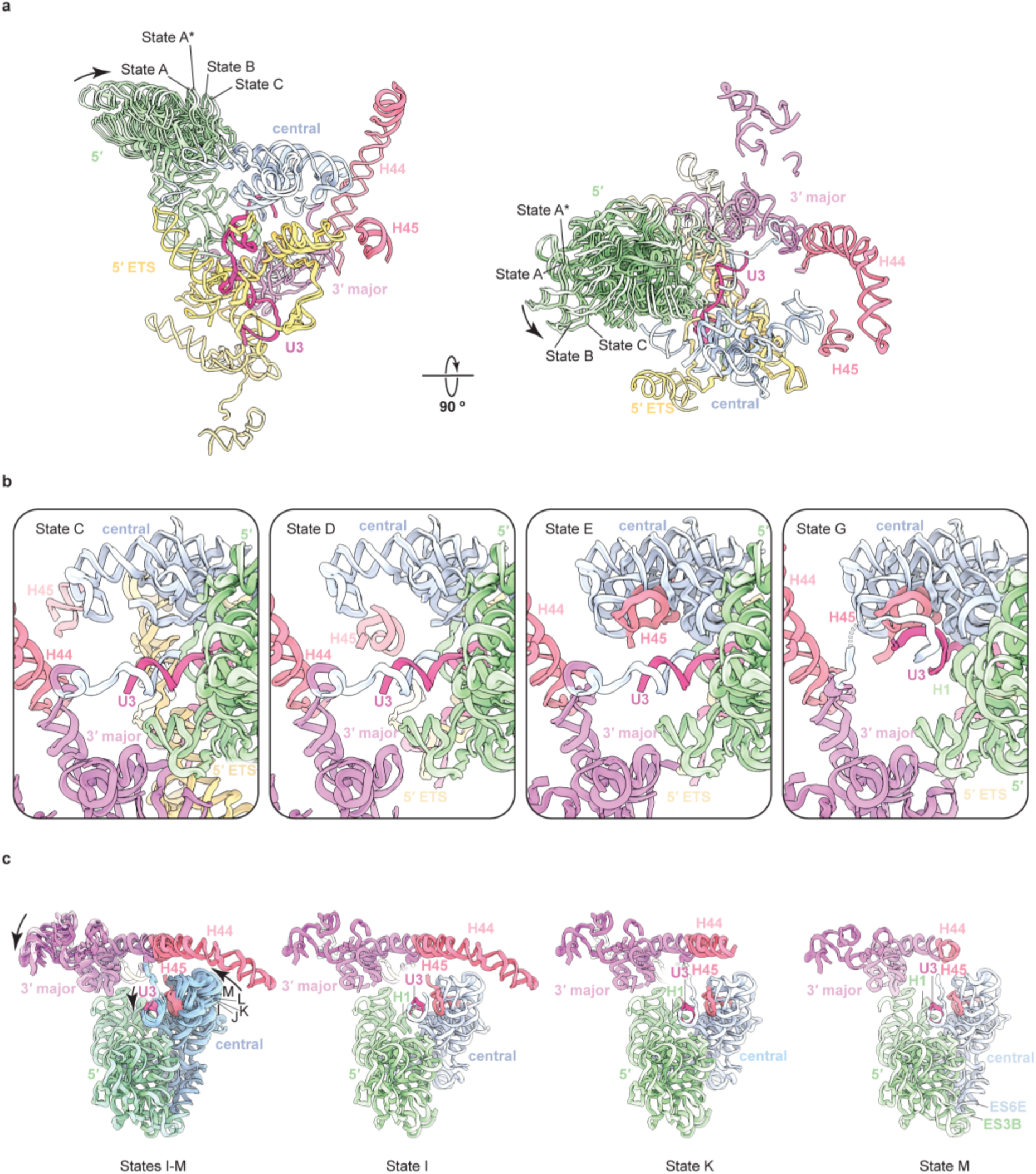
rRNA compaction and folding events across states. **(a)** Compaction of the 18*S* 5′ domain from States A-C side view (left) and top view (right). Labeled RNA elements are color coded. Arrows highlight compaction between states A and C. **(b)** Simplified rRNA folding events leading up to A1 cleavage. **(c)** Compaction of the central and 5′ domains during states I-M.

**Extended Data Fig. 3.**
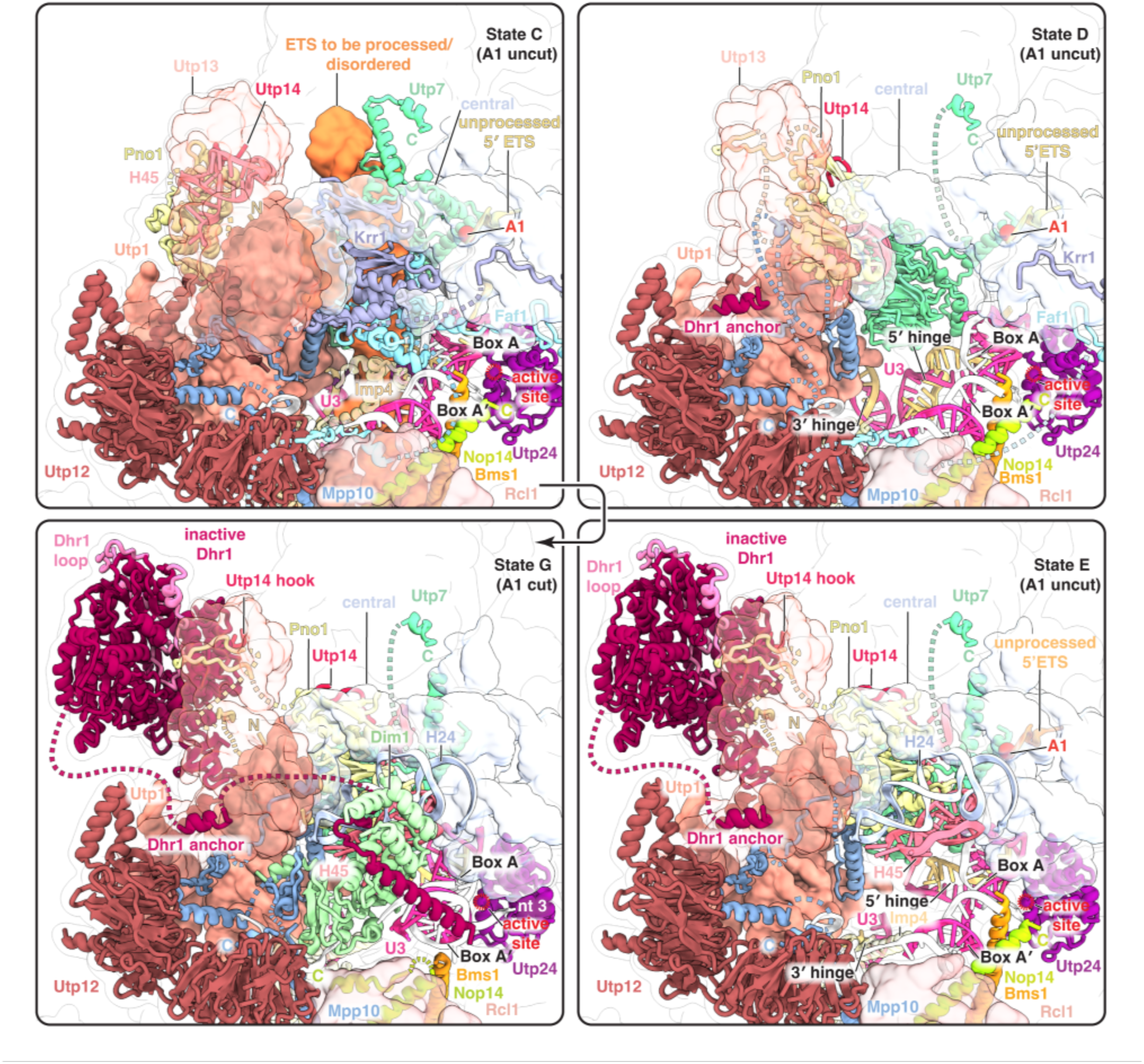
Molecular proofreading for A1 cleavage during SSU processome maturation. Detailed views involving states prior to (States C D E) and after A1 cleavage (State G) with color-coded proteins. The A1 cleavage site U3 architectural sites and active site of Utp24 are highlighted and flexible interactions indicated by dashed lines.

**Extended Data Fig. 4.**
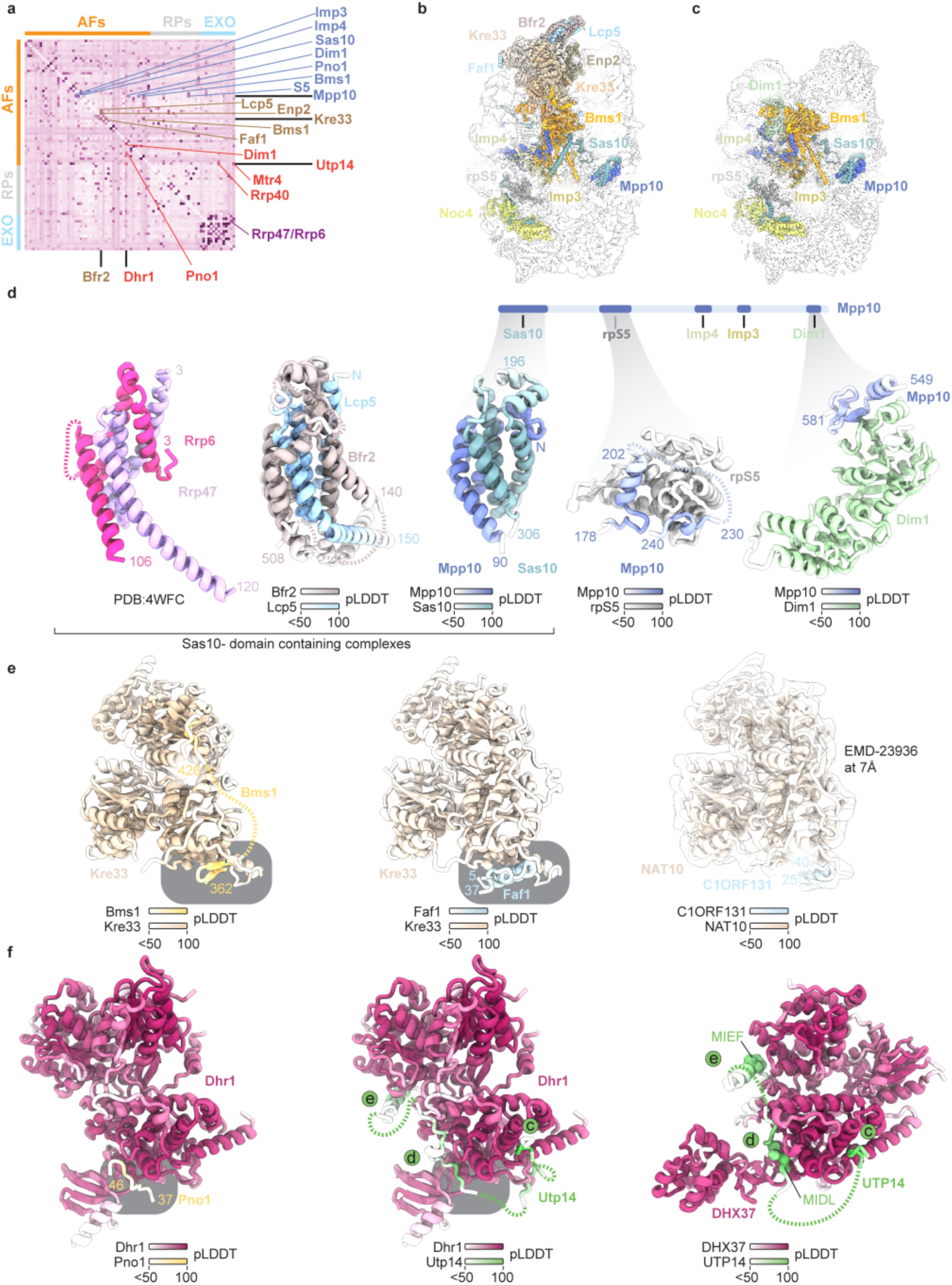
Yeast SSU processome predictome. **(a)** Color coded yeast SSU processome predictome between assembly factors (AFs) ribosomal proteins (RPs) and RNA exosome proteins (EXO). Key interactors of Mpp10 Kre33 Utp14 Dhr1 and Bfr2 are color coded. **(b)** State C containing proteins listed in (a). **(c)** State H containing proteins listed in (a). **(d)** Sas10-domain containing Rrp6/Rrp47 complex and predicted Bfr2-Lcp5 and Mpp10-Sas10 complexes. The schematic of Mpp10 contextualizes predictions of Mpp10 interactors. **(e)** Mutually exclusive interactions with Kre33 (human NAT10) predicted for Bms1 and Faf1 (human C1orf131). **(f)** Predictions of Dhr1/DHX37 interactions. Mutually exclusive binding of Pno1 and Utp14 to yeast Dhr1 are highlighted. Elements of human UTP14 known to activate human DHX37 (MIDL MIEF)^19^ are indicated. Confidence levels (pLDDT) are color coded for each protein.

**Extended Data Fig. 5.**
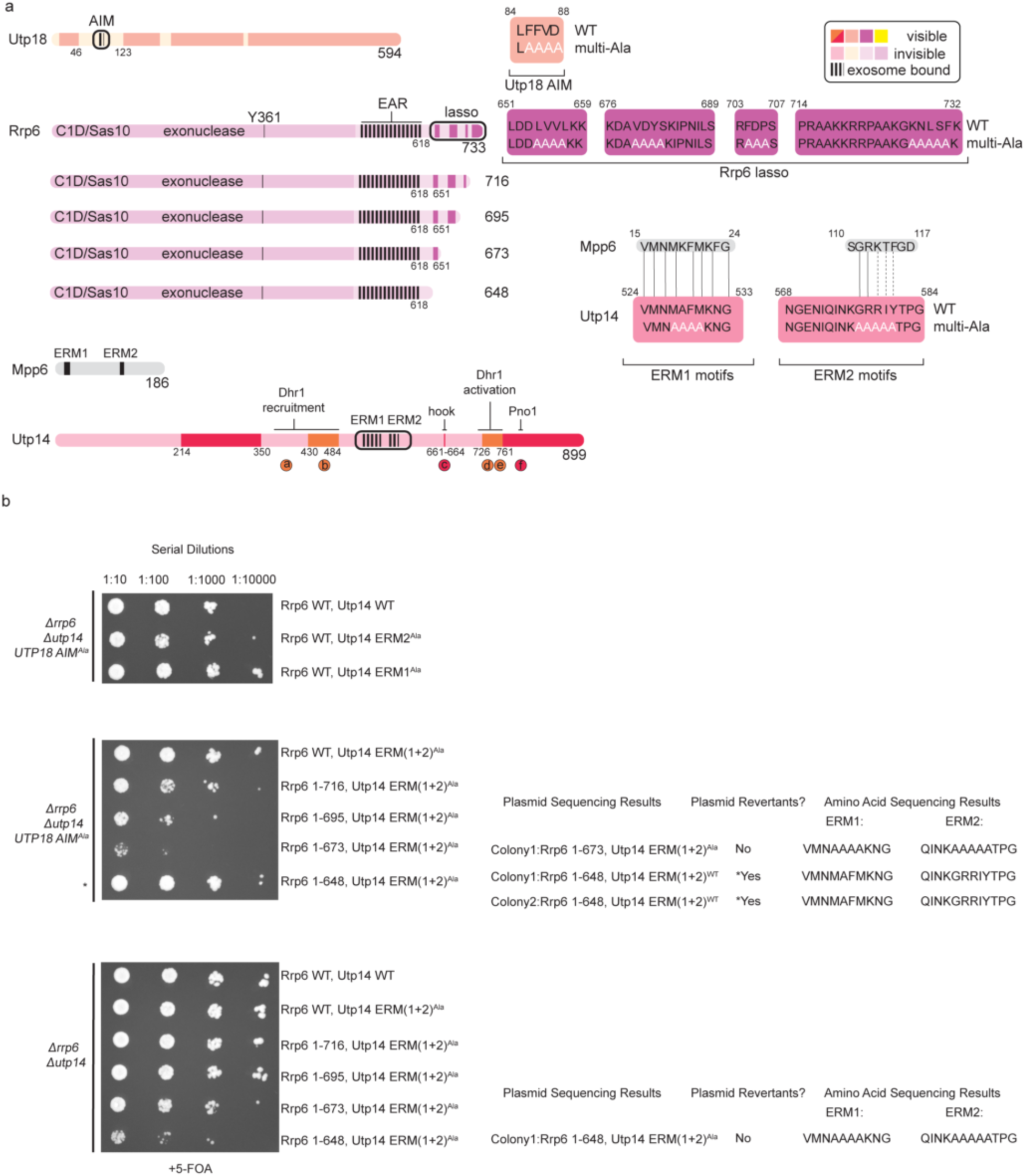
Yeast growth assays. **(a)** Schematics of Utp18 Rrp6 Mpp6 and Utp14 with interacting and functional elements labeled with multi-Ala mutations used in this study as well as an alignment to Mpp6 exosome recruitment motif indicated in grey. Designed truncations of Rrp6 lasso (bottom left) with interacting and functional elements labeled. Locations of all alanine substitutions for Utp18 Rrp6 and Utp14 are indicated on the right. **(b)** Growth analysis of Utp18 multi-Ala mutant in Δrrp6 and Δutp14 background (top and middle panels) and Utp18 WT in Δrrp6 and Δutp14 background (bottom panel). Growth assay using WT truncations and mutant Rrp6 or Utp14 constructs in the absence of genomic Rrp6 and Utp14. Colonies sequenced are indicated on the right of the growth assays

**Extended Data Fig. 6.**
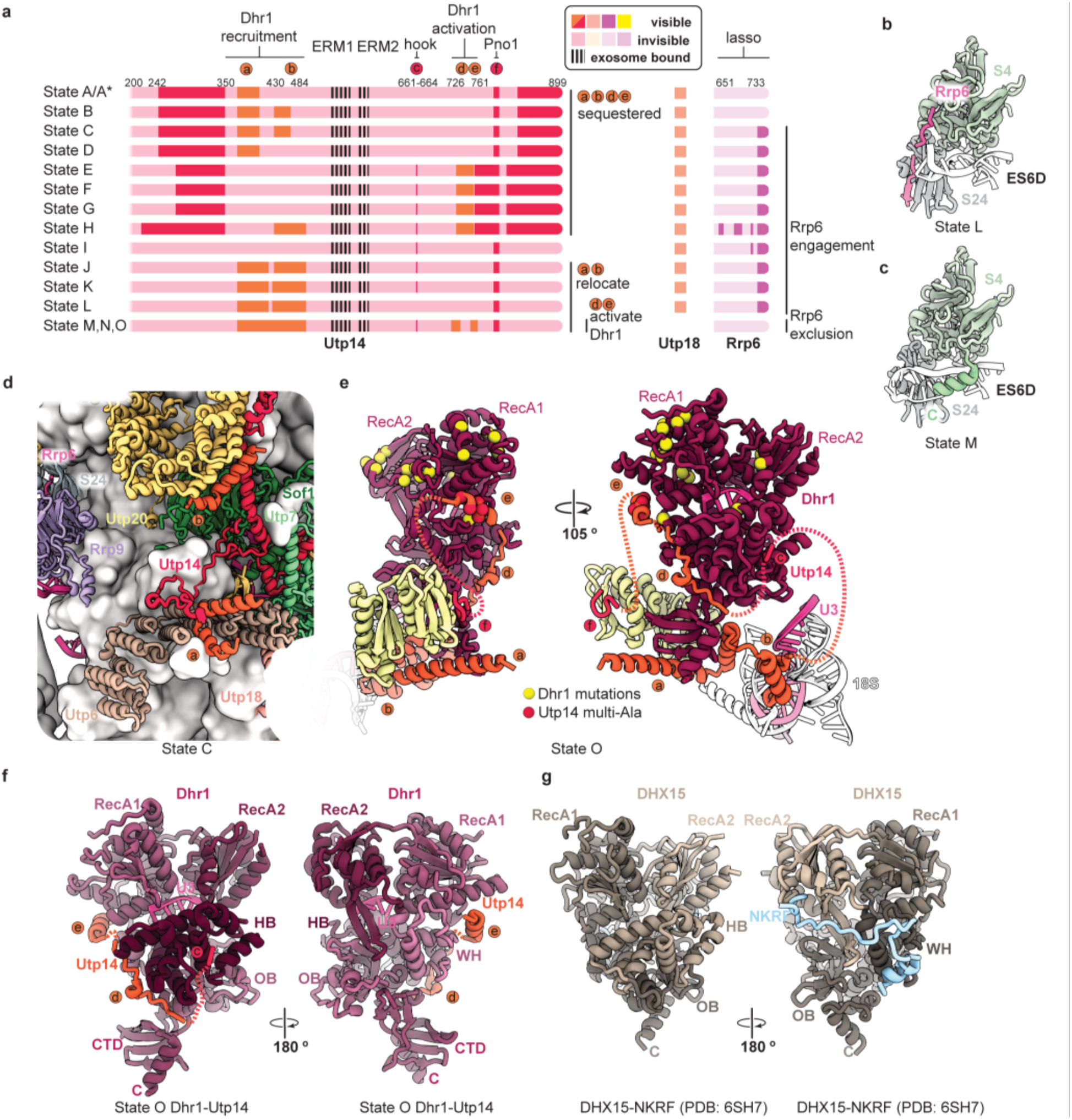
Functional elements of Utp14 Rrp6 Utp18 and Dhr1. **(a)** Schematics of Utp14 Utp18 and Rrp6 across States A-O. Utp14 functional sequence elements structured bound relocated or activating are indicated on the left and the presence/absence of Utp18 is indicated in the middle. Visible regions of the Rrp6 C-terminus (lasso) are indicated on the right. **(b)** Binding of Rrp6 lasso to ribosomal proteins S24 in State L. **(c)** Rrp6 exclusion by ES6D and S4 in State M. **(d)** State C highlights the initial binding of the Rrp6 lasso and the initial sequestration of Utp14 elements a and b that later recruit Dhr1. (**e)** Previously characterized mutations^1842^ mapped onto Dhr1 and Utp14 represented in State O. Functional elements of Utp14 are shown in orange. **(f)** Mode of Dhr1 activation by Utp14 in State O. **(g)** Mode of activation of DHX15 by G-patch protein NKRF (PDB: 6SH7).

