## Supplementary Information for "Helicase-mediated mechanism of SSU processome maturation and disassembly"

**TABLE OF CONTENTS**

Supplementary Fig. 1 - Cryo-EM processing of Dhr1 and Kre33 dataset.

Supplementary Fig. 2 - Cryo-EM processing of Dhr1 dataset.

Supplementary Fig. 3 - Cryo-EM processing of Dhr1 and Utp14 dataset.

Supplementary Fig. 4 - Cryo-EM focused maps and composite reconstruction of State A.

Supplementary Fig. 5 - Cryo-EM focused maps of State A continued.

Supplementary Fig. 6 - Cryo-EM focused maps and composite reconstruction of State A*.

Supplementary Fig. 7 - Cryo-EM focused maps of State A* continued.

Supplementary Fig. 8 - Cryo-EM focused maps and composite reconstruction of State B.

Supplementary Fig. 9 - Cryo-EM focused maps and composite reconstruction of State C.

Supplementary Fig. 10 - Cryo-EM focused maps and composite reconstruction of State D.

Supplementary Fig. 11 - Cryo-EM focused maps and composite reconstruction of State E.

Supplementary Fig. 12 - Cryo-EM focused maps and composite reconstruction of State F.

Supplementary Fig. 13 - Cryo-EM focused maps of State F continued.

Supplementary Fig. 14 - Cryo-EM focused maps and composite reconstruction of State G.

Supplementary Fig. 15 - Cryo-EM focused maps of State G continued.

Supplementary Fig. 16 - Cryo-EM focused maps and composite reconstruction of State H.

Supplementary Fig. 17 - Cryo-EM focused maps of State H continue.

Supplementary Fig. 18 - Cryo-EM focused maps and composite reconstruction of State I.

Supplementary Fig. 19 - Cryo-EM focused maps of State I continued.

Supplementary Fig. 20 - Cryo-EM focused maps and composite reconstruction of State J.

Supplementary Fig. 21- Cryo-EM focused maps and composite reconstruction of State K.

Supplementary Fig. 22 - Cryo-EM focused maps and composite reconstruction of State L.

Supplementary Fig. 23 - Cryo-EM focused maps and composite reconstruction of State M.

Supplementary Fig. 24 - Cryo-EM focused maps and composite reconstruction of State N.

Supplementary Fig. 25 - Cryo-EM focused maps and composite reconstruction of State O.

Supplementary Fig. 26 - Comparative view of yeast SSU maturation and disassembly pathways.

Supplementary Fig. 27- Representative Cryo-EM densities and models.

Supplementary Fig. 28 - Uncropped yeast growth assays.

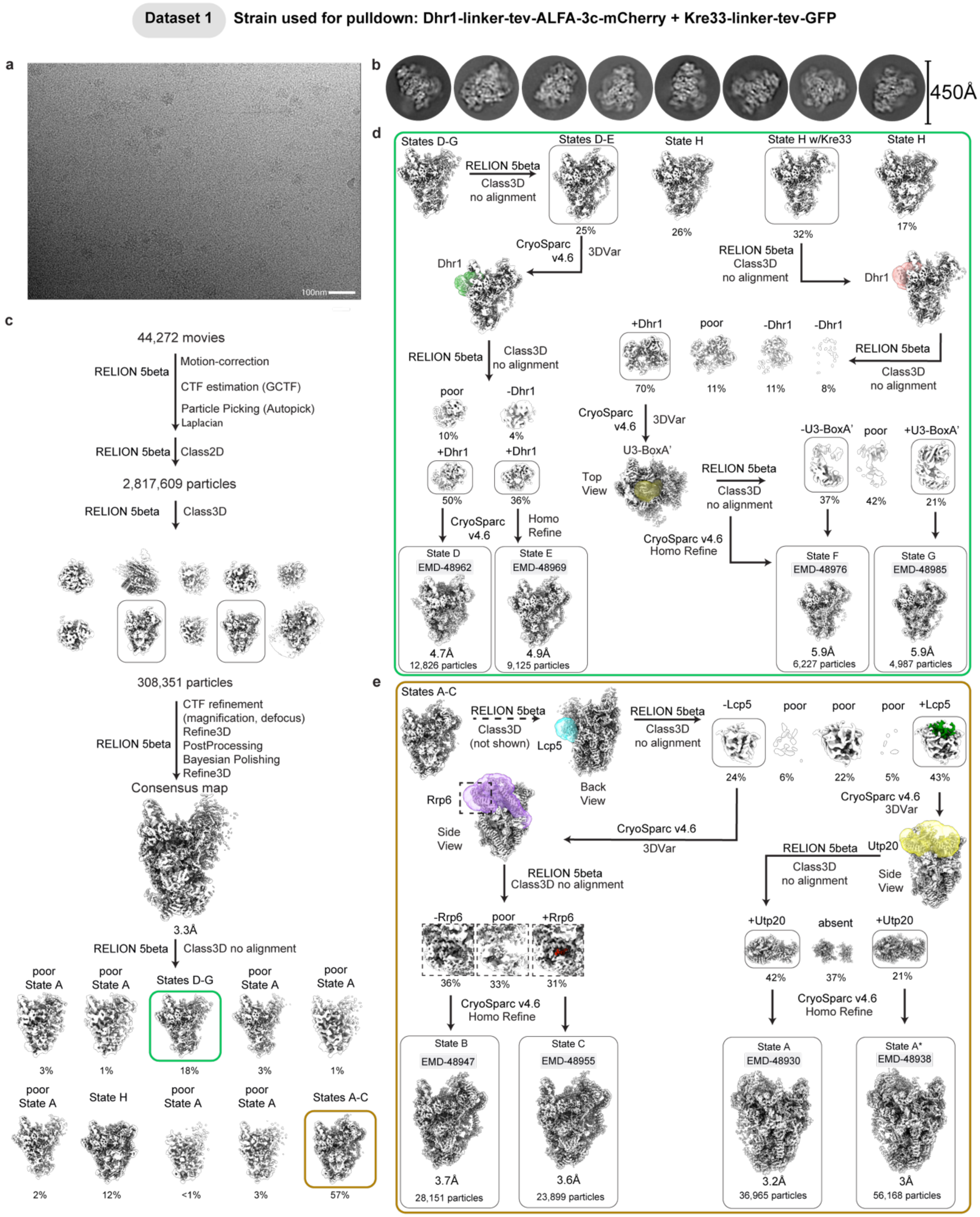


**Supplementary Fig. 1. Cryo-EM processing of Dhr1 and Kre33 dataset.**
(**a**) Representative motion corrected cryo-EM micrograph from 44,272 total micrographs. (**b**) 8 representative 2D class averages (4x binned, 4.32Å/px). (**c**) Initial data processing workflow for determination of a consensus map and identification of 2 mixed state populations. (**d**) and (**e**) Workflow for the identification and reconstruction of 8 SSU processome intermediates (States A-G).

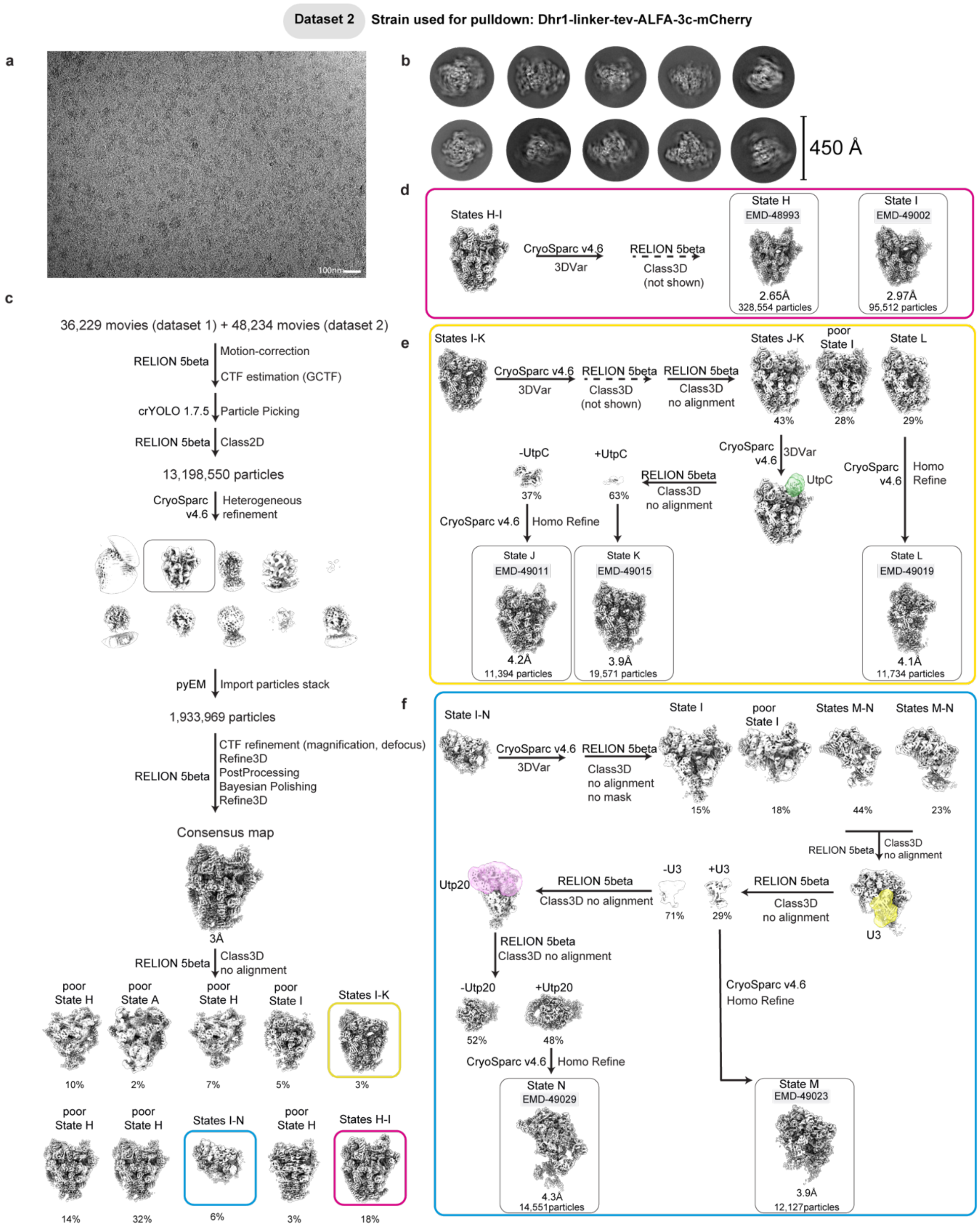


**Supplementary Fig. 2. Cryo-EM processing of Dhr1 dataset.**
(**a**) Representative motion corrected cryo-EM micrograph from 84,463 total micrographs. (**b**) 12 representative 2D class averages (4x binned, 4.32Å/px). (**c**) Initial data processing workflow for determination of a consensus map and identification of 3 mixed state populations. (**d**), (**e**) and (**f**) Workflow for the identification and reconstruction of 7 SSU processome intermediates (States H-N).

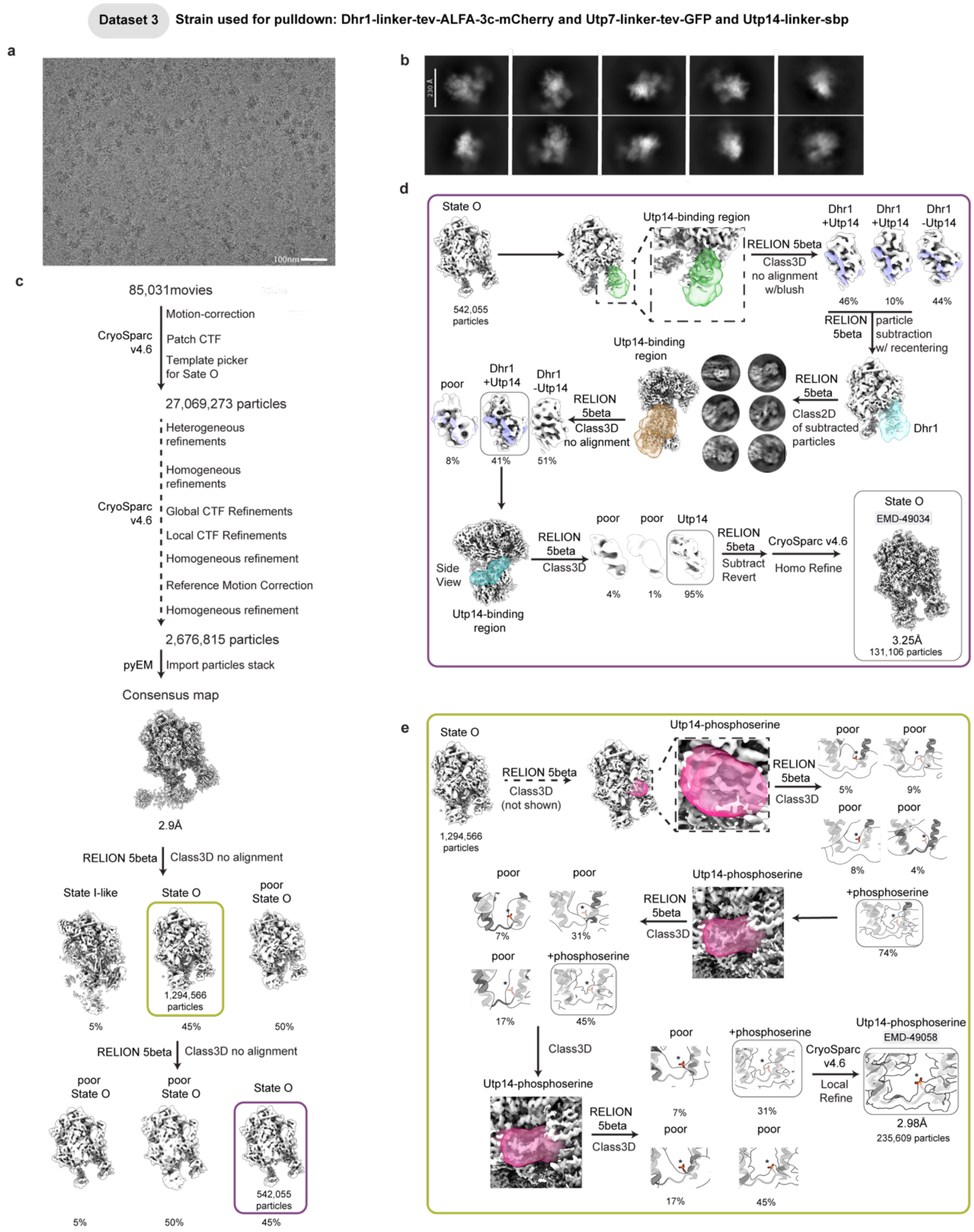


**Supplementary Fig. 3. Cryo-EM processing of Dhr1 and Utp14 dataset.**
(**a**) Representative motion corrected cryo-EM micrograph from 85,031 total micrographs. (**b**) 10 representative 2D class averages (4x binned, 4.32Å/px). (**c**) Initial data processing workflow for determination of a consensus map. (**d**) workflow for the identification and reconstruction of States O containing highest resolution Utp14 bound Dhr1. (**e**) Workflow for the identification of highest resolution focused map for Utp14-phoshoserine.

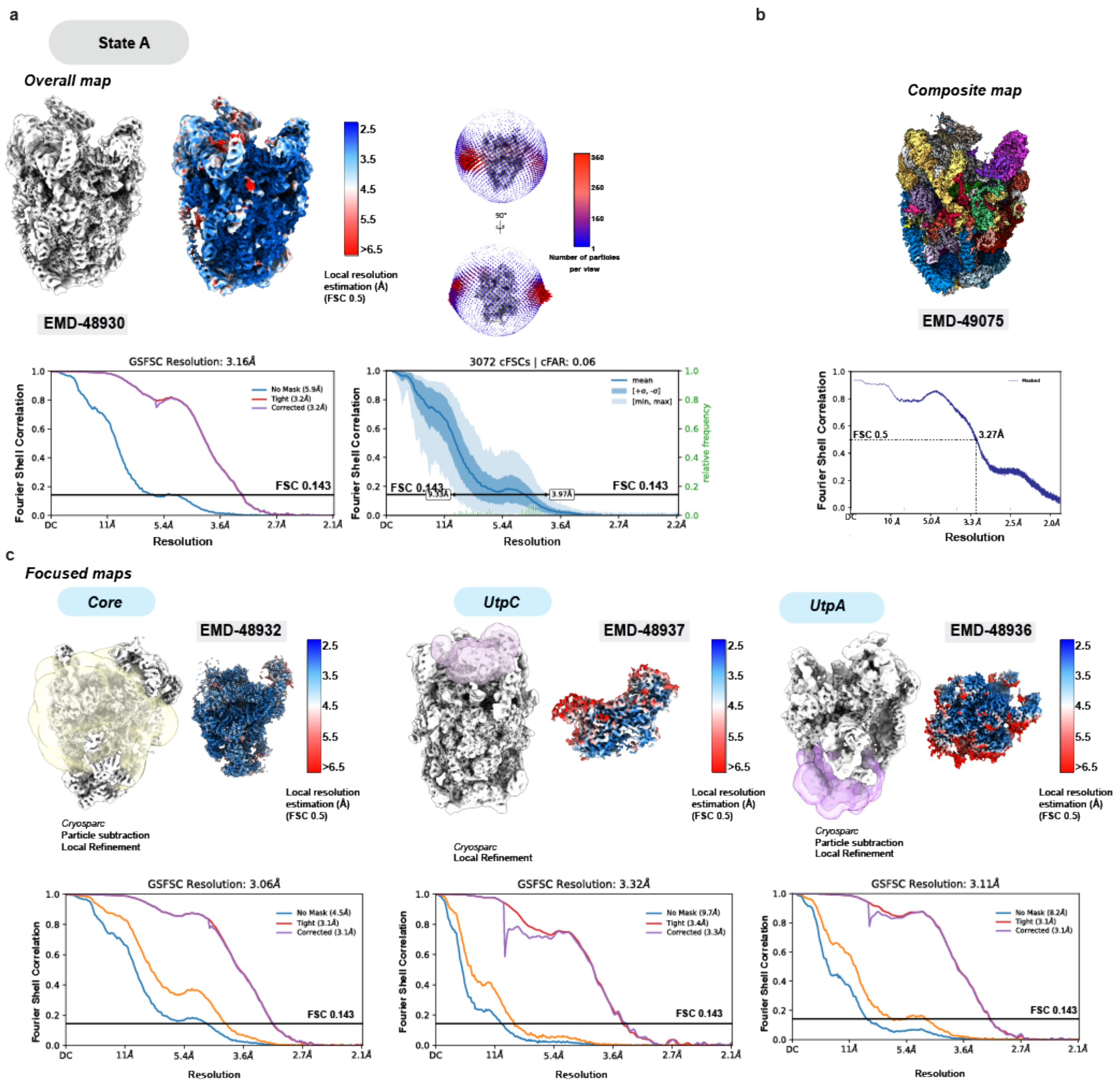

a
State A
Overall map
EMD-48930
Local resolution estimation (Å) (FSC 0.5)
Number of particles per view
GSFSC Resolution: 3.16Å
3072 cFSCs | cFAR: 0.06
FSC 0.143
Fourier Shell Correlation
Resolution
b
Composite map
EMD-49075
FSC 0.5
3.27Å
c
Focused maps
Core
EMD-48932
Cryosparc
Particle subtraction
Local Refinement
UtpC
EMD-48937
Cryosparc
Local Refinement
UtpA
EMD-48936
Cryosparc
Particle subtraction
Local Refinement
GSFSC Resolution: 3.06Å
GSFSC Resolution: 3.32Å
GSFSC Resolution: 3.11Å

**Supplementary Fig. 4. Cryo-EM focused maps and composite reconstruction of State A.** (**a**) Local-resolution filtered overall map of State A. FSC curves (no mask, tight mask and solvent corrected and 3D) are displayed on the bottom and Euler angle distribution is displayed on the right. (**b**) Composite map and FSC map-to-model curve displayed on the bottom. (**c**) Focused maps for the core, UtpC and UtpA modules used to generate the composite map are displayed along with the respective FSC curves (no mask, tight mask and solvent corrected) on the bottom.

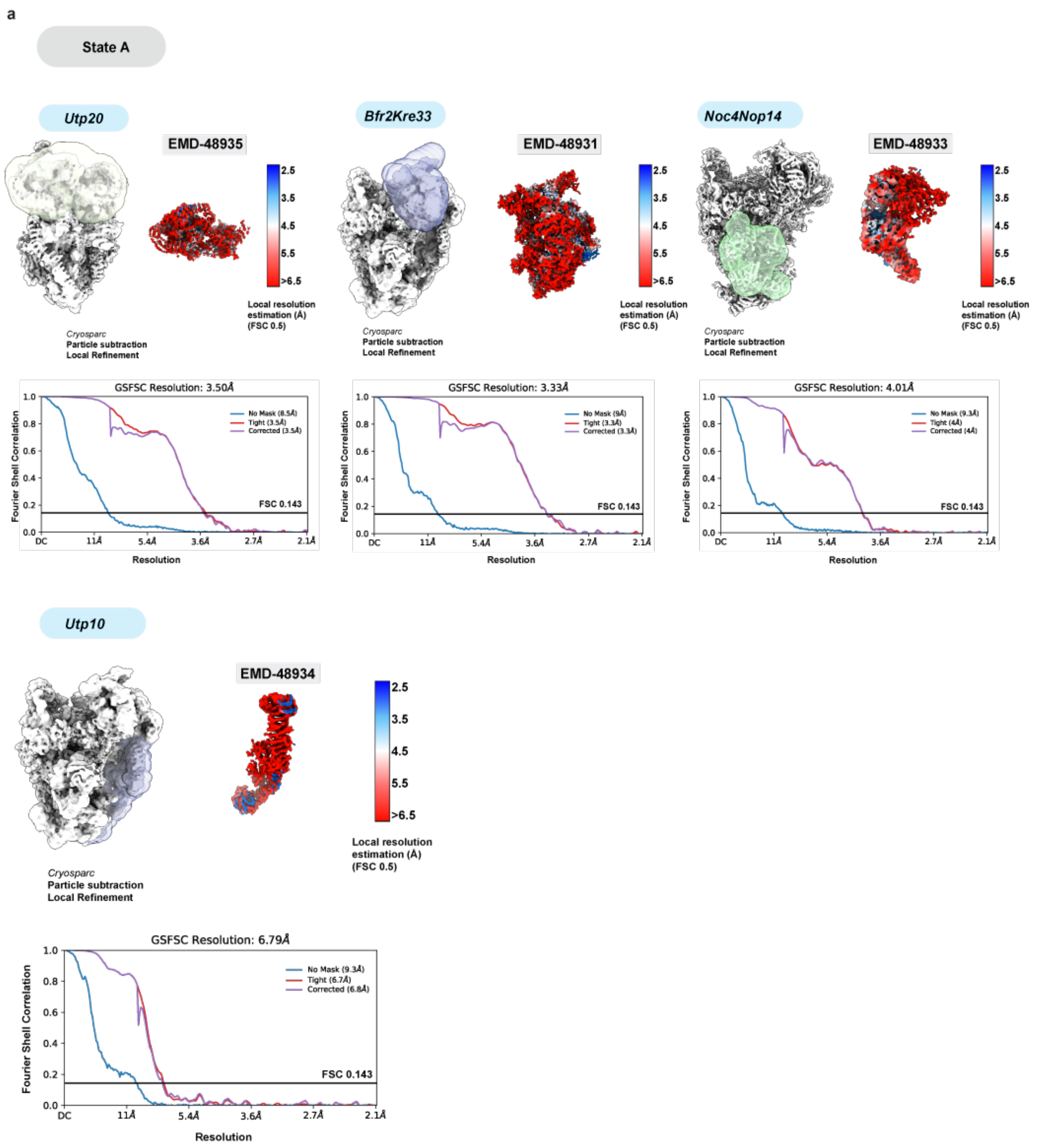


**Supplementary Fig. 5. Cryo-EM focused maps of State A continued.**
(**a**) Focused maps for the Utp20, Bfr2Kre33, Noc4Nop14, and Utp10 modules used to generate the composite map are displayed along with the respective FSC curves (no mask, tight mask and solvent corrected) on the bottom.

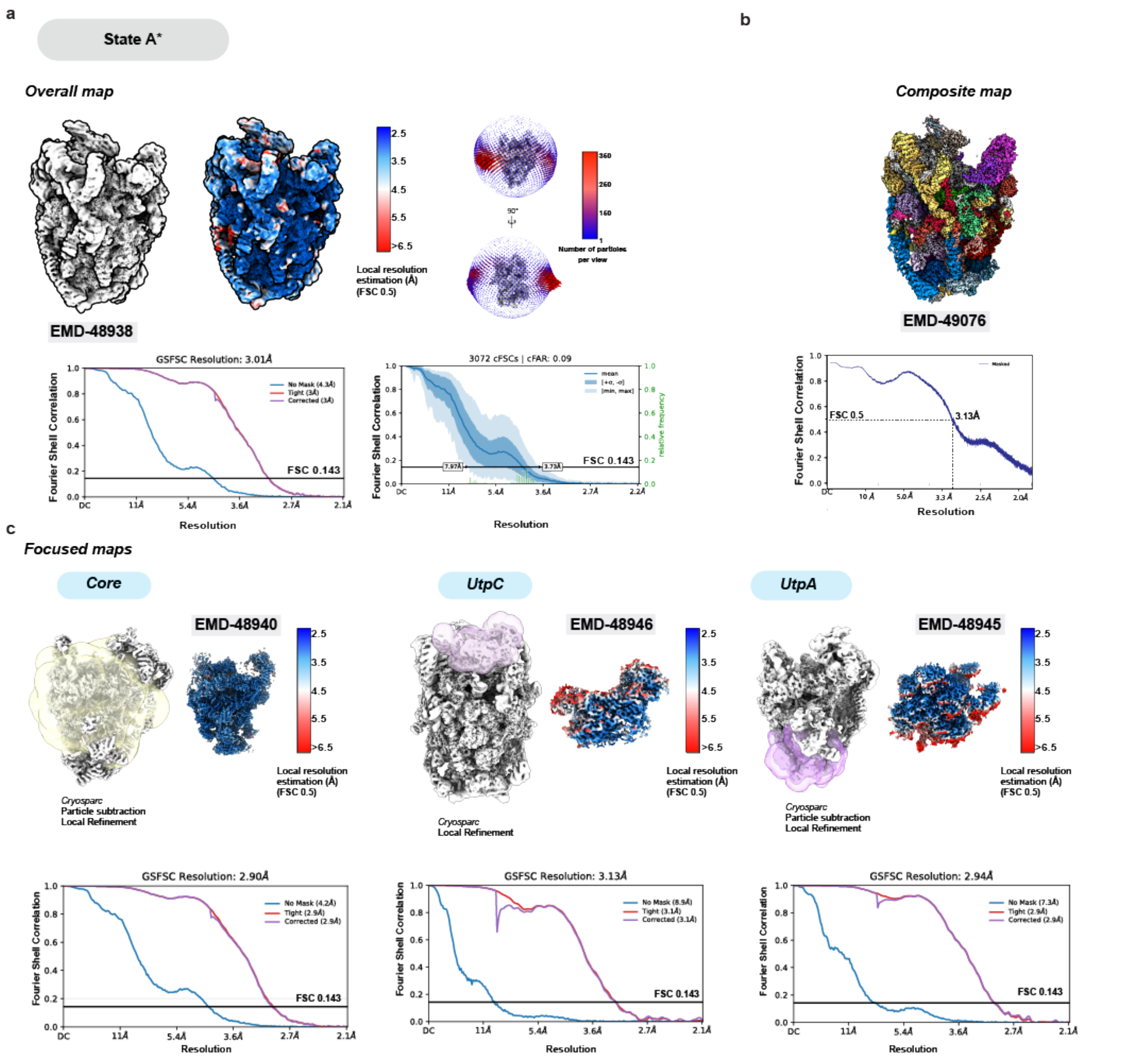


**Supplementary Fig. 6. Cryo-EM focused maps and composite reconstruction of State A*.** (**a**) Local-resolution filtered overall map of State A*. FSC curves (no mask, tight mask and solvent corrected and 3D) are displayed on the bottom and Euler angle distribution is displayed on the right. (**b**) Composite map and FSC map-to-model curve displayed on the bottom. (**c**) Focused maps for the core, UtpC and UtpA modules used to generate the composite map are displayed along with the respective FSC curves (no mask, tight mask and solvent corrected) on the bottom.

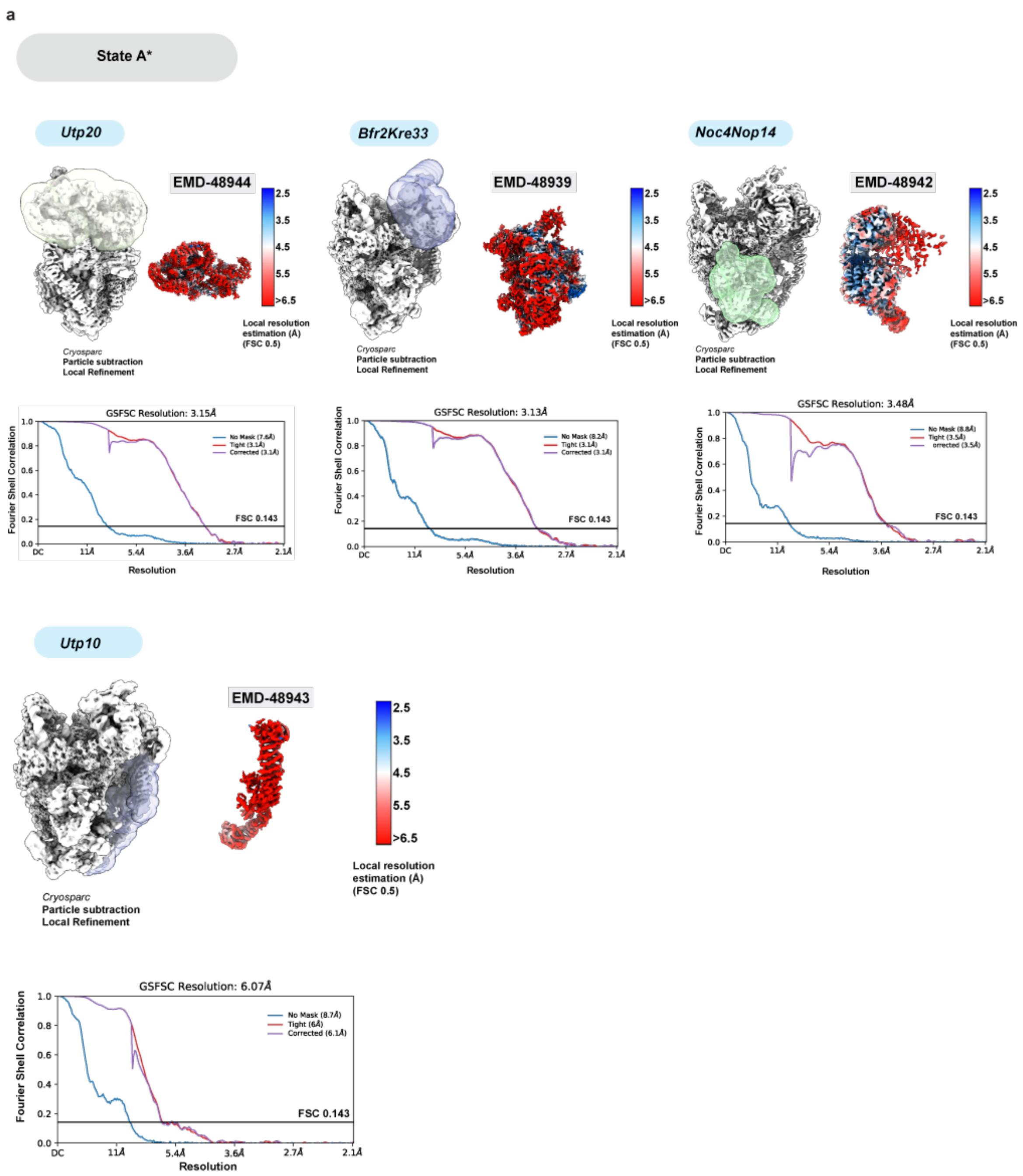


**Supplementary Fig. 7. Cryo-EM focused maps of State A* continued.**
(**a**) Focused maps for the Utp20, Bfr2Kre33, Noc4Nop14, and Utp10 modules used to generate the composite map are displayed along with the respective FSC curves (no mask, tight mask and solvent corrected) on the bottom.

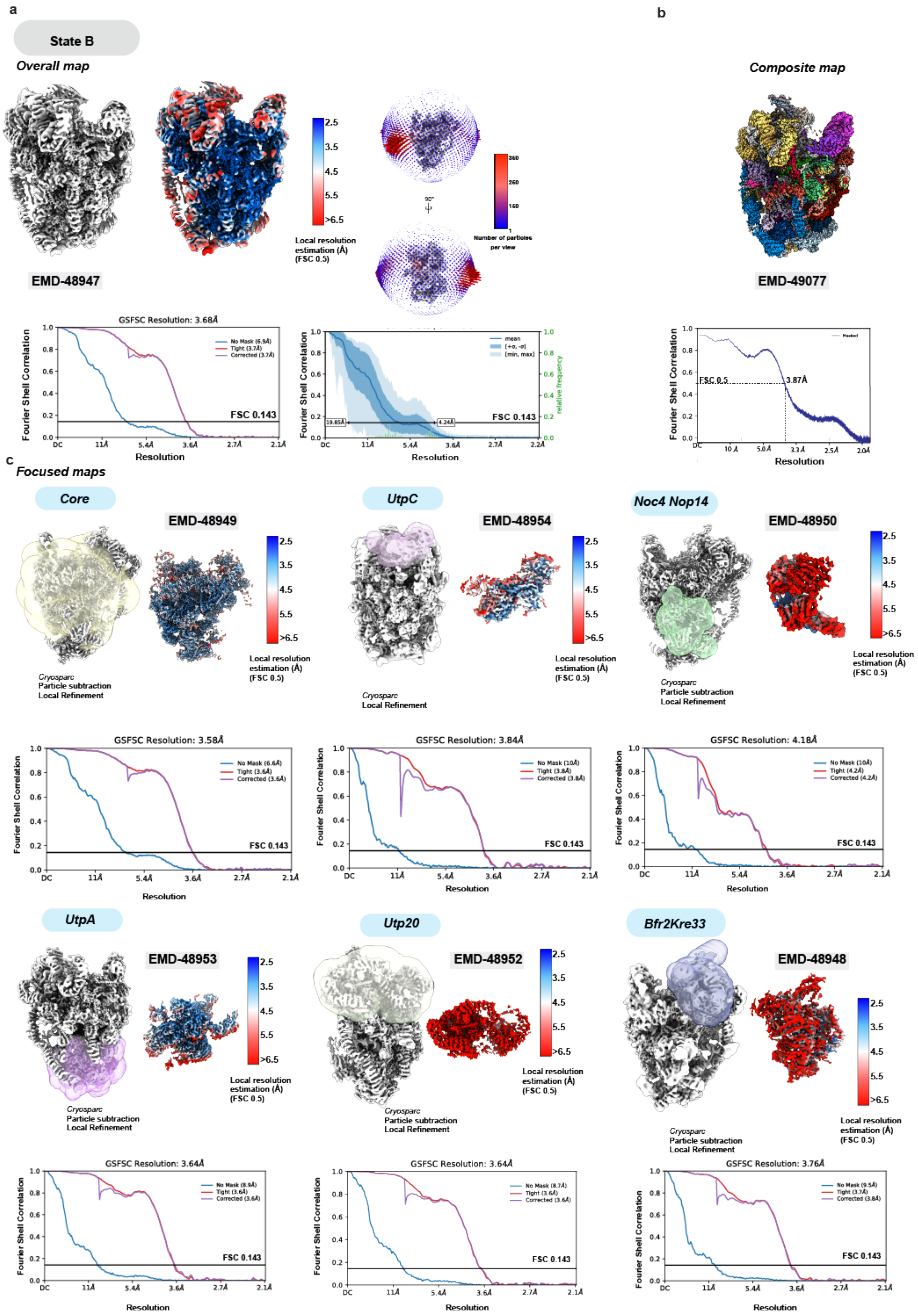
a
State B
Overall map
EMD-48947
Local resolution estimation (Å) (FSC 0.5)
Number of particles per view
GSFSC Resolution: 3.68Å
FSC 0.143
Fourier Shell Correlation
Resolution
b
Composite map
EMD-49077
FSC 0.5
3.87Å
c
Focused maps
Core
EMD-48949
Cryosparc
Particle subtraction
Local Refinement
UtpC
EMD-48954
Cryosparc
Local Refinement
Noc4 Nop14
EMD-48950
GSFSC Resolution: 3.58Å
GSFSC Resolution: 3.84Å
GSFSC Resolution: 4.18Å
UtpA
EMD-48953
Utp20
EMD-48952
Bfr2Kre33
EMD-48948
GSFSC Resolution: 3.64Å
GSFSC Resolution: 3.64Å
GSFSC Resolution: 3.76Å

**Supplementary Fig. 8. Cryo-EM focused maps and composite reconstruction of State B.** (**a**) Local-resolution filtered overall map of State B. FSC curves (no mask, tight mask and solvent corrected and 3D) are displayed on the bottom and Euler angle distribution is displayed on the right. (**b**) Composite map and FSC map-to-model curve displayed on the bottom. (**c**) Focused maps for the core, UtpC, Noc4Nop14, UtpA, Utp20, and Bfr2Kre33 modules used to generate the composite map are displayed along with the respective FSC curves (no mask, tight mask and solvent corrected) on the bottom.

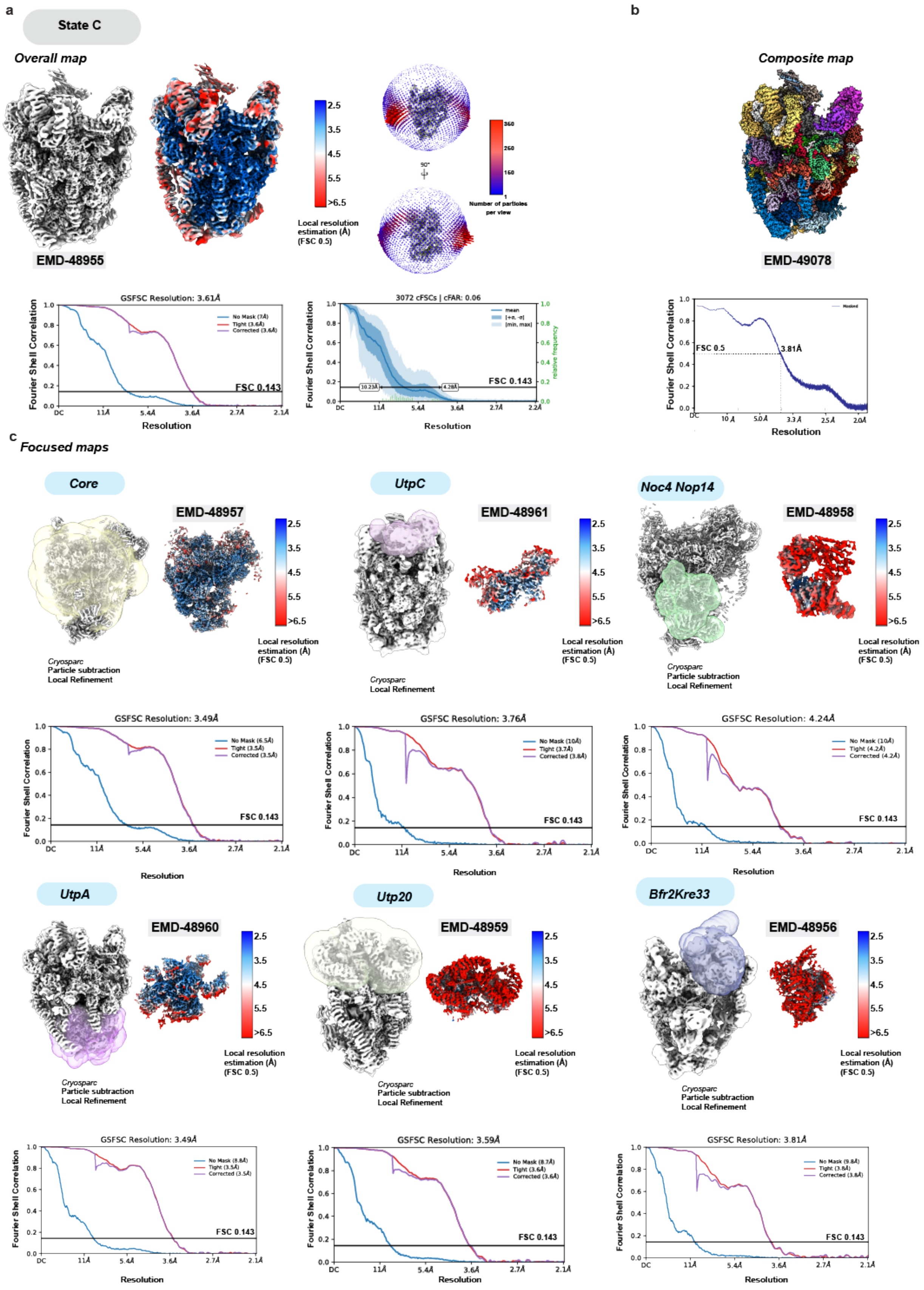

a
State C
Overall map
EMD-48955
Local resolution estimation (Å) (FSC 0.5)
Number of particles per view
GSFSC Resolution: 3.61Å
3072 cFSCs | cFAR: 0.06
FSC 0.143
Fourier Shell Correlation
Resolution
b
Composite map
EMD-49078
FSC 0.5
3.81Å
c
Focused maps
Core
EMD-48957
Cryosparc
Particle subtraction
Local Refinement
UtpC
EMD-48961
Cryosparc
Local Refinement
Noc4 Nop14
EMD-48958
GSFSC Resolution: 3.49Å
GSFSC Resolution: 3.76Å
GSFSC Resolution: 4.24Å
UtpA
EMD-48960
Utp20
EMD-48959
Bfr2Kre33
EMD-48956
GSFSC Resolution: 3.49Å
GSFSC Resolution: 3.59Å
GSFSC Resolution: 3.81Å

**Supplementary Fig. 9. Cryo-EM focused maps and composite reconstruction of State C.** (**a**) Local-resolution filtered overall map of State C. FSC curves (no mask, tight mask and solvent corrected and 3D) are displayed on the bottom and Euler angle distribution is displayed on the right. (**b**) Composite map and FSC map-to-model curve displayed on the bottom. (**c**) Focused maps for the core, UtpC, Noc4Nop14, UtpA, Utp20, and Bfr2Kre33 modules used to generate the composite map are displayed along with the respective FSC curves (no mask, tight mask and solvent corrected) on the bottom.

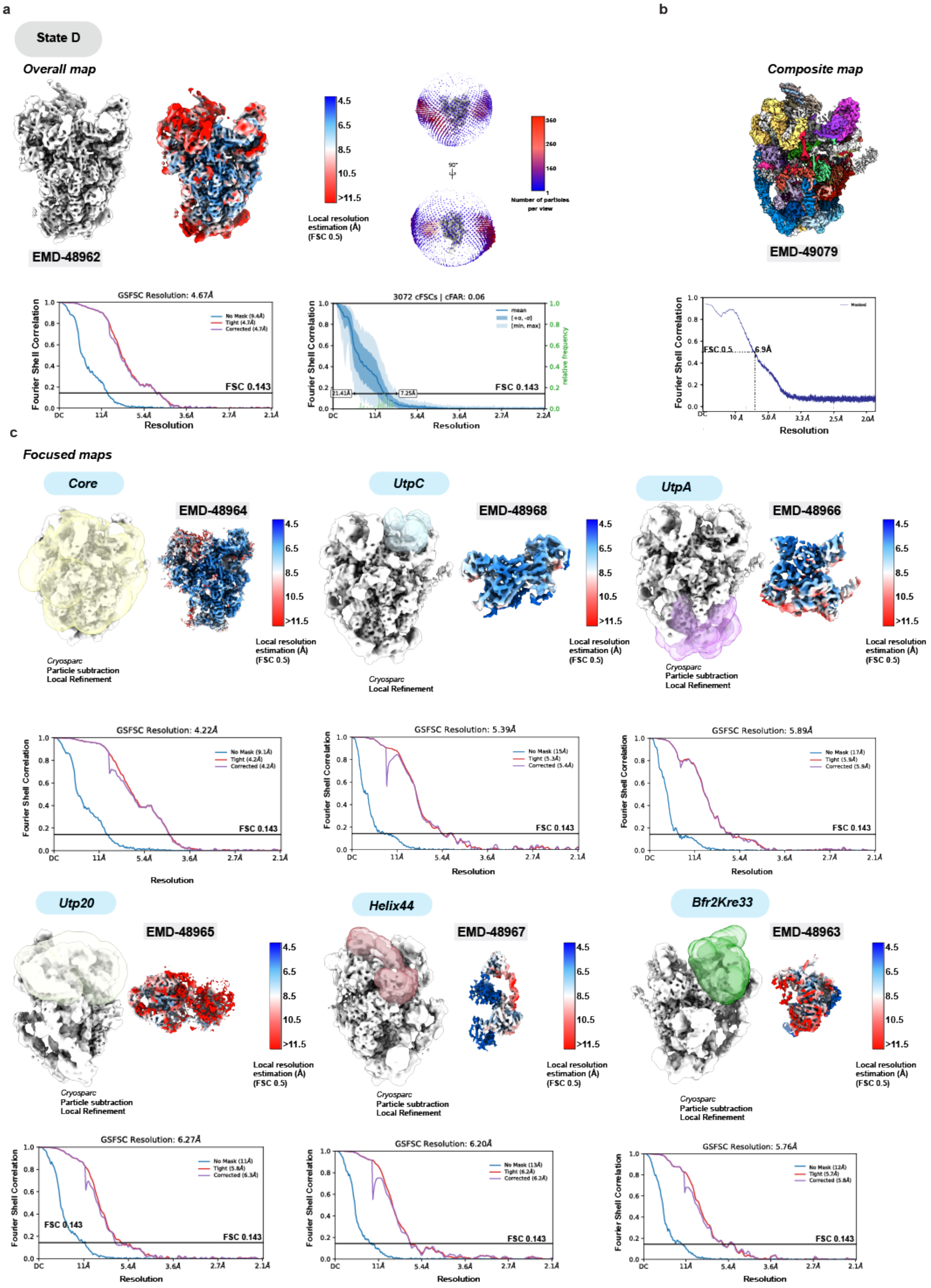

a
State D
Overall map
EMD-48962
Local resolution estimation (Å) (FSC 0.5)
Number of particles per view
GSFSC Resolution: 4.67Å
3072 cFSCs | cFAR: 0.06
FSC 0.143
Fourier Shell Correlation
Resolution
b
Composite map
EMD-49079
FSC 0.5
6.9Å
c
Focused maps
Core
EMD-48964
Cryosparc
Particle subtraction
Local Refinement
UtpC
EMD-48968
Cryosparc
Local Refinement
UtpA
EMD-48966
GSFSC Resolution: 4.22Å
GSFSC Resolution: 5.39Å
GSFSC Resolution: 5.89Å
Utp20
EMD-48965
Helix44
EMD-48967
Bfr2Kre33
EMD-48963
GSFSC Resolution: 6.27Å
GSFSC Resolution: 6.20Å
GSFSC Resolution: 5.76Å

**Supplementary Fig. 10. Cryo-EM focused maps and composite reconstruction of State D.** (**a**) Local-resolution filtered overall map of State D. FSC curves (no mask, tight mask and solvent corrected and 3D) are displayed on the bottom and Euler angle distribution is displayed on the right. (**b**) Composite map and FSC map-to-model curve displayed on the bottom. (**c**) Focused maps for the core, UtpC, UtpA, Utp20, Helix44 and Bfr2Kre33 modules used to generate the composite map are displayed along with the respective FSC curves (no mask, tight mask and solvent corrected) on the bottom.

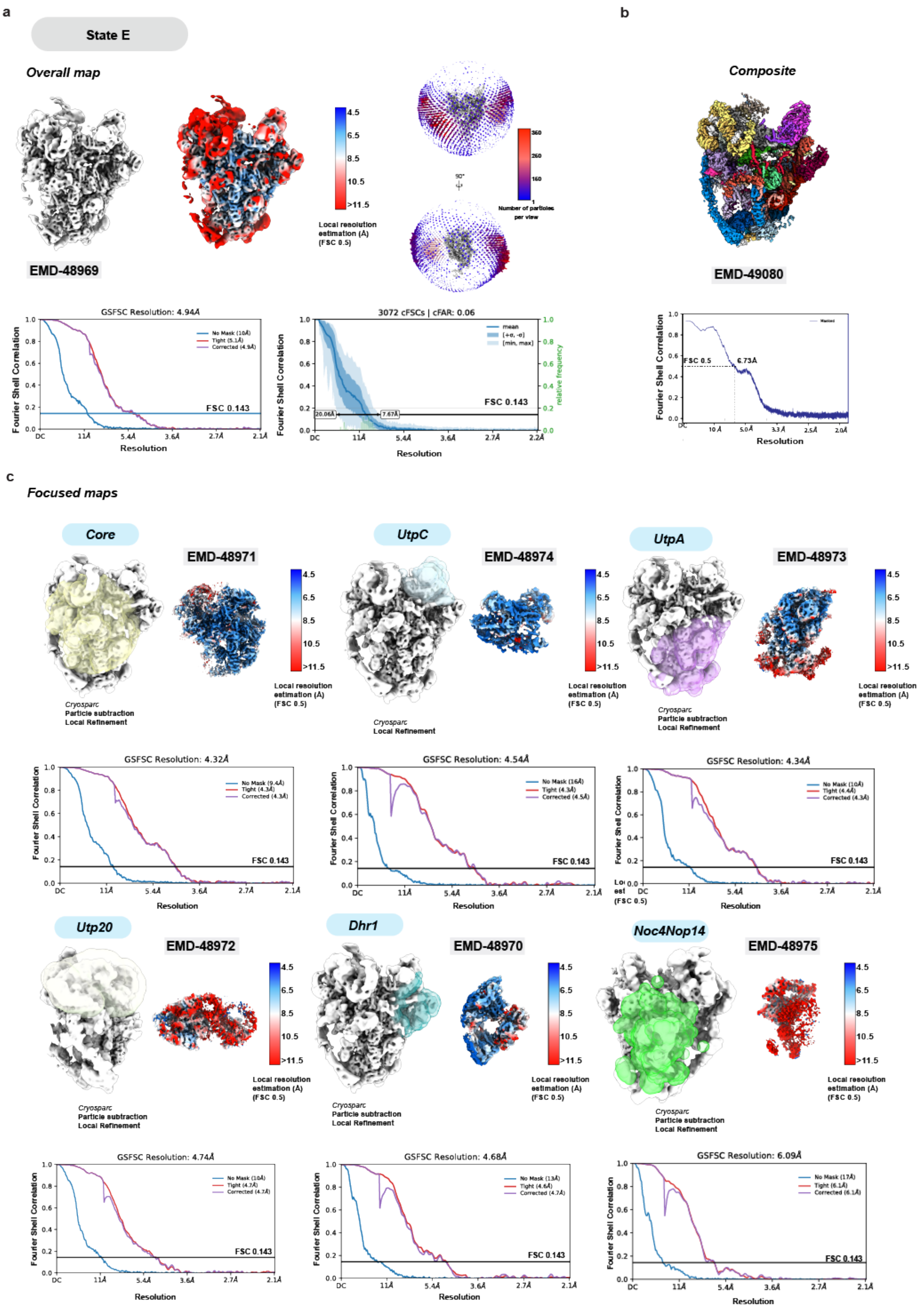

a
State E
Overall map
EMD-48969
4.5
6.5
8.5
10.5
>11.5
Local resolution estimation (Å) (FSC 0.5)
Number of particles per view
GSFSC Resolution: 4.94Å
No Mask (10Å)
Tight (5.1Å)
Corrected (4.9Å)
FSC 0.143
Fourier Shell Correlation
Resolution
3072 cFSCs | cFAR: 0.06
mean
relative frequency
20.06Å
7.67Å
b
Composite
EMD-49080
FSC 0.5
6.73Å
c
Focused maps
Core
EMD-48971
UtpC
EMD-48974
UtpA
EMD-48973
Cryosparc
Particle subtraction
Local Refinement
GSFSC Resolution: 4.32Å
GSFSC Resolution: 4.54Å
GSFSC Resolution: 4.34Å
Utp20
EMD-48972
Dhr1
EMD-48970
Noc4Nop14
EMD-48975
GSFSC Resolution: 4.74Å
GSFSC Resolution: 4.68Å
GSFSC Resolution: 6.09Å

**Supplementary Fig. 11. Cryo-EM focused maps and composite reconstruction of State E.** (**a**) Local-resolution filtered overall map of State E. FSC curves (no mask, tight mask and solvent corrected and 3D) are displayed on the bottom and Euler angle distribution is displayed on the right. (**b**) Composite map and FSC map-to-model curve displayed on the bottom. (**c**) Focused maps for the core, UtpC, UtpA, Utp20, Dhr1 and Noc4Nop14 modules used to generate the composite map are displayed along with the respective FSC curves (no mask, tight mask and solvent corrected) on the bottom.

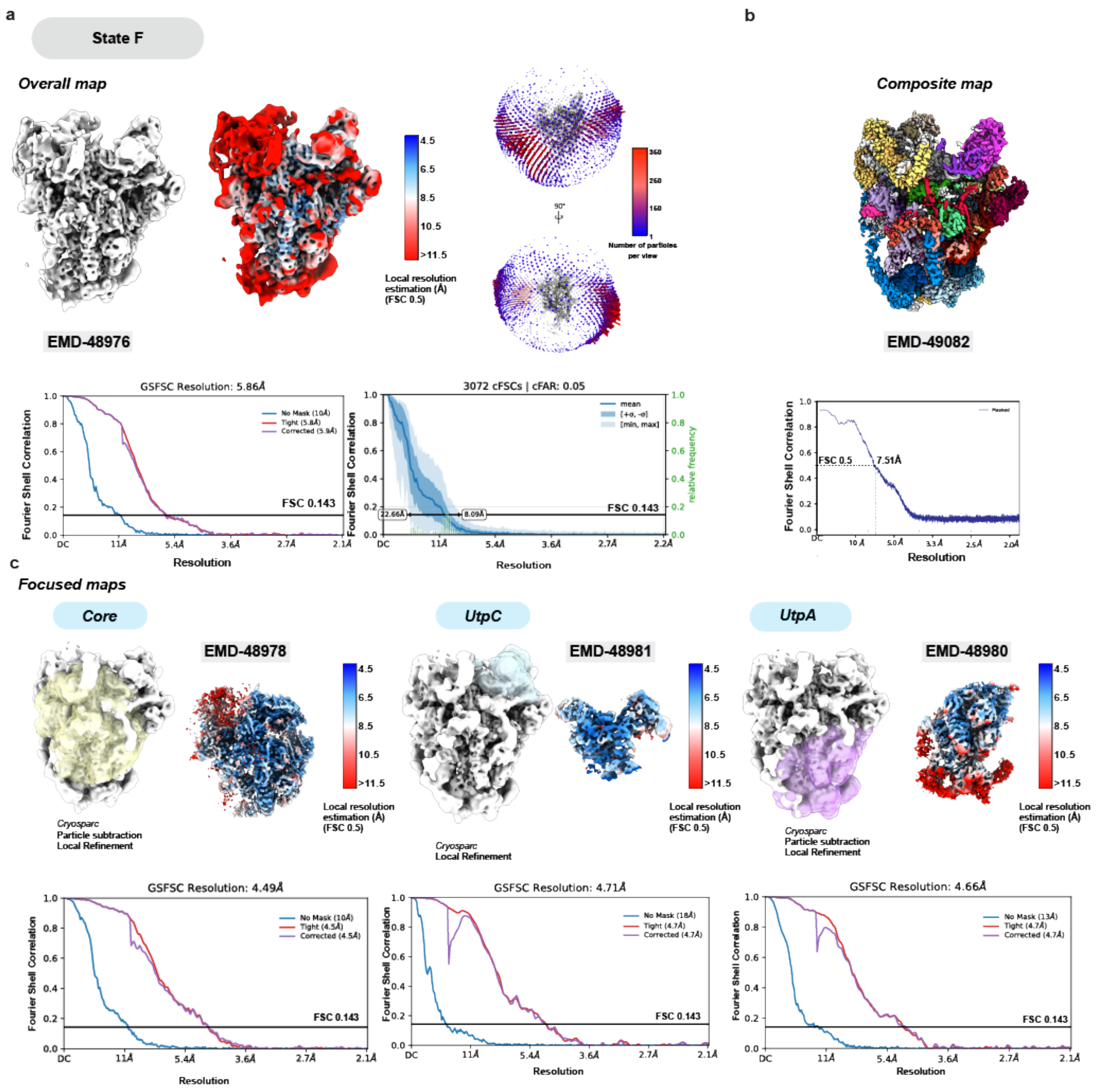


**Supplementary Fig. 12. Cryo-EM focused maps and composite reconstruction of State F.** (**a**) Local-resolution filtered overall map of State F. FSC curves (no mask, tight mask and solvent corrected and 3D) are displayed on the bottom and Euler angle distribution is displayed on the right. (**b**) Composite map and FSC map-to-model curve displayed on the bottom. (**c**) Focused maps for the core, UtpC and UtpA modules used to generate the composite map are displayed along with the respective FSC curves (no mask, tight mask and solvent corrected) on the bottom.

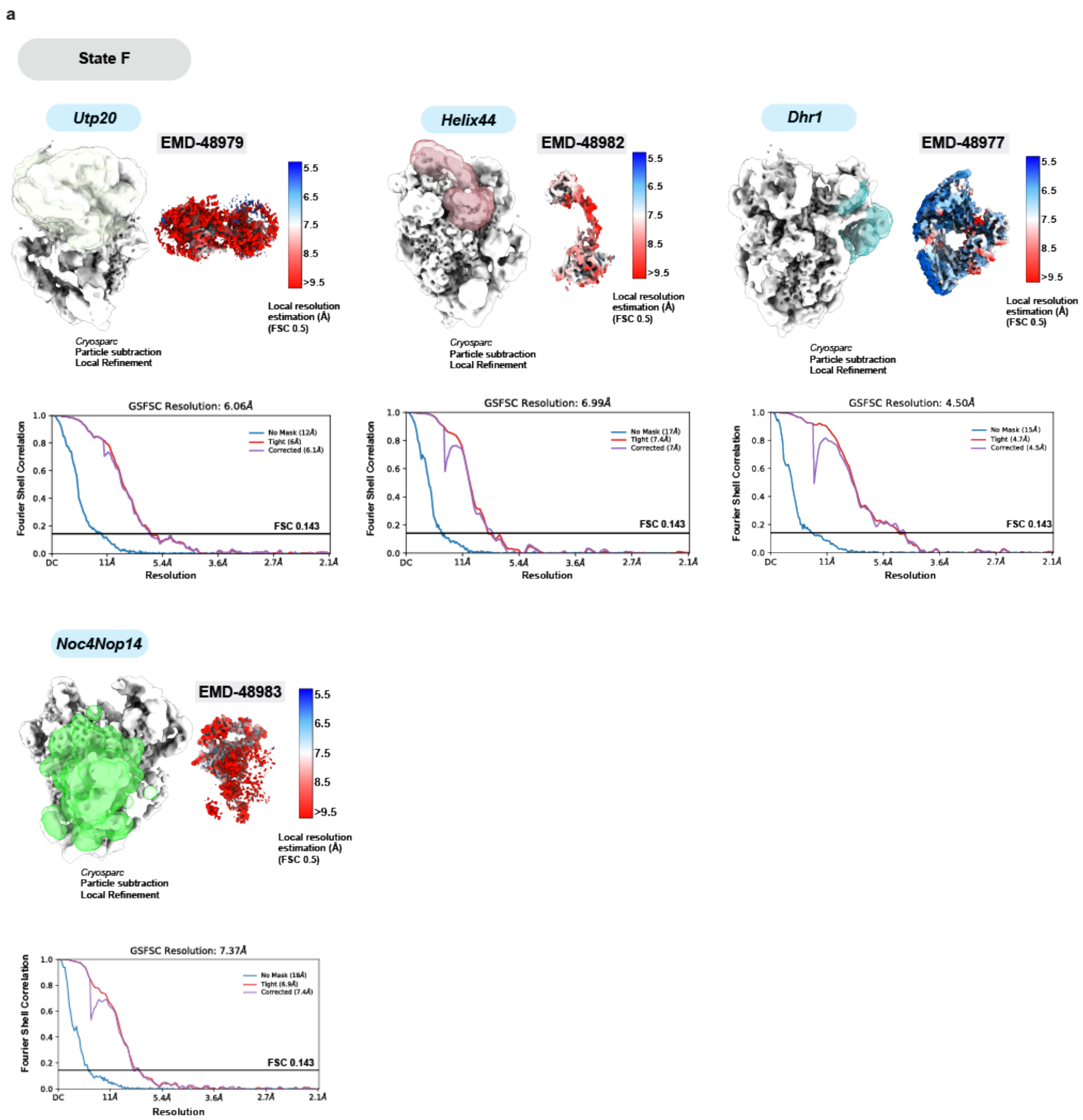


**Supplementary Fig. 13. Cryo-EM focused maps of State F continued.**
(**a**) Focused maps for the Utp20, Helix44, Dhr1 and Noc4Nop14 modules used to generate the composite map are displayed along with the respective FSC curves (no mask, tight mask and solvent corrected) on the bottom.

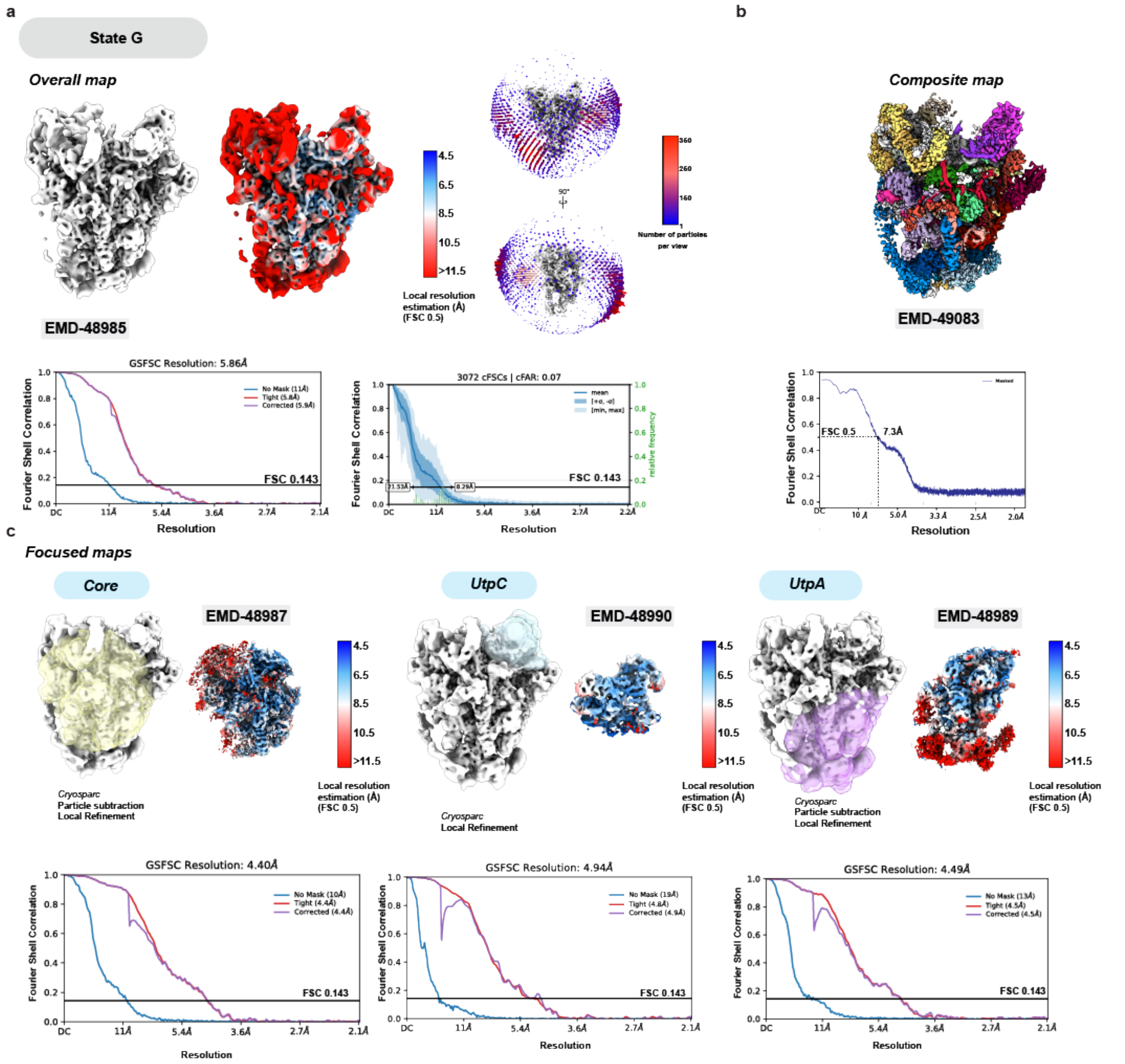


**Supplementary Fig. 14. Cryo-EM focused maps and composite reconstruction of State G.** (**a**) Local-resolution filtered overall map of State G. FSC curves (no mask, tight mask and solvent corrected and 3D) are displayed on the bottom and Euler angle distribution is displayed on the right. (**b**) Composite map and FSC map-to-model curve displayed on the bottom. (**c**) Focused maps for the core, UtpC and UtpA modules used to generate the composite map are displayed along with the respective FSC curves (no mask, tight mask and solvent corrected) on the bottom.

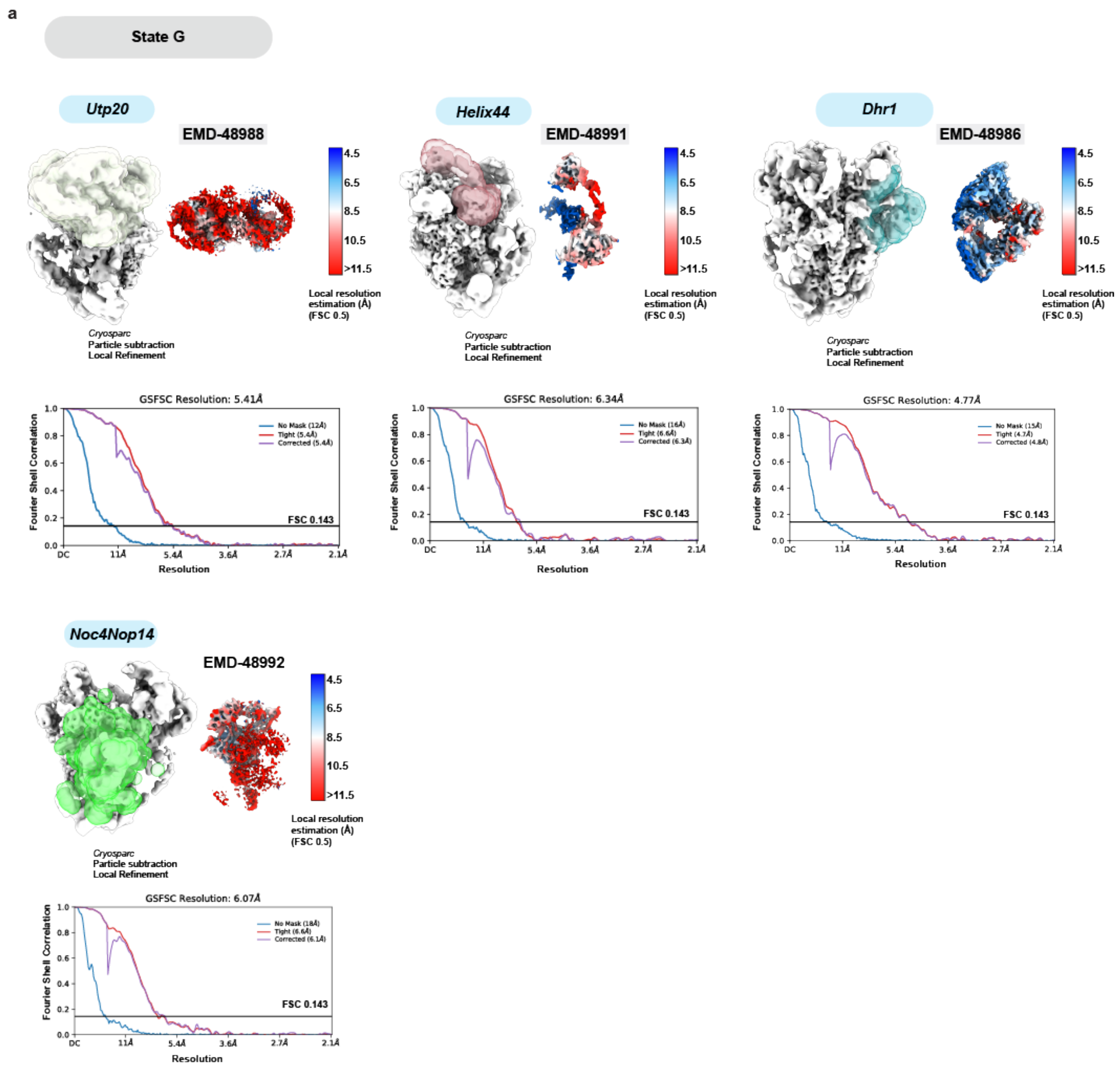


**Supplementary Fig. 15. Cryo-EM focused maps of State G continued.**
(**a**) Focused maps for the Utp20, Helix44, Dhr1 and Noc4Nop14 modules used to generate the composite map are displayed along with the respective FSC curves (no mask, tight mask and solvent corrected) on the bottom.

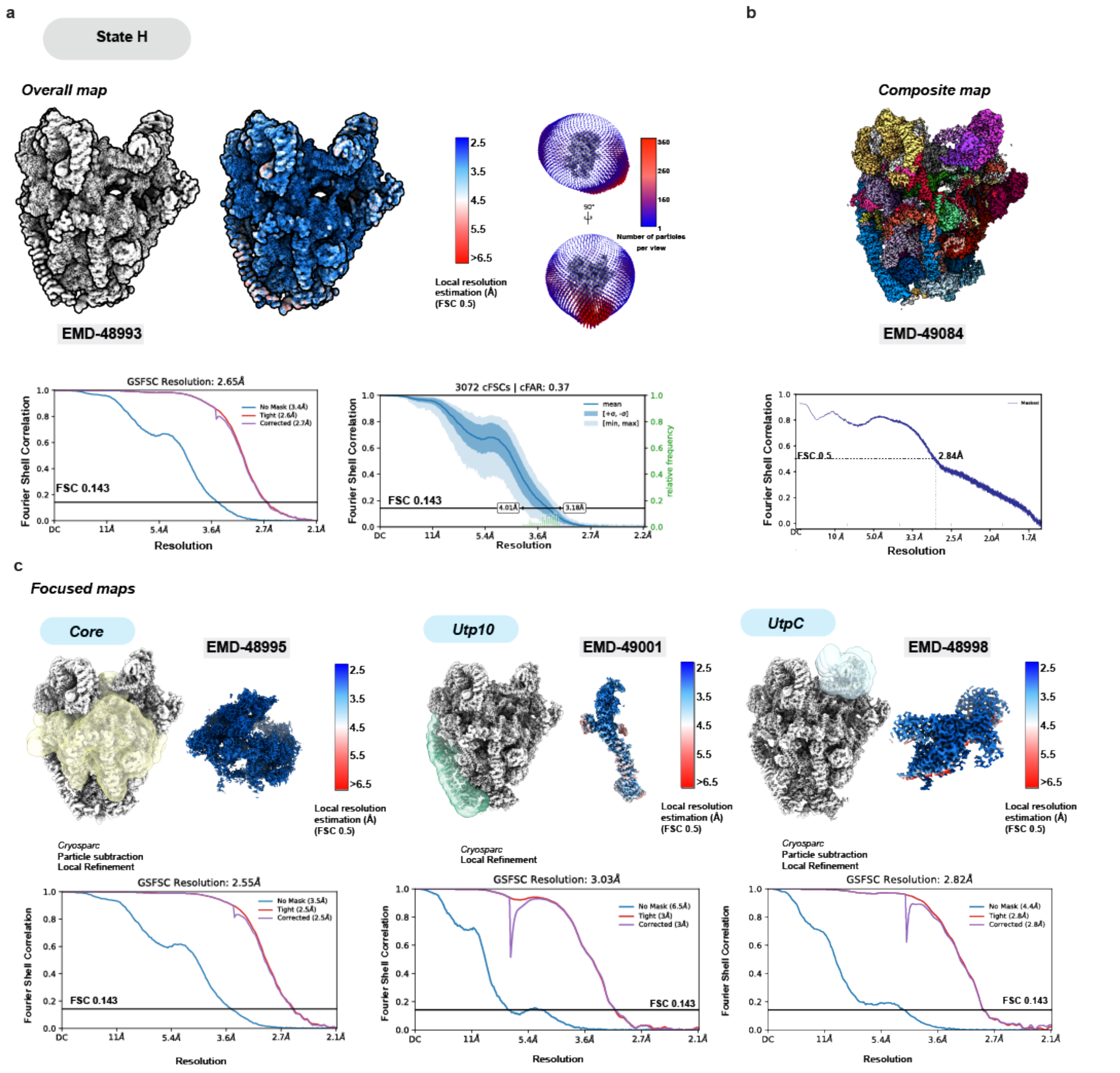


**Supplementary Fig. 16. Cryo-EM focused maps and composite reconstruction of State H.** (**a**) Local-resolution filtered overall map of State H. FSC curves (no mask, tight mask and solvent corrected and 3D) are displayed on the bottom and Euler angle distribution is displayed on the right. (**b**) Composite map and FSC map-to-model curve displayed on the bottom. (**c**) Focused maps for the core, Utp10 and UtpC modules used to generate the composite map are displayed along with the respective FSC curves (no mask, tight mask and solvent corrected) on the bottom.

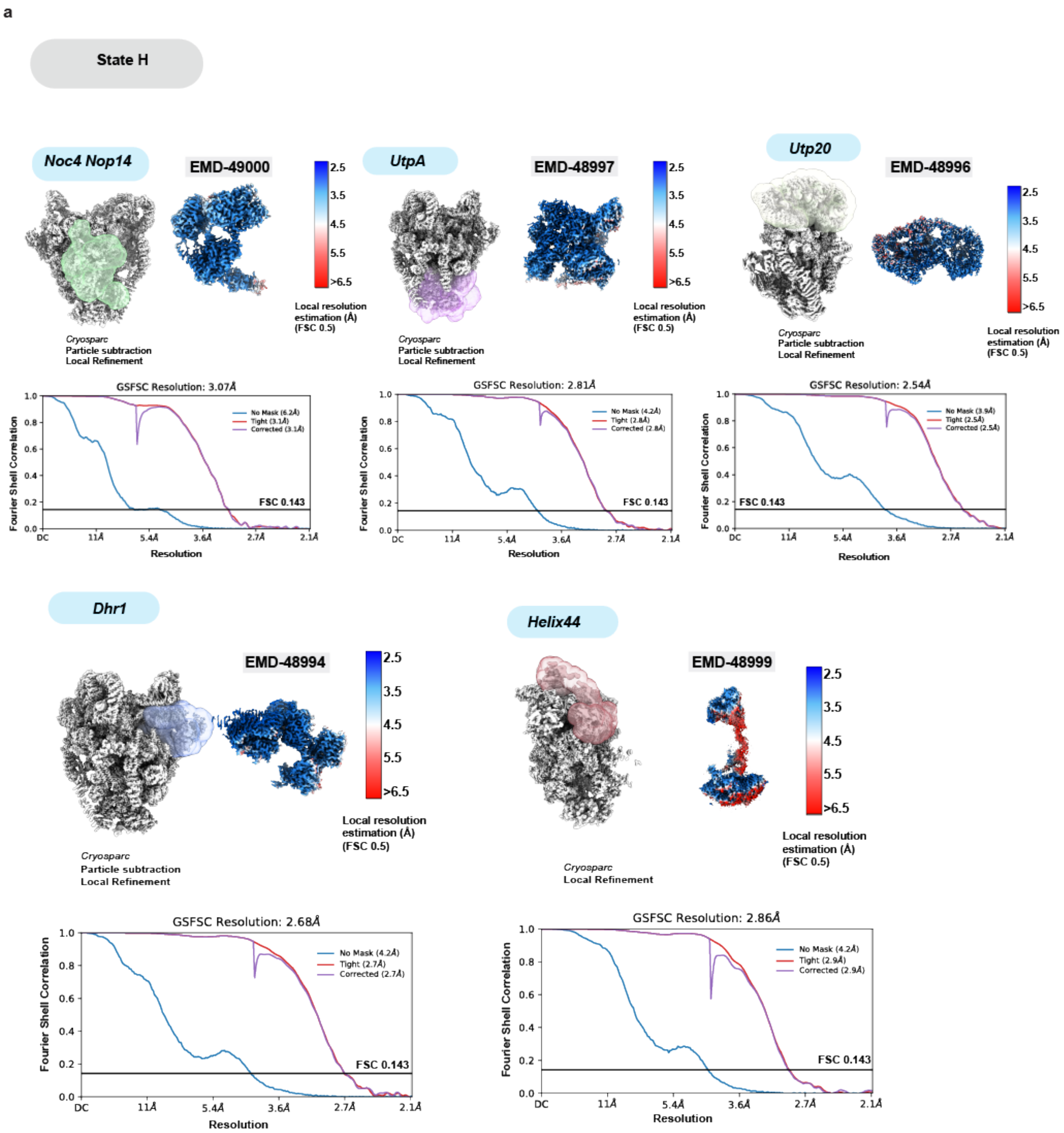


**Supplementary Fig. 17. Cryo-EM focused maps of State H continue.**
(**a**) Focused maps for the Noc4Nop14, UtpA, Utp20, Dhr1, and Helix44 modules used to generate the composite map are displayed along with the respective FSC curves (no mask, tight mask and solvent corrected) on the bottom.

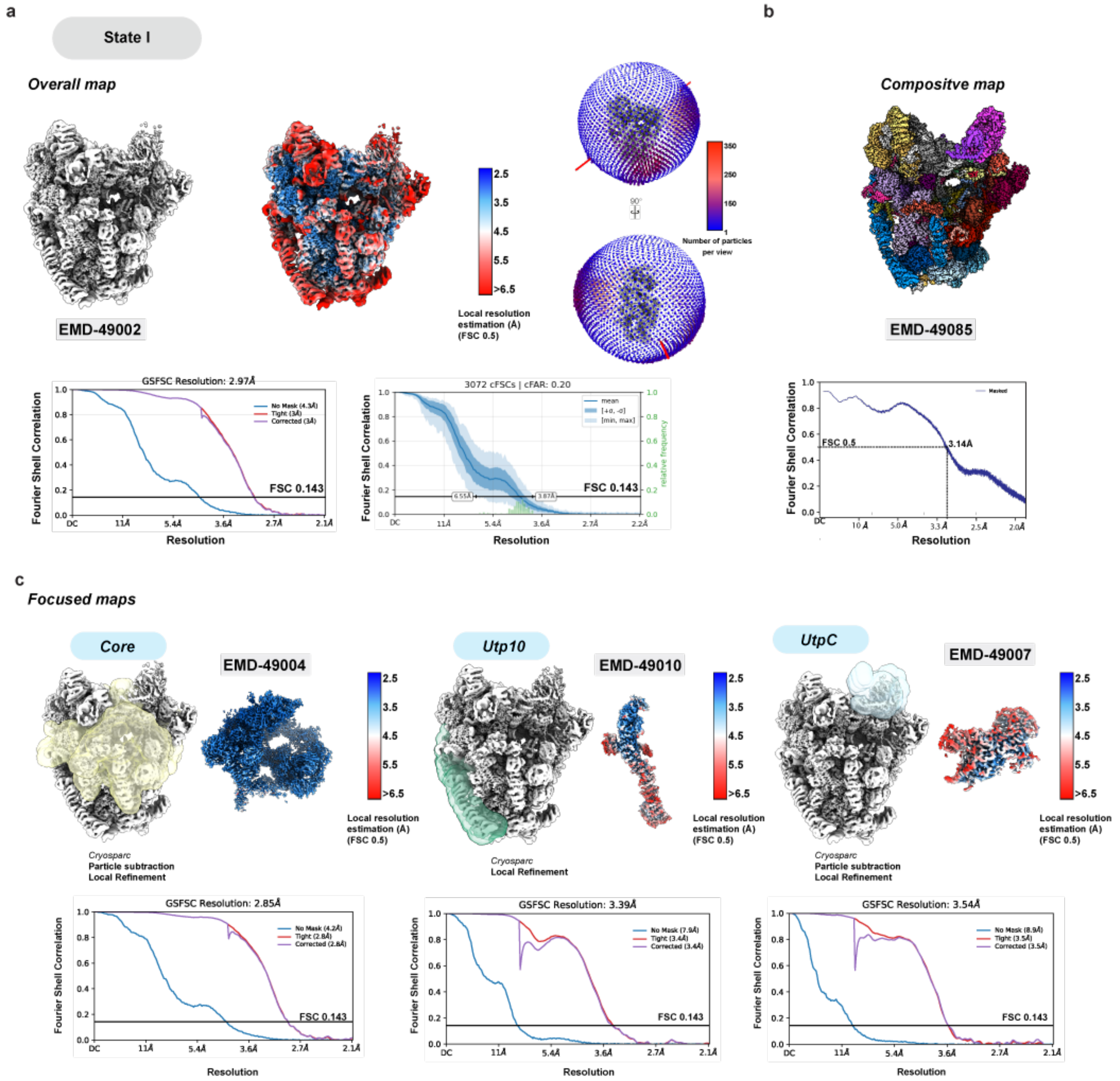


**Supplementary Fig. 18. Cryo-EM focused maps and composite reconstruction of State I.** (**a**) Local-resolution filtered overall map of State I. FSC curves (no mask, tight mask and solvent corrected and 3D) are displayed on the bottom and Euler angle distribution is displayed on the right. (**b**) Composite map and FSC map-to-model curve displayed on the bottom. (**c**) Focused maps for the core, Utp10 and UtpC modules used to generate the composite map are displayed along with the respective FSC curves (no mask, tight mask and solvent corrected) on the bottom.

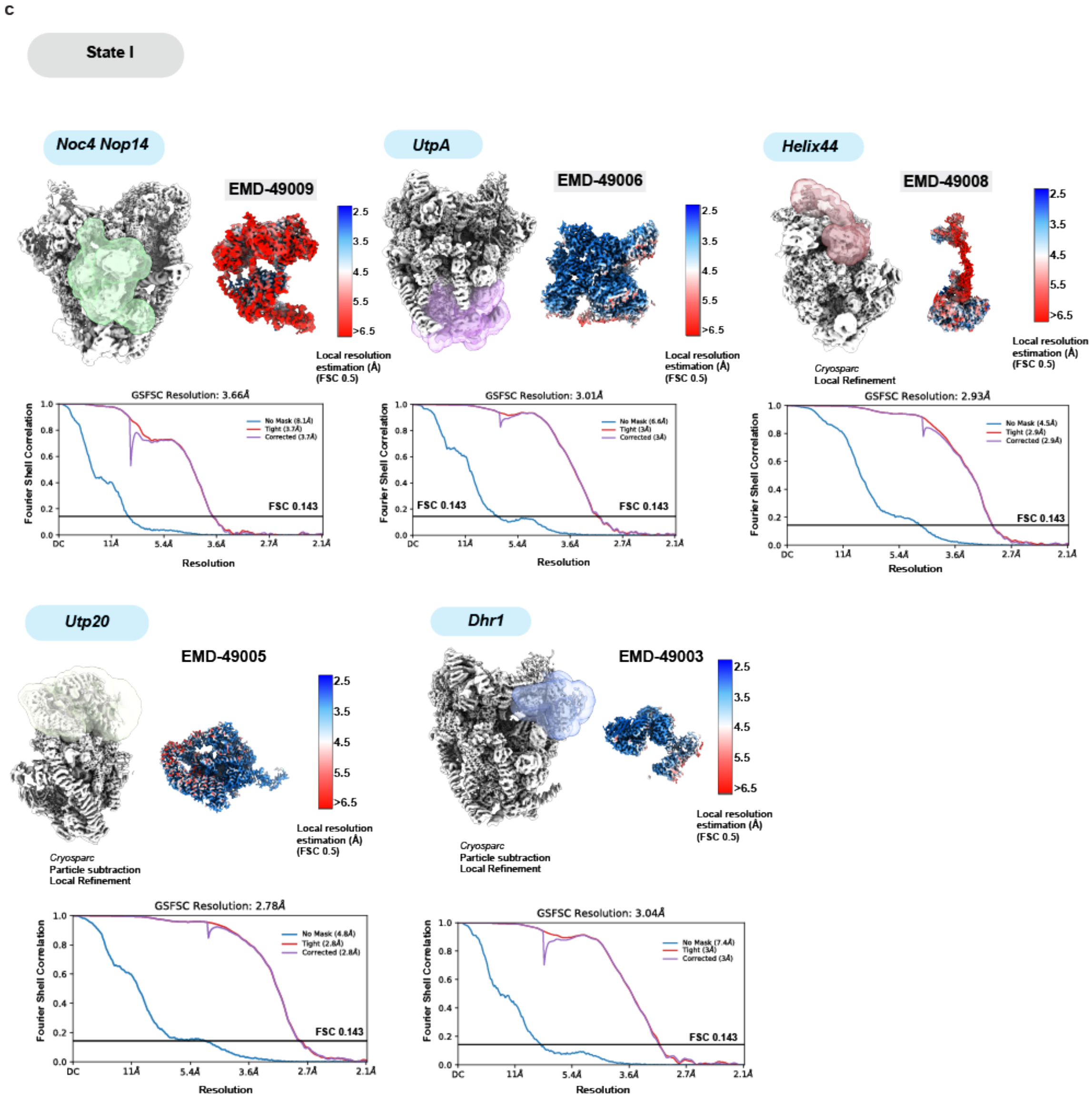


**Supplementary Fig. 19. Cryo-EM focused maps of State I continued.**
(**a**) Focused maps for the Noc4Nop14, UtpA, Helix44, Utp20, and Dhr1 modules used to generate the composite map are displayed along with the respective FSC curves (no mask, tight mask and solvent corrected) on the bottom.

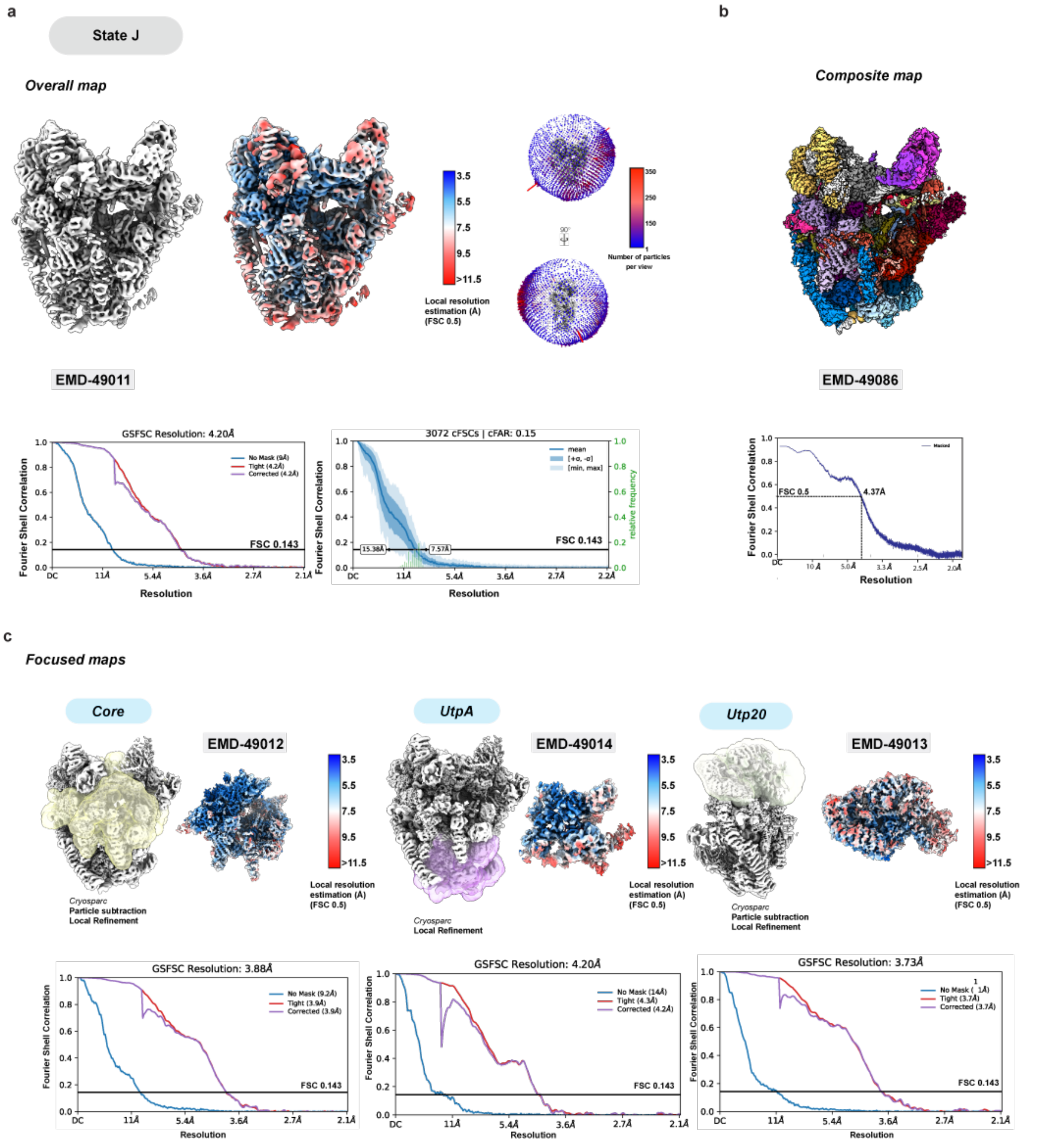


**Supplementary Fig. 20. Cryo-EM focused maps and composite reconstruction of State J.** (**a**) Local-resolution filtered overall map of State J. FSC curves (no mask, tight mask and solvent corrected and 3D) are displayed on the bottom and Euler angle distribution is displayed on the right. (**b**) Composite map and FSC map-to-model curve displayed on the bottom. (**c**) Focused maps for the core, UtpA and Utp20 modules used to generate the composite map are displayed along with the respective FSC curves (no mask, tight mask and solvent corrected) on the bottom.

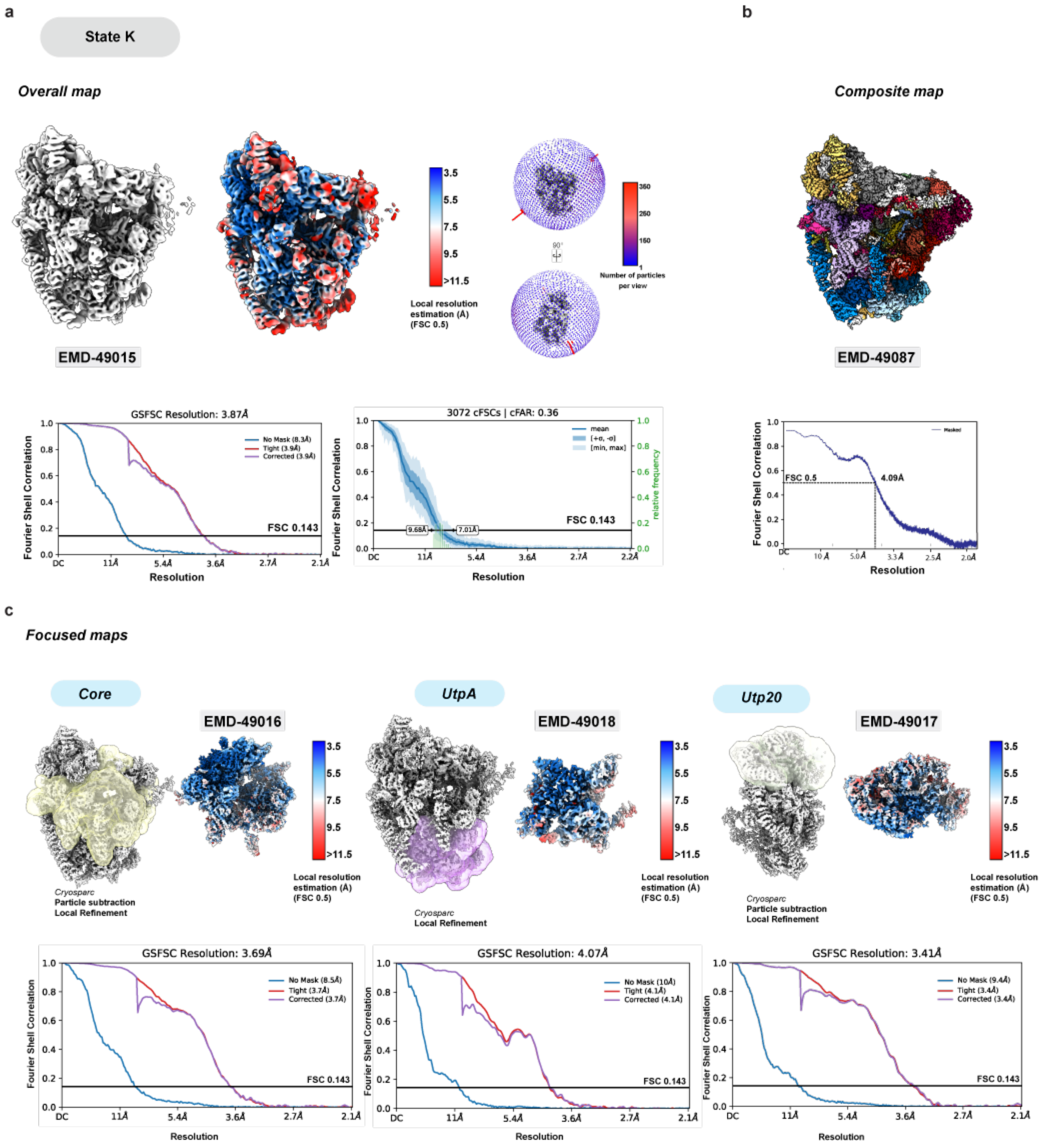


**Supplementary Fig. 21. Cryo-EM focused maps and composite reconstruction of State K.** (**a**) Local-resolution filtered overall map of State K. FSC curves (no mask, tight mask and solvent corrected and 3D) are displayed on the bottom and Euler angle distribution is displayed on the right. (**b**) Composite map and FSC map-to-model curve displayed on the bottom. (**c**) Focused maps for the core, UtpA, and Utp20 modules used to generate the composite map are displayed along with the respective FSC curves (no mask, tight mask and solvent corrected) on the bottom.

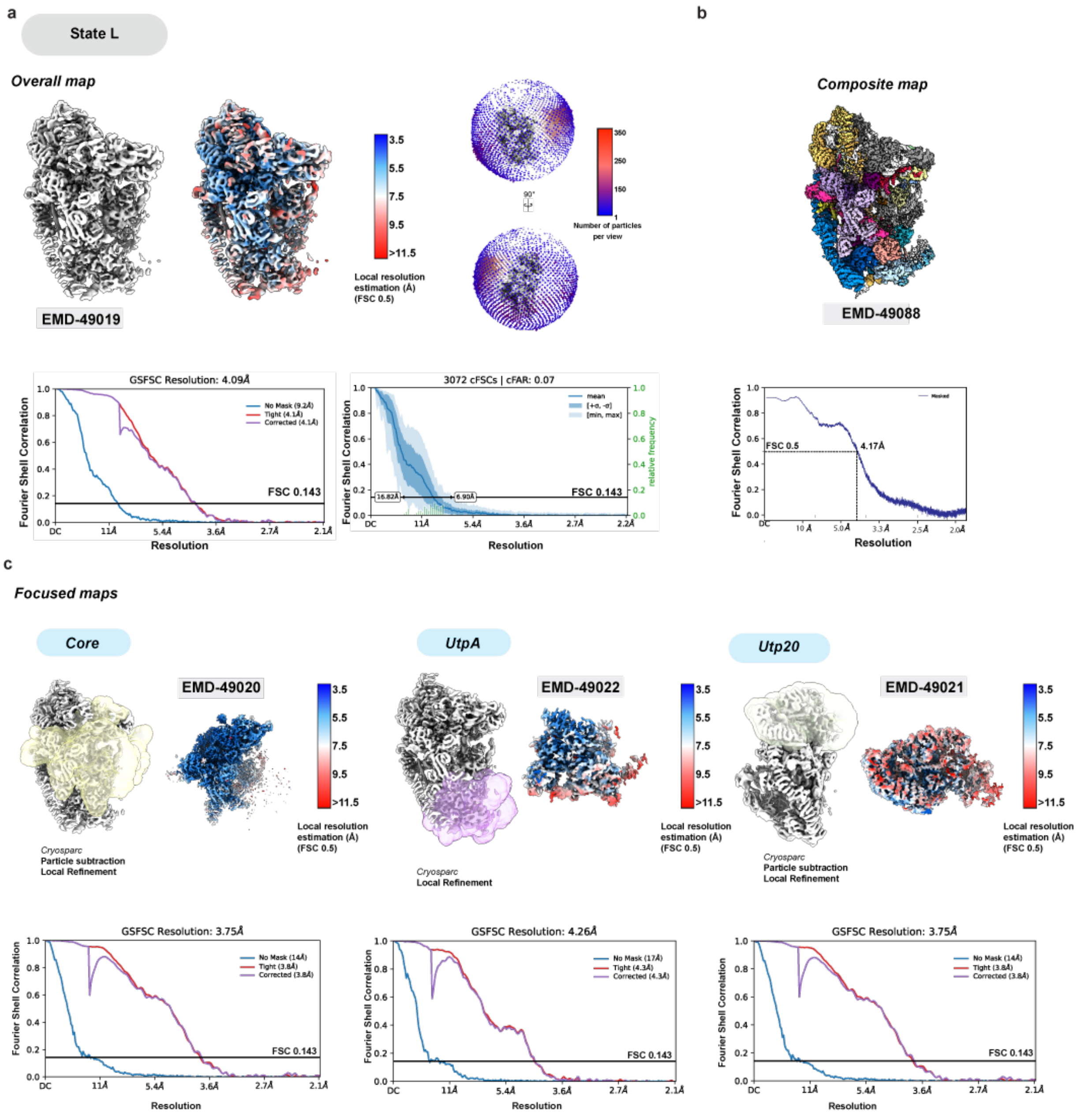


**Supplementary Fig. 22. Cryo-EM focused maps and composite reconstruction of State L.** (**a**) Local-resolution filtered overall map of State L. FSC curves (no mask, tight mask and solvent corrected and 3D) are displayed on the bottom and Euler angle distribution is displayed on the right. (**b**) Composite map and FSC map-to-model curve displayed on the bottom. (**c**) Focused maps for the core, UtpA, and Utp20 modules used to generate the composite map are displayed along with the respective FSC curves (no mask, tight mask and solvent corrected) on the bottom.

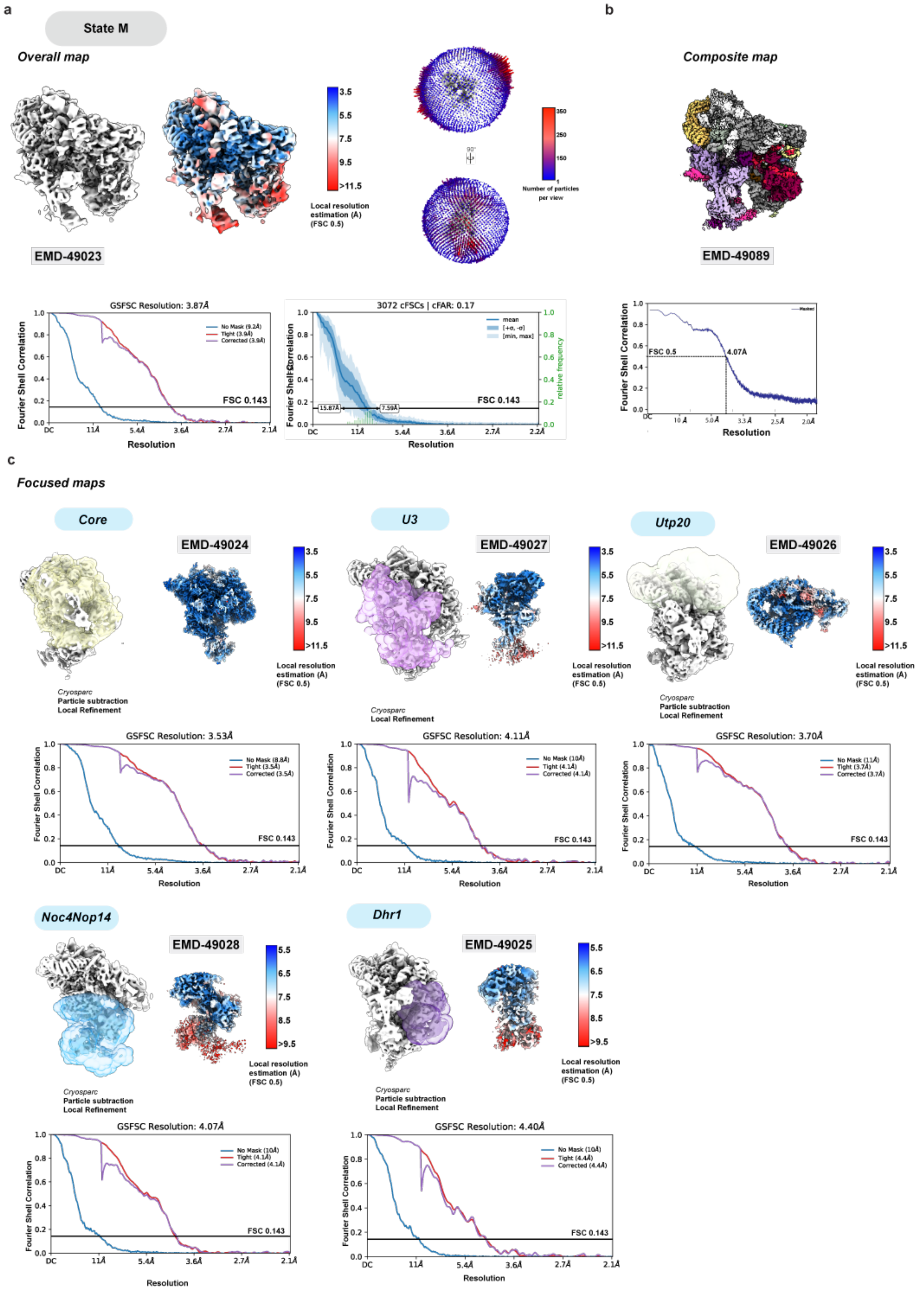
a
State M
Overall map
EMD-49023
Local resolution estimation (Å) (FSC 0.5)
Number of particles per view
GSFSC Resolution: 3.87Å
3072 cFSCs | cFAR: 0.17
FSC 0.143
Fourier Shell Correlation
Resolution
b
Composite map
EMD-49089
FSC 0.5
4.07Å
c
Focused maps
Core
EMD-49024
Cryosparc
Particle subtraction
Local Refinement
U3
EMD-49027
Cryosparc
Local Refinement
Utp20
EMD-49026
GSFSC Resolution: 3.53Å
GSFSC Resolution: 4.11Å
GSFSC Resolution: 3.70Å
Noc4Nop14
EMD-49028
Dhr1
EMD-49025
GSFSC Resolution: 4.07Å
GSFSC Resolution: 4.40Å

**Supplementary Fig. 23. Cryo-EM focused maps and composite reconstruction of State M.** (**a**) Local-resolution filtered overall map of State M. FSC curves (no mask, tight mask and solvent corrected and 3D) are displayed on the bottom and Euler angle distribution is displayed on the right. (**b**) Composite map and FSC map-to-model curve displayed on the bottom. (**c**) Focused maps for the core, U3, Utp20, Noc4nop14, and Dhr1 modules used to generate the composite map are displayed along with the respective FSC curves (no mask, tight mask and solvent corrected) on the bottom.

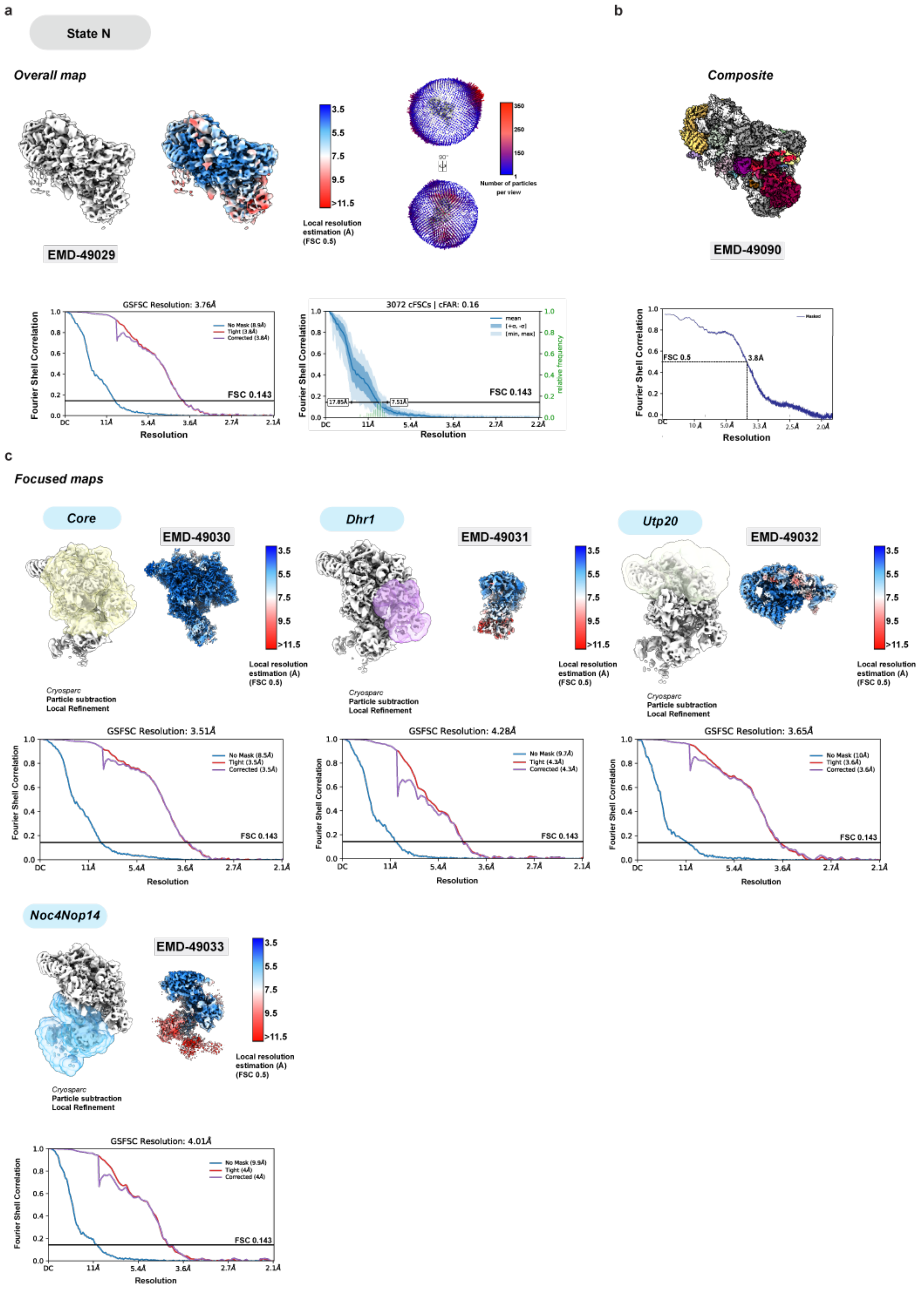
a
State N
Overall map
EMD-49029
3.5
5.5
7.5
9.5
>11.5
Local resolution estimation (Å) (FSC 0.5)
Number of particles per view
GSFSC Resolution: 3.76Å
No Mask (8.9Å)
Tight (3.8Å)
Corrected (3.8Å)
FSC 0.143
Fourier Shell Correlation
Resolution
3072 cFSCs | cFAR: 0.16
mean
[+σ, -σ]
[min, max]
17.85Å
7.51Å
relative frequency
b
Composite
EMD-49090
Masked
FSC 0.5
3.8Å
c
Focused maps
Core
EMD-49030
Cryosparc
Particle subtraction
Local Refinement
GSFSC Resolution: 3.51Å
No Mask (8.5Å)
Tight (3.5Å)
Corrected (3.5Å)
Dhr1
EMD-49031
GSFSC Resolution: 4.28Å
No Mask (9.7Å)
Tight (4.3Å)
Corrected (4.3Å)
Utp20
EMD-49032
GSFSC Resolution: 3.65Å
No Mask (10Å)
Tight (3.6Å)
Corrected (3.6Å)
Noc4Nop14
EMD-49033
GSFSC Resolution: 4.01Å
No Mask (9.9Å)
Tight (4Å)
Corrected (4Å)

**Supplementary Fig. 24. Cryo-EM focused maps and composite reconstruction of State N.** (**a**) Local-resolution filtered overall map of State N. FSC curves (no mask, tight mask and solvent corrected and 3D) are displayed on the bottom and Euler angle distribution is displayed on the right. (**b**) Composite map and FSC map-to-model curve displayed on the bottom. (**c**) Focused maps for the core, Dhr1, Utp20, and Noc4nop14 modules used to generate the composite map are displayed along with the respective FSC curves (no mask, tight mask and solvent corrected) on the bottom.

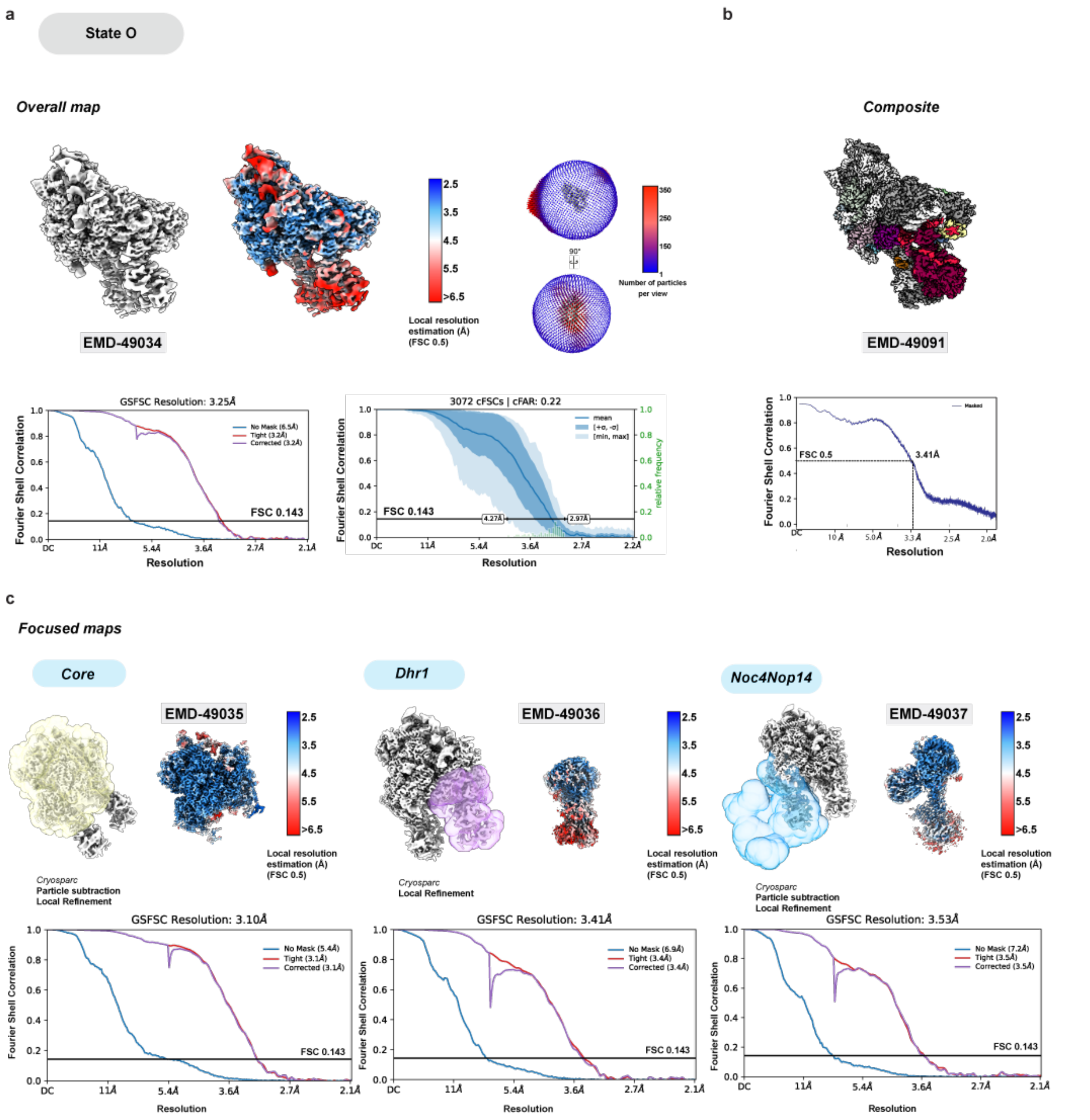


**Supplementary Fig. 25. Cryo-EM focused maps and composite reconstruction of State O.** (**a**) Local-resolution filtered overall map of State O. FSC curves (no mask, tight mask and solvent corrected and 3D) are displayed on the bottom and Euler angle distribution is displayed on the right. (**b**) Composite map and FSC map-to-model curve displayed on the bottom. (**c**) Focused maps for the core, Dhr1, and Noc4nop14 modules used to generate the composite map are displayed along with the respective FSC curves (no mask, tight mask and solvent corrected) on the bottom.

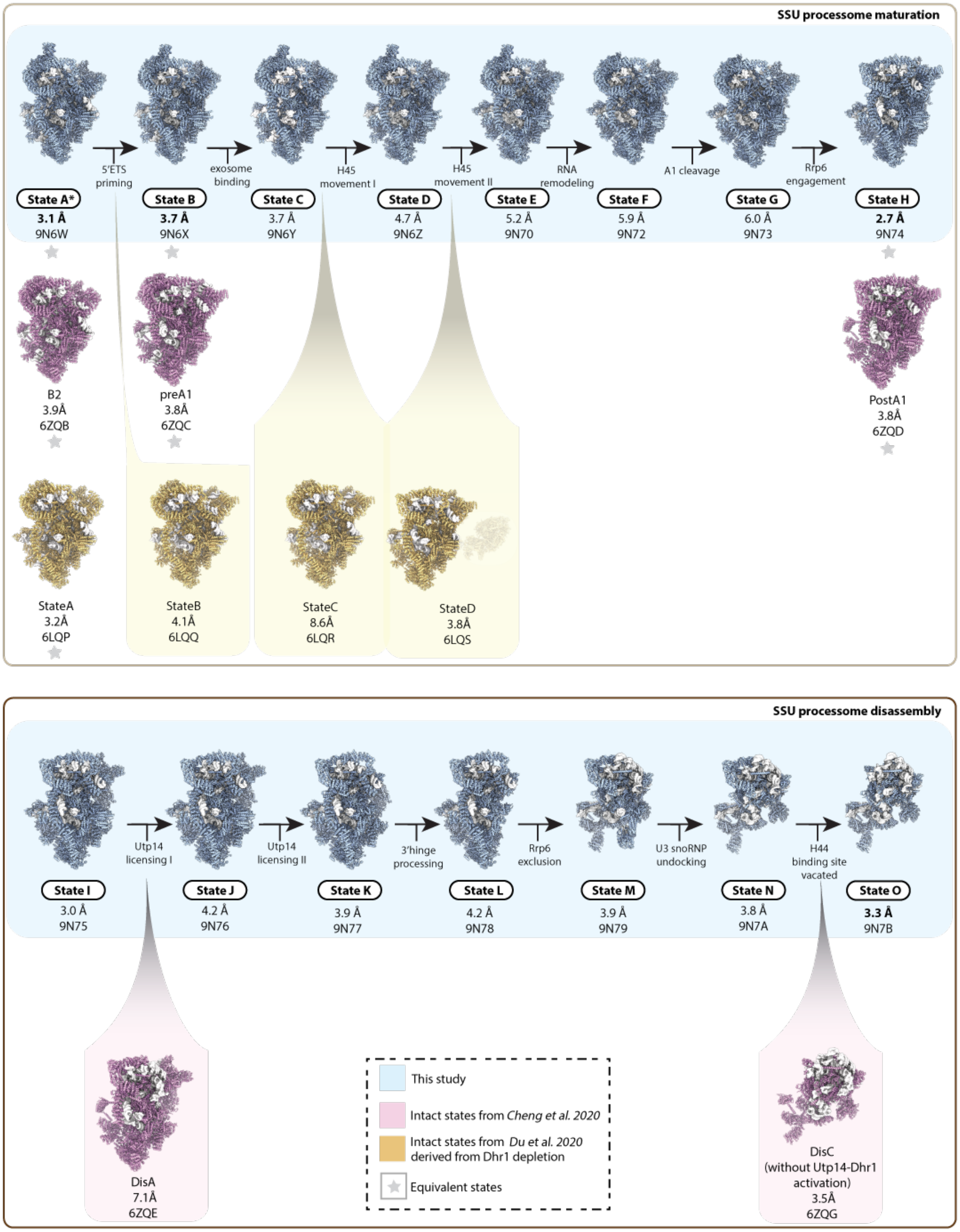
SSU processome maturation
State A*
3.1 Å
9N6W
5'ETS priming
State B
3.7 Å
9N6X
exosome binding
State C
3.7 Å
9N6Y
H45 movement I
State D
4.7 Å
9N6Z
H45 movement II
State E
5.2 Å
9N70
RNA remodeling
State F
5.9 Å
9N72
A1 cleavage
State G
6.0 Å
9N73
Rrp6 engagement
State H
2.7 Å
9N74
B2
3.9Å
6ZQB
preA1
3.8Å
6ZQC
PostA1
3.8Å
6ZQD
StateA
3.2Å
6LQP
StateB
4.1Å
6LQQ
StateC
8.6Å
6LQR
StateD
3.8Å
6LQS
SSU processome disassembly
State I
3.0 Å
9N75
Utp14 licensing I
State J
4.2 Å
9N76
Utp14 licensing II
State K
3.9 Å
9N77
3'hinge processing
State L
4.2 Å
9N78
Rrp6 exclusion
State M
3.9 Å
9N79
U3 snoRNP undocking
State N
3.8 Å
9N7A
H44 binding site vacated
State O
3.3 Å
9N7B
DisA
7.1Å
6ZQE
This study
Intact states from Cheng et al. 2020
Intact states from Du et al. 2020 derived from Dhr1 depletion
Equivalent states
DisC
(without Utp14-Dhr1 activation)
3.5Å
6ZQG

**Supplementary Fig. 26. Comparative view of yeast SSU maturation and disassembly pathways.**

Schematic representation of the *Saccharomyces cerevisiae* SSU processome maturation (top) and disassembly (bottom). Only intact states are depicted, states from this study are highlighted with blue background, intact states from other studies are highlighted with yellow and pink backgrounds and similar states are indicated with a grey star. States from other studies that follow between those observed in this study are indicated with a zoom out. Poor density for the RNA exosome in State D (6LQS) relative to the rest of the map is depicted with shading. Resolutions and PDB codes are indicated.

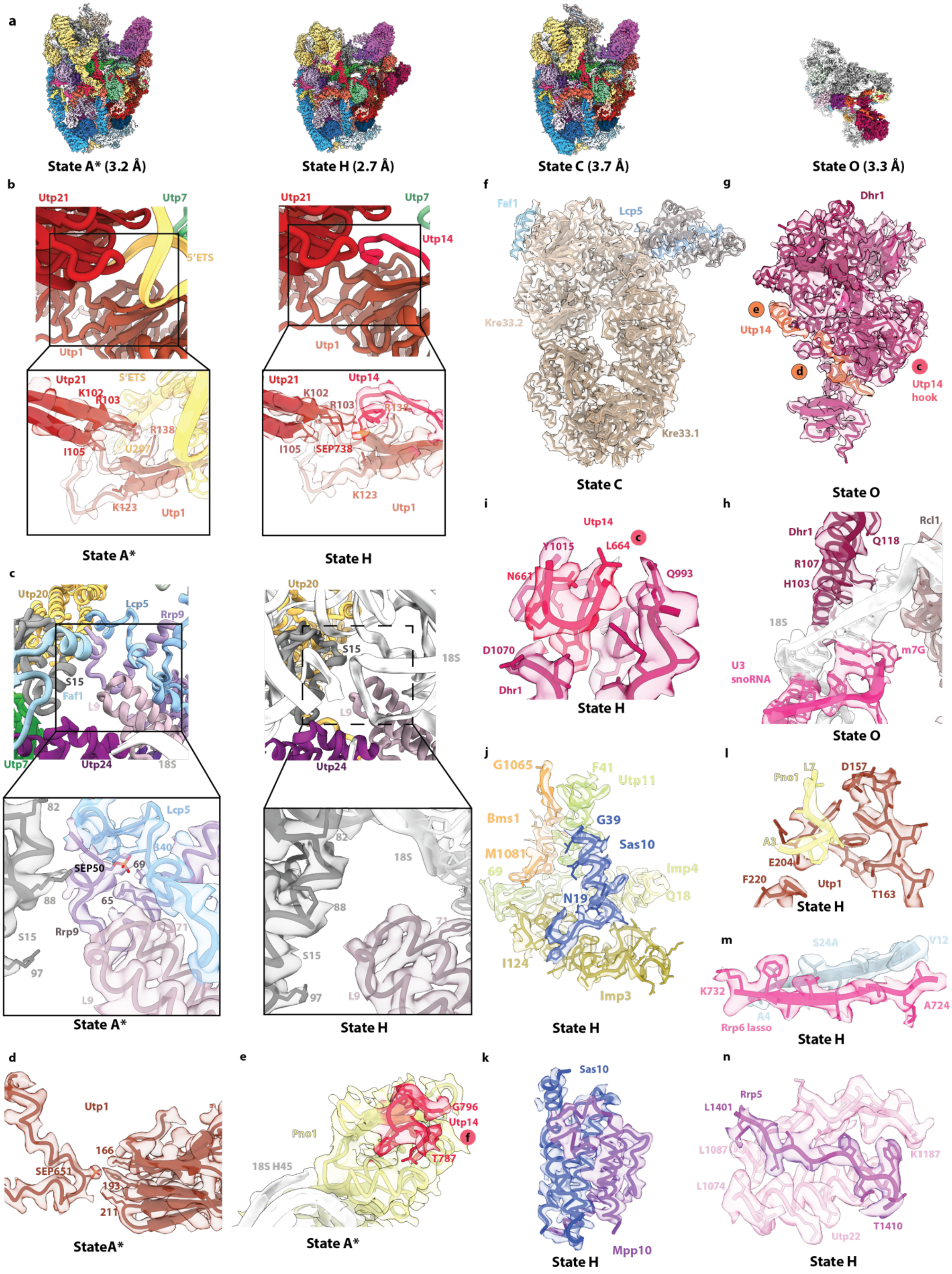
a
State A* (3.2 Å)
State H (2.7 Å)
State C (3.7 Å)
State O (3.3 Å)
b
Utp21
Utp7
5'ETS
Utp1
K102
R103
R138
I105
U297
K123
State A*
Utp14
SEP738
State H
c
Utp20
Lcp5
Rrp9
S15
Faf1
L9
Utp7
Utp24
18S
SEP50
State A*
State H
d
Utp1
166
SEP651
193
211
StateA*
e
Pno1
G796
18S H45
T787
State A*
f
Faf1
Lcp5
Kre33.2
Kre33.1
State C
g
Dhr1
Utp14 hook
State O
h
Rcl1
Q118
R107
H103
18S
U3 snoRNA
m7G
State O
i
Y1015
L664
N661
Q993
D1070
State H
j
G1065
F41
Utp11
Bms1
G39
Sas10
M1081
Imp4
N19
Q18
I124
Imp3
State H
k
Mpp10
State H
l
Pno1
L7
D157
A3
E204
F220
T163
State H
m
S24A
V12
K732
A4
A724
Rrp6 lasso
State H
n
Rrp5
L1401
L1087
K1187
L1074
T1410
Utp22
State H

**Supplementary Fig. 27. Representative Cryo-EM densities and models.**
(**a**) Representative composite cryo-EM maps for States A*, State H, State C and State O with resolutions indicated in parenthesis below each state. (**b-n**) Selection of representative densities for assembly factors and RNAs from States A*, State H, State C and State O. The states from which density and models are derived are shown below each panel. Densities illustrated as continuous transparent volumes and models are shown as ribbons and sticks.

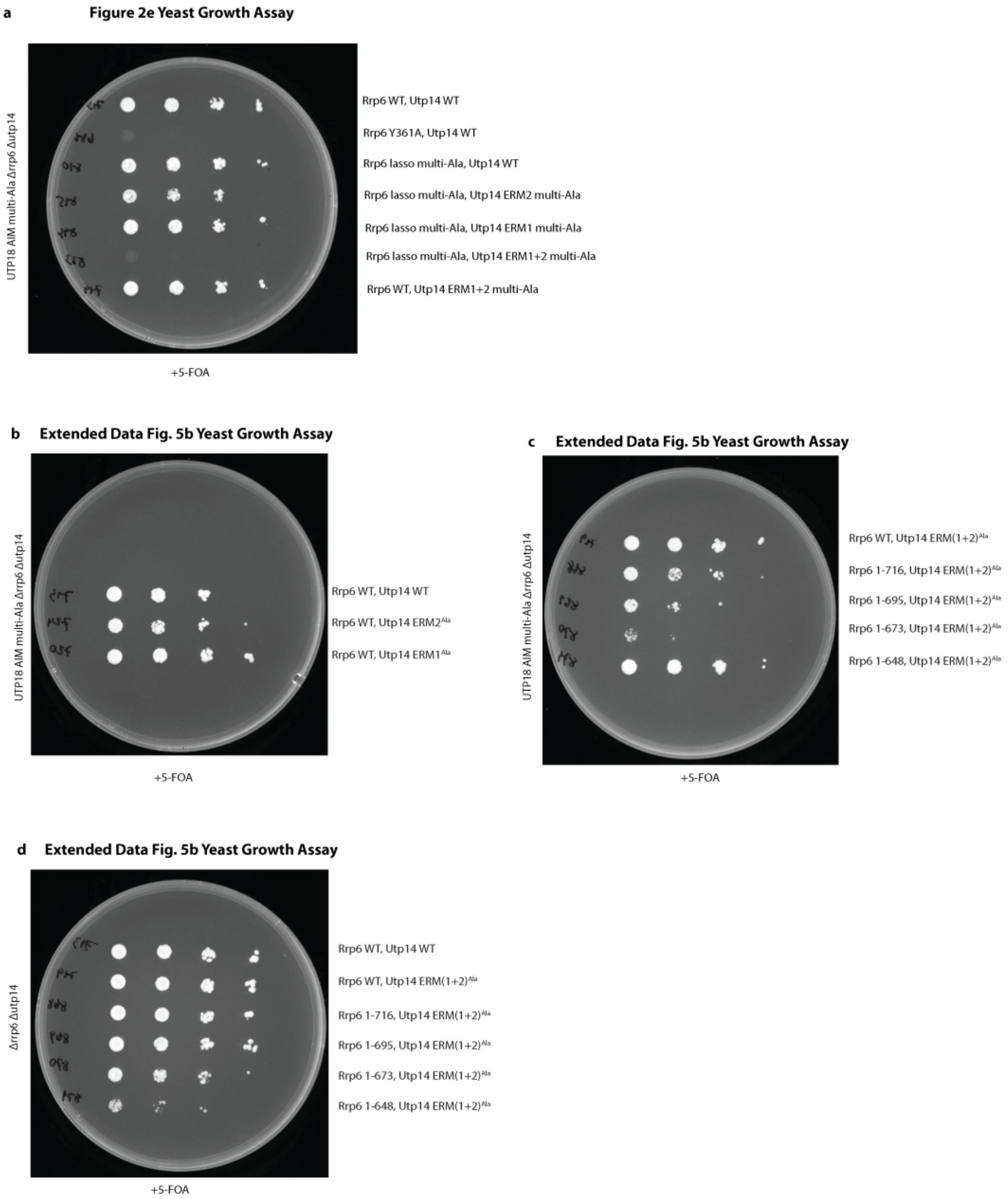


**Supplementary Fig. 28. Uncropped yeast growth assays.**
Uncropped yeast growth assay plates relating to: (**a**) Fig. 2e, (**b-d**) Extended Data Fig.5b
